# Multiscale spatial transcriptomics resolves the cellular and molecular architecture of the human amygdala

**DOI:** 10.64898/2026.08.22.746381

**Authors:** Michael S. Totty, Svitlana V. Bach, Madeline R. Valentine, Madhavi Tippani, Sarah E. Maguire, Ishbel Del Rosario Alvia, Ryan A. Miller, Joel E. Kleinman, Kristen R. Maynard, Stephanie Cerceo Page, Thomas M. Hyde, Stephanie C. Hicks, Keri Martinowich

## Abstract

The amygdala is a central hub for emotional learning that is dysregulated across numerous psychiatric disorders, yet its molecular architecture in humans remains poorly defined. We here present a spatial transcriptomic atlas of the human amygdala from nine neurotypical donors. Integrating Visium, Xenium, and VisiumHD technologies, we profiled over 1.1 million cells across 13 spatial domains spanning the basolateral complex, central, medial, and cortical nuclei, as well as the intercalated islands, each defined by distinct marker genes and gene co-expression networks. By integrating snRNA-seq reference atlases, we found that each subdivision of the basolateral subnucleus contains distinct excitatory neuron classes. We additionally found three transcriptionally distinct populations of intercalated neuron types, two of which form spatially distinct islands neighboring the basolateral complex, and were able to refine cell type diversity across the central nucleus and related amygdalostriatal transition areas. Finally, we localized psychiatric genetic risk across the amygdala, revealing both broad neuronal enrichment and subnuclear specificity. Together, this atlas provides a foundational resource and framework for accelerating cross-species comparisons and human disease-focused investigations.

## 1 Introduction

The amygdala is a central hub for emotional learning and affective processing that integrates sensory inputs with internal state signals to shape adaptive and maladaptive behavioral responses ^1–3^. Dysregulation of amygdala circuits is commonly implicated in neuropsychiatric conditions including post-traumatic stress disorder ^4,5^, major depressive disorder ^6,7^, and substance use disorders ^8,9^. Despite its central role in brain function and disease, the cellular and molecular architecture of the human amygdala remains incompletely understood. Much of our current understanding of amygdala organization originates from rodent studies ^10–13^ or classical primate and human neuroanatomical work that lack comprehensive molecular characterization ^14–17^. Consequently, it remains unclear how molecular identity, cellular composition, and spatial organization collectively define the functional specialization of the human amygdala. Resolving this cellular and molecular architecture is essential for identifying cell type-specific mechanisms of psychiatric vulnerability and for guiding the development of more precise, circuit-informed therapeutic strategies. In humans, the amygdala is a large complex structure composed of multiple anatomically and functionally distinct subnuclei with complex cell type organization, including the basolateral amygdalar (BLA) complex, central (CeA) and medial nuclei (MeA), the cortical nucleus (CoA), and the intercalated amygdalar islands (IA). Decades of classical neuroanatomical work have defined major subdivisions of the BLA complex, including the lateral (LA), basolateral (BL), paralaminar (PL), and basomedial nuclei (BM), based on cytoarchitecture, connectivity, and immunohistochemical features ^18^. More recently, single-nucleus RNA sequencing (snRNA-seq) studies have identified transcriptionally distinct neuronal populations enriched within specific subnuclei, substantially expanding our understanding of amygdala cell type diversity ^19–23^. However, because these approaches lack spatial information, they cannot comprehensively resolve all subnuclei, precisely localize transcriptionally defined populations, or determine how distinct neuronal populations are organized within individual subnuclei. This limitation is particularly consequential for emerging snRNA-seq studies of neuropsychiatric disorders, which identify disease-associated transcriptional changes, but cannot assign them to specific amygdala subnuclei ^24,25^. Consequently, how transcriptionally defined cell populations map onto classical neuroanatomical subnuclei, and how disease-associated molecular changes are distributed across these subnuclei, remain largely unresolved.

Here, we developed a multiscale spatial transcriptomic framework to resolve the cellular and molecular architecture of the human amygdala by integrating complementary transcriptome-wide and single cell spatial profiling technologies. Transcriptome-wide Visium profiling was used to reconstruct the molecular organization of the entire amygdala and identify subnucleus-specific gene programs, Xenium *in situ* profiling provided single-cell resolution mapping of transcriptionally-defined neuronal populations, and VisiumHD resolved the fine-scale cellular architecture of specialized, spatially compact nuclei including the IA and CeA. Applying this framework, we resolved the molecular identity of magnocellular and parvocellular BL nucleus, defined three spatially distinct IA cell types, localized several putative CeA populations to adjacent striatal transition zones, and assessed the spatial enrichment of heritability for various complex traits and psychiatric disorders. Together, this multimodal framework defines the spatial organization of molecular programs across the human amygdala, links transcriptional identity to classical anatomical domain organization, and provides a foundation for cross-species studies and the spatial interpretation of human transcriptomic datasets.

## 2 Results

### 2.1 Multiscale spatial transcriptomic profiling of the human amygdala

To comprehensively characterize the molecular organization of the human amygdala, we combined three complementary spatially-resolved transcriptomic (SRT) platforms. Visium provides transcriptome-wide profiling at 55 µm resolution over 6.5 mm^2^ areas, Xenium provides single cell resolution with a limited gene panel across the full extent of the amygdala, and VisiumHD combines transcriptome-wide profiling with single cell resolution, but like standard Visium, is limited to small 6.5 mm^2^ capture areas. Together, these platforms combine transcriptome-wide profiling across the full extent of the amygdala with targeted single cell-resolution molecular characterization of specific subnuclei (**Fig. 1**). Tissue blocks from nine neurotypical control brain donors (5 male/4 female) were dissected from coronal hemislabs of fresh frozen postmortem human tissue at the midpoint of the amygdala on the anterior-posterior (A-P) axis, a location where all major amygdala subnuclei are present (**Fig. 1A-C**). Inclusion of the amygdala within each tissue block was confirmed using anatomical landmarks visible in both the frozen tissue block and H&E stained sections (**SFig 1-9**).

**Fig. 1.**
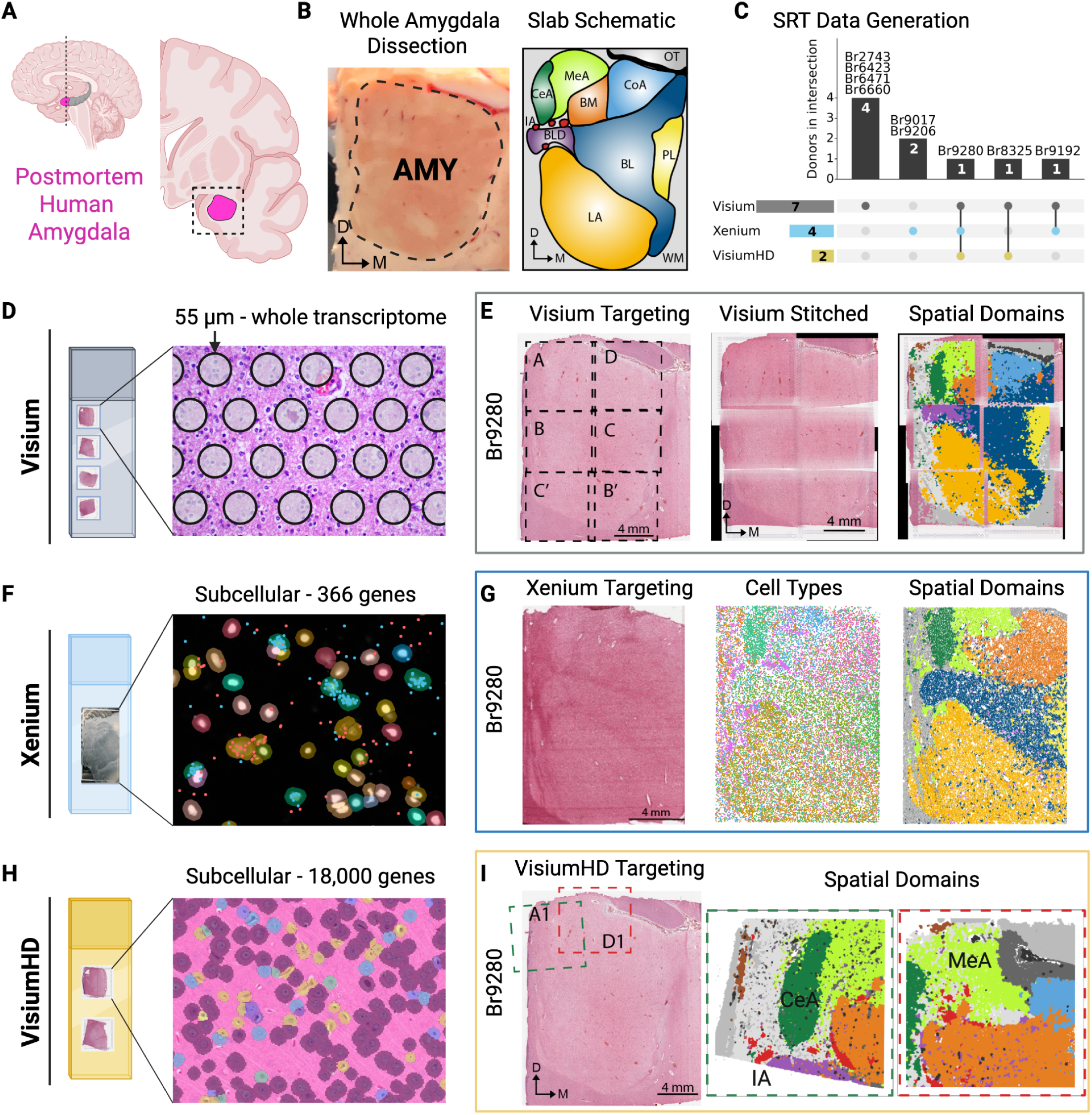
Schematic of postmortem human brain tissue collection and spatial transcriptomics data generation. **(A)** Schematic representation of the human amygdala (pink) and hippocampus (gray) in midsagittal (left) and coronal (right) views. The level of the coronal section is represented by the dashed line on the midsagittal view. The dashed box denotes the boundaries of the dissected brain block. **(B)** Dissected coronal human brain block containing the outlined amygdala (AMY, left) and a schematic of amygdala subnuclei for the corresponding A-P axis (right). Basolateral amygdala (BL), basolateral dorsal amygdala (BLD), basomedial amygdala (BM), central amygdala (CeA), cortical amygdala (CoA), dorsal (D), intercalated amygdalar islands (IA), lateral amygdala (LA), medial (M), medial amygdala (MeA), optic tract (OT), paralaminar amygdala (PL), white matter (WM). **(C)** UpSet plot depicting donors used for the three SRT data generation applications: Visium 7 donors (Br2743, Br6423, Br6471, Br6660, Br8325, Br9280, Br9192); Xenium 4 donors (Br9280, Br9192, Br9206, Br9017); VisiumHD 2 donors (Br8325, Br9280). **(D)** Schematic representation of the Visium slide with four 6.5 mm^2^ capture arrays, each containing ∼5,000 barcoded spots 55 µm in diameter to capture whole-transcriptome of multiple cells in a spot. **(E)** Visium data generation workflow for one representative donor, Br9280. Left to right: AMY tissue is scored in six 6.5 mm^2^ strips to match the width of the Visium capture arrays (Visium Targeting). After the Visium assay, H&E and SRT data are reassembled to represent the whole tissue (Visium Stitched) and Spatial Domains are defined. **(F)** Schematic representation of the Xenium slide with one large capture array containing probes for 366 genes, allowing for subcellular resolution. **(G)** Xenium data generation workflow for one representative donor, Br9280. Left to right: unscored AMY tissue is placed on the Xenium array (Xenium Targeting), subcellular resolution allows to determine cell types and spatial domains within the sample. **(H)** Schematic representation of the VisiumHD slide with two 6.5 mm^2^ capture arrays, each containing a continuous lawn of 2 µm barcoded sequences, allowing for subcellular resolution of ∼18,000 genes. **(I)** Visium HD data generation workflow for one representative donor, Br9280. Left to right: AMY tissue is scored in two 6.5 mm^2^ strips to match the width of the Visium capture arrays (Visium Targeting). After the Visium HD assay, detailed spatial domains are determined for selected AMY subnuclei: IA, CeA, and MeA. Created in BioRender.

To generate transcriptome-wide molecular maps spanning the entire amygdala, we performed Visium Spatial Gene Expression profiling from seven of the nine donors (**Fig. 1D-E**). For each donor, cryosections were tiled across 6–8 capture areas spanning two slides per donor (*n*=51 total capture areas), which were computationally aligned and stitched to construct a continuous transcriptome-wide map of the intact amygdala ^26^. The final Visium dataset comprised 224,021 high-quality spots following preprocessing and quality control (**SFig 10**).

To complement the transcriptome-wide Visium atlas with higher resolution molecular profiling, we performed both Xenium *in situ* profiling and VisiumHD Spatial Gene Expression on selected amygdala subnuclei (**Fig. 1F-I**). Xenium was performed in four donors, including two from the Visium cohort (Br9192 and Br9280, 2 males) (**SFig 1,2**) and two additional donors (Br9017 and Br9206, 2 females) using a custom 366-gene panel derived from Visium-defined spatial domain markers and our previously published human amygdala snRNA-seq dataset ^19^. The final Xenium dataset comprised 946,896 high-quality cell segmentations following quality control (**SFig 11**). To obtain high-resolution, transcriptome-wide characterization of molecularly specialized nuclei, we generated five VisiumHD samples across two donors (Br9280 and Br8325, 1 male/1 female, **SFig 2 and 5**), targeting the CeA (*n*=2) and MeA (*n*=2) nuclei, as well as the intercalated IA (*n*=1) cell masses. The final VisiumHD dataset comprised 281,929 high-quality cell segmentations following quality control (**SFig 12**).

**Fig. 2.**
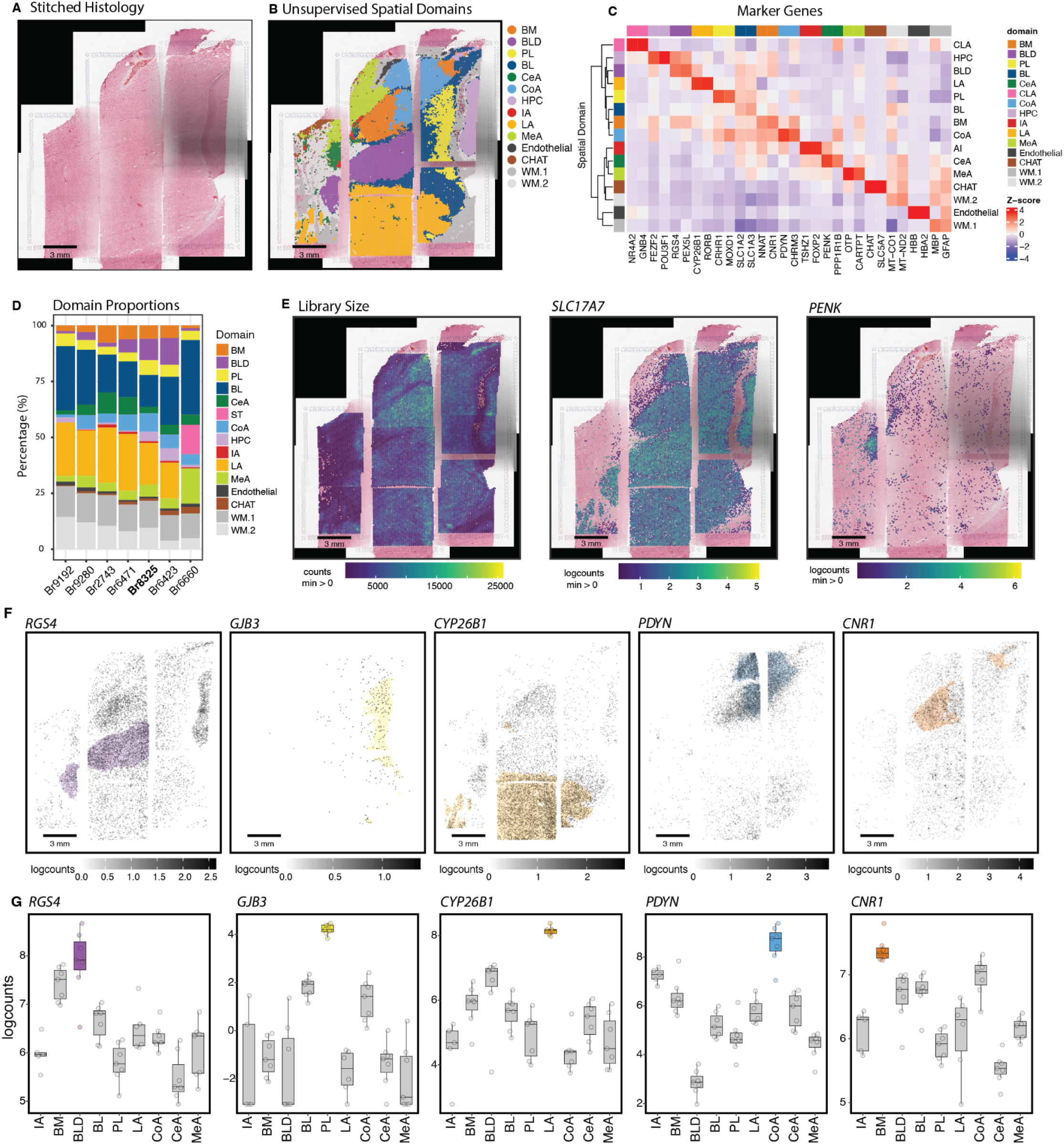
Whole-transcriptome spatial profiling reveals novel molecular markers across amygdala nuclei. **(A)** Stitched histology (H&E) of coronal human amygdala Visium sections from a representative donor. Scale bar, 3 mm. **(B)** Unsupervised spatial domain assignments for the sections in (A), delineating amygdala nuclei (LA, BL, BM, PL, CoA, CeA, MeA, IA, BLD) alongside adjacent hippocampus (HPC), claustrum (CLA), white matter (WM.1, WM.2), endothelium (Endo), and cholinergic-enriched (CHAT) domains. **(C)** Scaled expression (z-score) of representative marker genes across spatial domains. Rows are ordered by hierarchical clustering of pseudobulked gene expression profiles across domains. **(D)** Percentage of spots assigned to each spatial domain, per donor (*n*=7), showing consistent domain recovery across tissue sections. Donors are ordered from the most anterior (left) to most posterior (right). **(E)** Spatial distribution of library size (total UMI counts) and of *SLC17A7* and *PENK* (log-normalized counts) for the donor in (A). *SLC17A7* marks excitatory neurons and is broadly enriched across the basolateral complex, whereas *PENK* is restricted to the central nucleus (CeA). **(F)** Spatial expression (log-normalized counts) of *RGS4*, *GJB3*, *CYP26B1*, *PDYN*, and *CNR1*, marking the BLD, PL, LA, BL/BM, and CoA, respectively. Each gene is scaled independently; scale bars, 3 mm. **(G)** Domain-level expression of the genes in (F) across donors (log-normalized counts). Boxes show median and interquartile range across donors, with points indicating individual donor means; the domain of maximal enrichment is colored.

### 2.2 Whole-transcriptome spatial profiling resolves the molecular organization of the human amygdala

To define the molecular organization of the human amygdala in an unbiased manner, we applied unsupervised spatial clustering to the stitched whole-amygdala Visium samples (**Fig. 2A-B**). To account for batch effects introduced by tiling across multiple Visium capture areas, we implemented an iterative, reference-guided clustering strategy with additional cell type mapping to accurately identify the IA islands (**SFig. 13**; Methods). This approach resolved spatially distinct molecular domains corresponding to the major amygdala nuclei, including the lateral (LA), basolateral (BL), basolateral dorsal (BLD), paralaminar (PL), basomedial (BM), central (CeA), cortical (CoA), medial (MeA), and intercalated (IA) (**Fig. 2A-B**). Domain annotations were supported by established marker genes including *RORB* (LA), *CRHR1* (PL), *NNAT* (BM), *TSZH1* and *FOXP2* (IA), *PENK* (CeA), and *OTP* (MeA), while hierarchical clustering of domain-averaged gene expression recapitulated known molecular relationships among amygdala nuclei, including transcriptional similarity among nuclei of the BLA complex (LA, BLD, BL, PL and BM) (**Fig. 2C**).

We next examined the anterior-posterior distribution of these molecular domains by classifying the seven whole amygdala sections from each of the seven donors into one of four anatomical levels: anterior-intermediate (*n*=1, Br9192), intermediate (*n*=3, Br9280, Br2743, Br6471), posterior-intermediate (*n*=2, Br8325, Br6423), and posterior (*n*=1, Br6660). Molecular domains were consistently identified across the seven donors, with relative abundance being largely consistent, and differences along the A-P axis explained by established human amygdala neuroanatomy (**Fig. 2D**) ^27^. The LA was most prominent at anterior levels and absent posteriorly, whereas the BM, BLD, CeA, and CoA were most abundant at intermediate levels (**Fig. 2D**). Visualization of library size and representative marker genes further confirmed anatomical fidelity of the reconstructed atlas (**Fig. 2E**). *SLC17A7*, encoding vesicular glutamate transporter 1, was enriched throughout the glutamatergic BLA complex, whereas *PENK* showed complementary enrichment within the predominantly GABAergic CeA (**Fig. 2E**).

Although previously established marker genes supported our annotations of several domains, relatively few markers have been reported that distinguish individual subnuclei of the human BLA complex. We therefore leveraged our spatially resolved dataset to identify the top marker genes distinguishing individual subnuclei. One-vs-all pseudobulk differential expression (**SFig. 14**) analysis identified *RGS4* as a prominent marker for the BLD, *GJB3* for the PL, *CYP26B1* for the LA, *PDYN* for CoA, and *CNR1* for the BM (**Fig. 1F-G**). A full list of the top 100 marker genes for each domain is provided in **Supplementary Data 2**.

### 2.3 Spatial co-expression networks define domain-specific molecular programs in the human amygdala

While differential expression analysis identified specific genes distinguishing individual amygdala domains, coordinated gene expression programs can provide complementary insight into the molecular networks underlying domain identity and function. We therefore performed spatial co-expression network analysis across the seven Visium donors. We identified 30 co-expression modules containing at least 10 genes, of which 10 modules exhibited strong domain-specific spatial enrichment (**Fig. 3A-C**). These included modules corresponding to the LA (M6; 287 genes), BLD (M27; 24 genes), PL (M5; 287 genes), CoA (M10; 120 genes), CeA (M13; 99 genes), MeA (M19; 60 genes), and IA (M20; 45 genes), as well as neighboring HPC (M15; 86 genes), CLA (M7; 245 genes), and CHAT domains (M15; 88 genes) .

**Figure 3.**
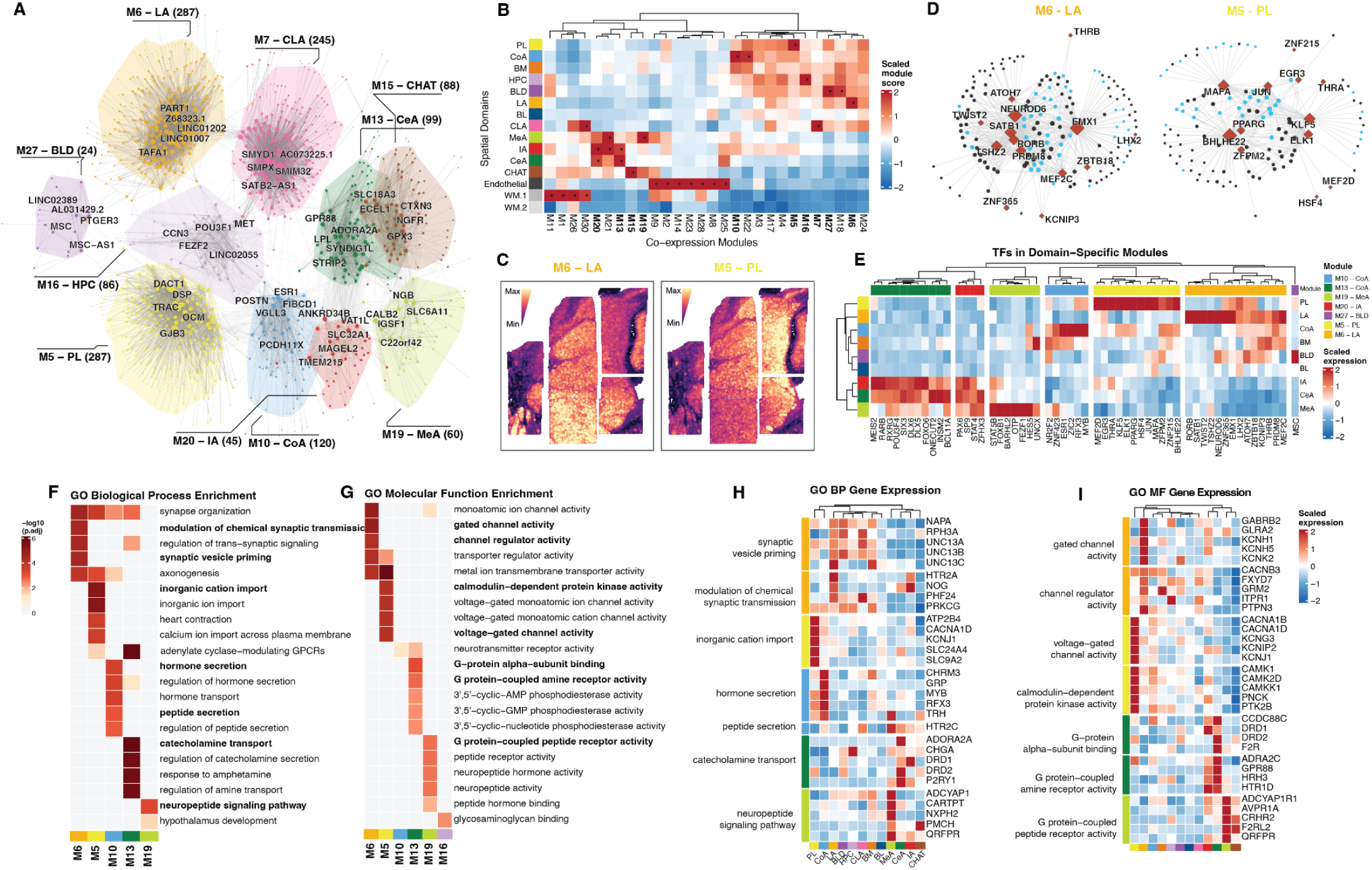
Spatial co-expression network analysis identifies domain-specific gene programs across the human amygdala. **(A)** Spatial co-expression networks for the 10 representative domain-specific modules. Nodes are genes, colored by the spatial domain to which the module is assigned; node size is proportional to within-module degree. The five most highly connected genes per module are labeled, and module labels give the domain and gene count. **(B)** Scaled module scores across spatial domains for all recovered modules. Rows are domains, columns are modules, and both are ordered by hierarchical clustering. Asterisks mark domain–module pairs with z-score > 1.5. **(C)** Spot-level module scores for M6–LA and M5–PL in a representative donor (Br8325), showing spatial enrichment of each program to its assigned domain. **(D)** Transcription factor–centric subnetworks for M6–LA and M5–PL, retaining only edges linking a transcription factor (red diamonds, labeled) to its co-expressed targets. Blue circles marker genes independently identified as domain-enriched by pseudobulk differential expression; grey circles are remaining module members. **(E)** Scaled mean expression (z-score) of module-resident transcription factors across spatial domains. Column annotation indicates each factor’s module of origin. **(F–G)** GO enrichment across the 10 domain-specific modules for biological process **(F)** and molecular function **(G)**, showing up to the top five terms per module ranked by adjusted p-value. Color denotes −log₁₀(adjusted p-value); terms enriched in more than one module are shown once. **(H–I)** Scaled expression (z-score) across spatial domains of top genes from selected GO biological process **(H)** and molecular function **(I)** terms, grouped by term.

To identify candidate transcriptional regulators underlying these domain-specific gene programs, we examined transcription factor (TF) connectivity within each co-expression module (**Fig. 3 D-E**). Using the Lambert et al. (2018) human TF census ^28^, we found that each module contained between 1 and 15 highly connected TFs (**SFig. 15**), several of which corresponded to domain-specific marker genes identified in the previous analysis. These included *NEUROD6*, *EMX1*, *RORB*, and *SATB1* in the LA module, *ESR1* in the CoA module, *OTP* and *FEZF1* in the MeA module, and *MEIS2* in the CeA module (**Fig. 3D**).

To determine the biological functions associated with each domain-specific gene program, we performed gene ontology (GO) enrichment analyses (**Fig. 3F-G**). Distinct functional themes emerged across amygdala domains, consistent with their known specializations. For example, the LA and PL modules were enriched for ion channel-related functions, including gated channel activity (driven by *GABRB2*, *GLRA2*, and *KCNH1*), voltage-gated channel activity (driven by *CACNA1B* and *CACNA1D*), and calmodulin-dependent protein kinase activity (including *CAMK1* and *PNCK*). These enrichments are consistent with the specialized electrophysiological properties of nuclei within the BLA complex ^29^ (**Fig. 3H-I**). In contrast, the CeA module was enriched for G-protein alpha-subunit binding and G protein-coupled amine receptor activity (*DRD1* and *DRD2*), as well as serotonin (*HTR1D*), noradrenaline (*ADRA2C*), and histamine receptors (*HRH3*) (**Fig. 3H-I**). These findings are consistent with the established role of the CeA as an integrative hub for neuromodulatory signaling involved in emotional and behavioral regulation^30,31^ . Together, these spatially resolved co-expressed networks provide a framework for linking the anatomical organization of the human amygdala to coordinated molecular programs and prioritizing candidate genes, regulators, and pathways for future investigation. The complete gene membership and functional enrichment results for each co-expression module are provided in **Supplementary Data 3**.

### 2.4 Single-cell spatial transcriptomics validates domain-specific molecular organization

To validate the molecular organization defined by the transcriptome-wide Visium atlas at single-cell resolution, we performed Xenium *in situ* transcriptomic profiling across coronal sections spanning the whole amygdala in four donors (**Fig 4; SFig 16**). Using unsupervised clustering (**Fig. 4A**), we identified spatiomolecular domains corresponding to the major amygdala nuclei including the lateral (LA), basolateral (BL), and paralaminar (PL) nuclei, basomedial and cortical nuclei (BM-CoA), central nucleus (CeA), medial nucleus and intercalated cell islands (MeA-IA), as well as surrounding structures including the entorhinal cortex (EC), subiculum (SUB), and other cortical structures (Ctx) (**Fig. 4B,C**). We also found domains corresponding to white matter (WM) and a vascular endothelial domain (Endo). Given the targeted 366-gene Xenium panel (**Supplementary Data 4**), closely related or spatially-adjacent regions that could not be reliably separated were annotated as joint domains (e.g., MeA-IA). These molecular domains were consistently identified across the four donors at similar proportions, with the LA and BL comprising the largest proportion of the amygdala.

**Fig. 4:**
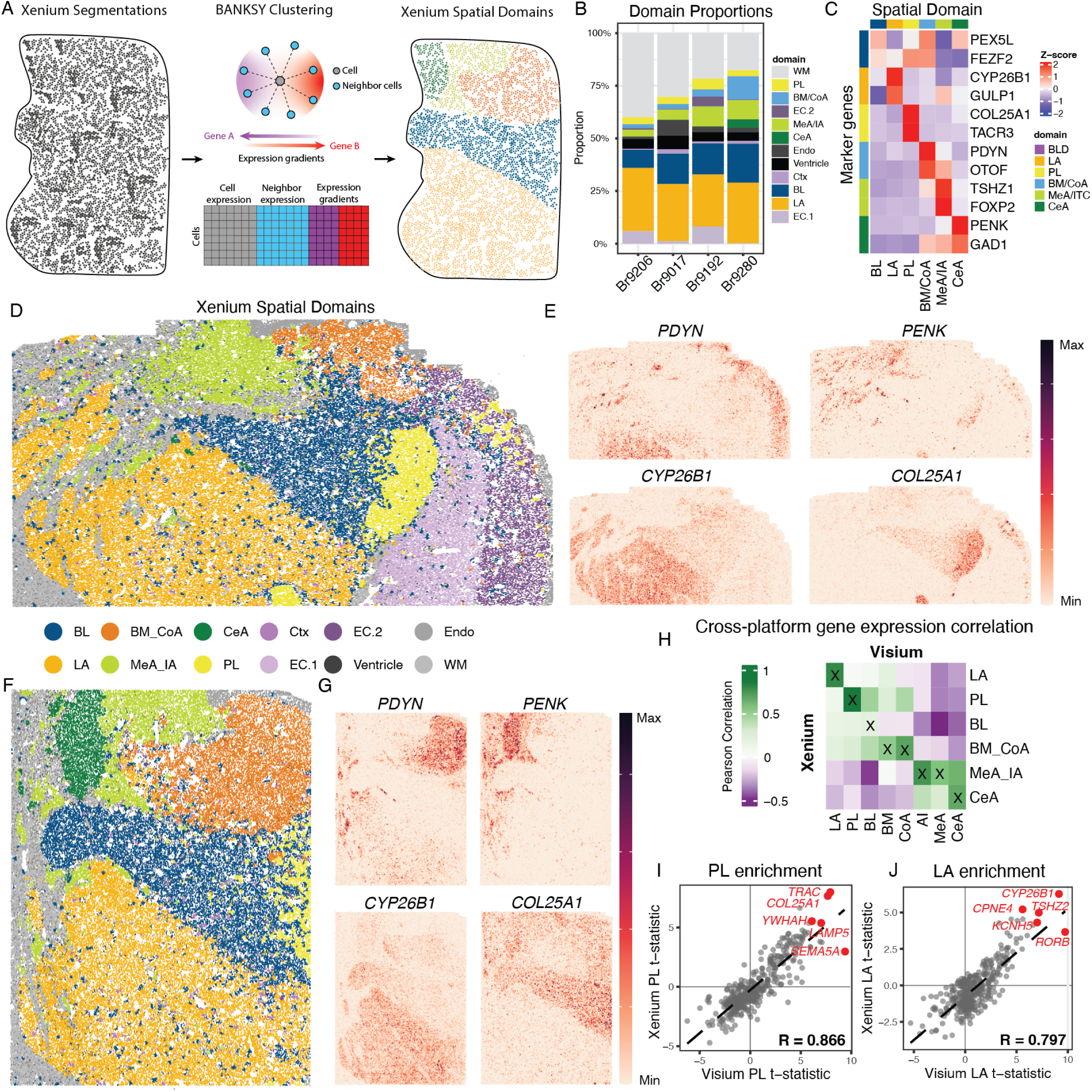
Xenium imaging-based transcriptomics resolves amygdala spatial domains at cellular resolution and validates Visium-derived marker genes. **(A)** Schematic of the Xenium clustering workflow using BANKSY, which performs spatially-aware clustering by combining single-cell gene expression profiles with neighborhood-averaged expression and directional expression gradients to discover smooth, contiguous spatial domains.**(B)** Proportion of cells assigned to each spatial domain, per donor, illustrating domain composition across tissue sections. **(C)** Heatmap of scaled expression (z-score) for selected marker genes across Xenium spatial domains, with genes ordered by domain of maximal enrichment. **(D)** Xenium spatial domain map for a representative donor (Br9206). Each point is a segmented cell colored by BANKSY domain assignment, resolving amygdala subnuclei (LA, BL, BM_CoA, MeA_IA, CeA, PL) alongside adjacent cortex, entorhinal cortex, white matter, ventricle, and endothelium. **(E)** Rasterized (∼25 µm bins) spatial expression of *PDYN*, *PENK*, *CYP26B1*, and *COL25A1* in the donor shown in (D). Expression is scaled independently per gene (min–max); enrichment patterns correspond to the domains identified in Visium. **(F)** As in (D–E) for a second donor (Br9280), showing reproducibility of domain annotation and marker gene localization across tissue sections and individuals. **(G)** Rasterized (∼25 µm bins) spatial expression of *PDYN*, *PENK*, *CYP26B1*, and *COL25A1* in the donor shown in (F). **(H)** Pearson correlation of domain-level gene expression t-statistics between Visium (columns) and Xenium (rows) across matched amygdala domains. Diagonal enrichment indicates cross-platform concordance of domain identity. **(I–J)** Scatter plots of per-gene t-statistics from Visium (x-axis) against Xenium (y-axis) for PL **(I)**, R = 0.866, and LA **(J)**, R = 0.797. Each point is one gene; the top domain-enriched genes are labeled in red.

We next asked whether the domain-specific marker genes identified by Visium exhibited concordant spatial enrichment in the Xenium data. Across Xenium-defined domains, established marker genes showed highly concordant domain-specific expression patterns that replicated both the Visium (**Fig. 4D,F**) and our previous snRNA-seq findings^19^. These included *CYP26B1* and *GULP1* in the LA, *PEX5L* in the BL, *COL25A1* in the PL, *PDYN* in the CoA, and *PENK* in the CeA (**Fig. 4E**). The spatial expression patterns of these genes sharply delineated individual amygdala subnuclei at both the anterior-intermediate and intermediate levels (**Fig. 4G,H**). To quantitatively assess concordance between Visium and Xenium, we computed Pearson correlations of domain-level log-fold changes across all 366 shared genes (**Fig. 4H**). Strong agreement was observed across matched spatial domains particularly for the PL (R=0.867; **Fig. 4I**) and LA (R=0.797; **Fig 4J**), where previously identified marker genes, including *COL25A1* and *CYP26B1*, ranked among the most highly concordant markers across platforms. These findings demonstrate that Xenium robustly recapitulates the molecular organization defined by Visium while extending spatial resolution to the single cell level.

### 2.5 Spatial cell type mapping resolves the molecular identities of neuronal subtypes in amygdala subnuclei

Although classical neuroanatomical studies have long distinguished cytoarchitectural subdivisions of the human amygdala, the molecular identities of the neuronal populations within these regions remains incompletely understood. This is particularly true for the basolateral complex, where parvocellular and magnocellular domains have historically been defined by differences in neuronal morphology ^18^, but their underlying molecular identities currently remain unresolved. To address this question, we integrated our whole amygdala Visium and Xenium datasets with a previously published snRNA-seq atlas of the human amygdala Yu et al ^20^ (**Fig. 5A, B**). Using cell type deconvolution, we mapped reference cell types onto the Visium and Xenium datasets to infer the spatial distribution of transcriptionally-defined neuronal populations across the amygdala (**SFig. 17**; Methods).

**Fig. 5:**
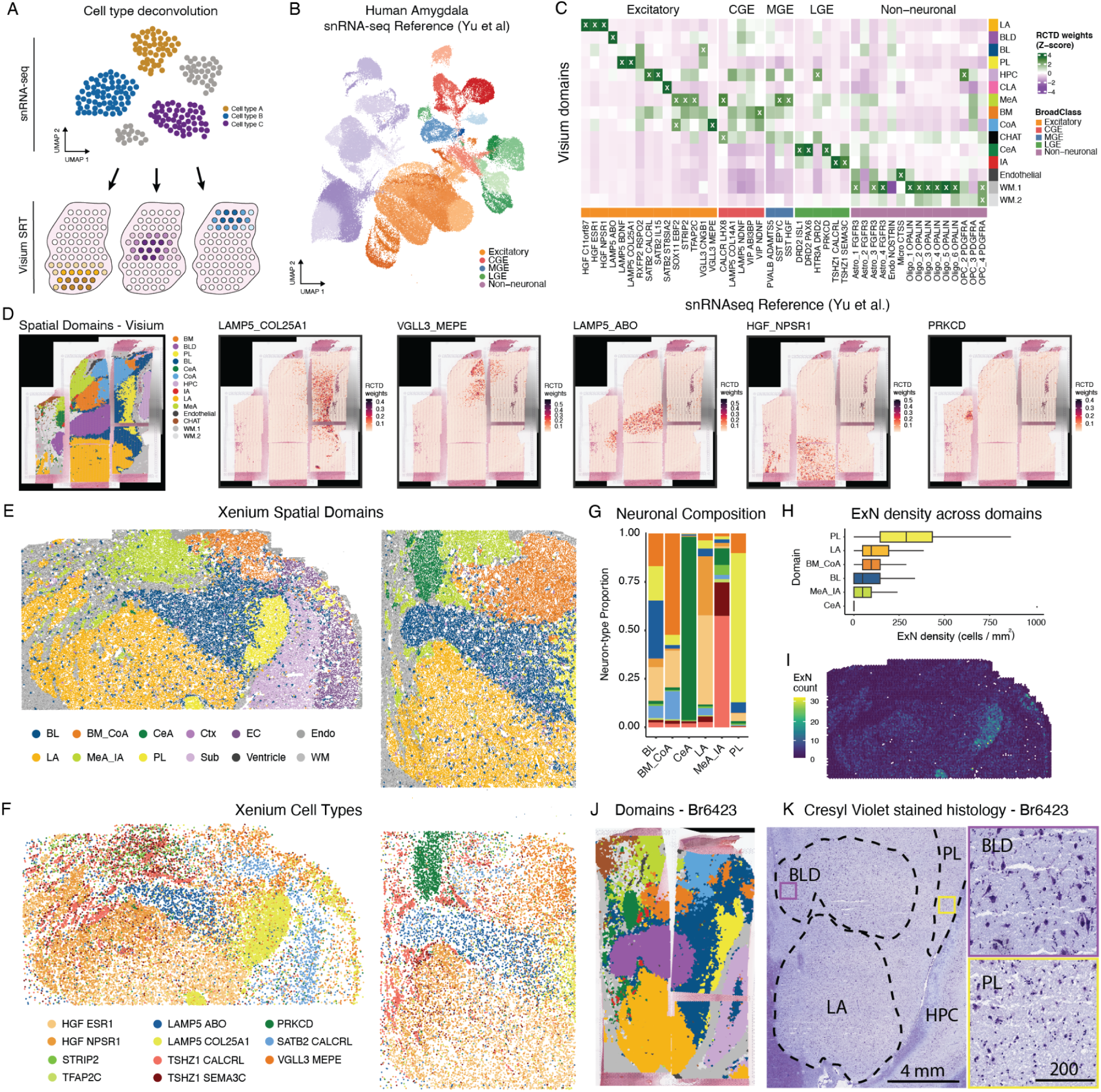
Cross-modality spatial mapping of amygdala cell types reveals the molecular identity of magnocellular and parvocellular projection neurons. **(A)** Schematic of the deconvolution and label-transfer strategy. A published human amygdala snRNA-seq reference (Yu et al.) is used with RCTD to estimate cell type composition per Visium spot and to assign cell types to segmented Xenium cells. **(B)** UMAP of the snRNA-seq reference, colored by broad class: excitatory neurons, inhibitory neurons of caudal (CGE), medial (MGE), and lateral (LGE) ganglionic eminence origin, and non-neuronal cells. **(C)** Enrichment of each reference cell type (columns) across Visium spatial domains (rows), shown as row-wise z-scored RCTD weights. Columns are grouped by broad class; domain–cell type pairs with z-score > 2 are marked with an X. **(D)** Visium spatial domains (left) and RCTD weights for representative cell types (LAMP5_COL25A1, VGLL3_MEPE, LAMP5_ABO, HGF_NPSR1, and PRKCD) in a representative donor (Br8325). Weights are scaled per cell type. **(E)** BANKSY-defined spatial domains in two representative Xenium sections, with each point representing a segmented cell. **(F)** RCTD-assigned cell type labels for the same cells shown in (E), colored to correspond with their domain of predominant occupancy. **(G)** Neuronal composition of each amygdala domain, aggregated across donors (*n* = 4). Bars show the proportion of neuron types assigned to each neuronal subtype. **(H)** Excitatory neuron (ExN) density per hexbin across amygdala domains (cells/mm²). Domains are ranked by median density; boxes show median and interquartile range. **(I)** Spatial distribution of ExN counts per hexbin in a representative Xenium section, showing highest density in the PL. **(J)** Visium spatial domain annotation for donor Br6423, delineating major amygdala nuclei and surrounding structures. **(K)** Cresyl violet–stained section from the same donor at low (left) and high (right) magnification, contrasting the smaller, densely packed parvocellular neurons of the PL with the larger magnocellular neurons of the BLD. Yellow and purple boxes on the low-magnification image indicate the regions shown at high magnification for PL and BLD, respectively. Scale bars: 4 mm (low), 200 µm (high).

Mapping transcriptionally-defined cell types onto the Visium spatial domains revealed striking subnuclei specificity (**Fig. 5C-D**). The resulting enrichment patterns largely replicated our previous findings from anatomically-targeted snRNA-seq ^19^, while providing comprehensive anatomical context across the entire amygdala. Excitatory neuron subtypes exhibited particularly strong subnuclear specificity, with *VGLL3+/MEPE+* neurons mapping to the CoA, *LAMP5+/BDNF+* and *LAMP5+/COL25A1+* neurons mapping to the PL, *LAMP5+ABO+* neurons mapping to the BLD, all the three *HGF+* neuronal populations mapping to the LA, and *STRIP2+* and *TFAP2C+* populations mapping the MeA. Of note, the previously described *SOX11+/EBF2+* population, originally proposed to represent a PL cell type because of its immature neuronal gene expression profile^20^, instead mapped predominantly to the MeA and CoA. This finding is consistent with our previous observation that this population most closely resembles *SLC17A6*+ MeA neurons ^19^. Additional excitatory neuronal populations also exhibited highly selective spatial localization, including two *SATB2*+ populations within the HPC, and a *SATB2+/ST8SIA2+* population that mapped almost exclusively to the claustrum (CLA). Likewise, three of the four putative CeA neuronal populations localized to the CeA domain, whereas the two *TSHZ1+* populations mapped specifically to the IA domain. Among non-neuronal cell types most mapped preferentially to the WM.1, except for the Micro_CTSS population, which mapped to the Endothelial domain and the OPC_2 *PDGFRA+* class, which mapped to the HPC. For many amygdala-specific neuronal populations enrichment was largely restricted to a single subnucleus (**Fig. 5C**), demonstrating a close correspondence between transcriptional identity and classical subnuclei annotation. We additionally used cell type deconvolution with our previously published, anatomically targeted amygdala snRNA-seq dataset [19]. This analysis replicated the major domain-specific enrichment patterns that we previously found (**SFig. 18**).

To validate these findings at single cell resolution, we next examined the distribution of transcriptionally-defined neuronal populations in the Xenium dataset (**Fig. 5E,F; SFig. 19**). Spatial mapping recapitulated the subnuclear specificity observed by Visium deconvolution, with *VGLL3+/MEPE3+* neurons again localizing to the CoA and BM, *LAMP5+/COL25A1+* neurons to the PL, *LAMP5+/ABO+* neurons to the BL/BLD, and *HGF+* neurons to the LA. We also confirmed that *PRKCD+* neurons, but not the two *DRD2+* populations, mapped to the CeA, whereas *TSHZ1+*, *STRIP2+*, and *TFAP2C+* populations localized to the adjacent IA and MeA domains (**Fig. 5E-F**) . Quantification of neuronal composition across spatial domains further highlighted marked differences in cellular architecture (**Fig. 5G**). The PL was composed almost exclusively of *LAMP5*+/*COL25A1*+ neurons, while the more dorsal region of the BL was instead dominated by *LAMP5*+ / *ABO*+ neurons. Consistent with the classical description of the PL as a densely packed parvocellular division ^18^, excitatory neuron density was markedly higher in the PL compared to other subnuclei (**Fig. 5H-I**). Comparison with adjacent Nissl-stained sections demonstrated that this molecular boundary closely recapitulates the well-established cytoarchitectonic transition from the densely packed parvocellular PL to the larger, magnocellular neurons of the dorsal BL. (**Fig. 5J,K; SFig. 20**). These findings establish the molecular identities of the classical parvocellular and magnocellular divisions of the human basolateral amygdala, linking long-recognized cytoarchitectural organization with transcriptionally-defined neuronal populations.

### 2.6 Integrative analysis links molecularly distinct intercalated cell subtypes to distinct spatial organizations in the human amygdala

We have thus far used “IA” to denote the anatomically defined intercalated islands of the amygdala, consistent with the human brain atlas. Here, as we turn from spatial domains to defined cell types, we choose to use “ITC”, the terminology more commonly used to refer to the neuron types that make up IA islands. While we know that ITCs are small clusters of densely packed GABAergic neurons that regulate information flow between the BLA and central amygdala, their molecular diversity and spatial organization in the human brain remains poorly understood. To identify reproducible human ITC subtypes, we integrated and compared three independently generated human amygdala snRNA-seq datasets using MetaNeighbor to assess cross-dataset correspondence (**Fig. 6A**)^19,20,22^. This identified three highly reproducible ITC subtypes (ITC_1, ITC_2, and ITC_3) that were consistently recovered across all three datasets (**Fig. 6B; SFig. 21**). Although the relative abundance of these subtypes differed across datasets (**Fig. 6C**), all three populations were well represented in the Siletti et al., dataset, which was therefore used as the reference cell type comparisons and spatial mapping (**Fig. 6D**). All three subtypes expressed high levels of transcription factors critical to ITC development, including *TSHZ1* and *FOXP2* **(Fig. 6E)**, as well as the canonical marker genes *DRD1* and *OPRM1* (**Fig. 6G**). In addition to these shared features, the subtypes exhibited some distinct molecular signatures. ITC_1 was enriched for genes associated with MSN development, including *PPP1R1B* and *RARB* **(Fig. 6F,G)**, whereas ITC_3 was instead enriched for *DSCAM* **(Fig. 6F)**, a cell adhesion molecule involved in synapse formation and neuronal self-avoidance, as well as genes (*GRM1*, *GABRA1*) previously identified in large aspiny subtypes (**Fig. 6H; Supplementary Data 5**). These molecular features suggest that ITC_1 likely represent MSN-like neurons that form densely packed intercalated islands, whereas ITC_3 may correspond to the large aspiny intercalated neurons previously described in rodent and non-human primates ^32,33^, which reside adjacent to ITC islands and interspersed throughout the intercalated space rather than forming islands themselves.

**Fig. 6.**
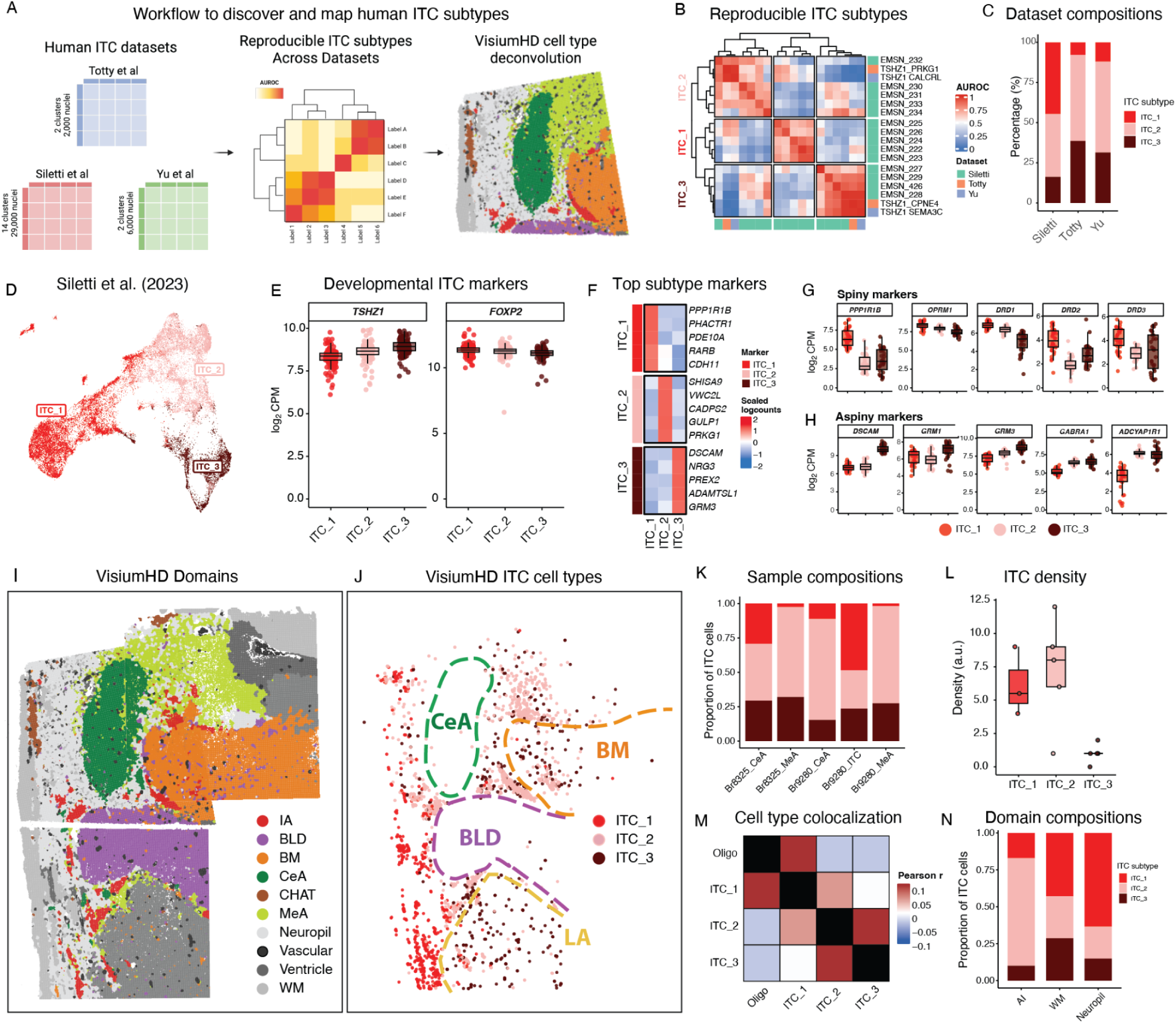
Integrative single-nucleus and high resolution spatial transcriptomics identifies distinct intercalated cell subtypes in the human amygdala. **(A)** Workflow for cross-dataset identification and spatial mapping of human ITC subtypes. Three independent snRNA-seq datasets (Totty et al.^19^; Siletti et al.^22^; Yu et al.^20^) were compared using MetaNeighbor to identify reproducible ITC clusters, and the resulting subtype labels were projected onto VisiumHD data by cell type deconvolution. **(B)** MetaNeighbor AUROC heatmap of pairwise cluster correspondence across the three datasets. Rows and columns are ordered by hierarchical clustering; three blocks of mutually high AUROC define the reproducible subtypes ITC_1, ITC_2, and ITC_3. Annotation bars indicate the dataset of origin. **(C)** Relative proportion of the three ITC subtypes within each snRNA-seq dataset. **(D)** UMAP of ITC nuclei from Siletti et al., colored by subtype. **(E)** Expression (log₂ CPM) of the canonical ITC developmental markers *TSHZ1* and *FOXP2* across subtypes. Each point is one pseudobulked donor sample from Siletti et al.; boxes show median and interquartile range. **(F)** Scaled expression (z-score) of the top five differentially expressed marker genes per ITC subtype. **(G)** Expression (log₂ CPM) of canonical and previously unreported markers of medium spiny ITC neurons, plotted as in (E). **(H)** As in (G), for markers of large aspiny ITC neurons. **(I)** BANKSY spatial domains across three VisiumHD capture arrays (16 µm bins) from donor Br9280, spanning the BLD, BM, CeA, MeA, and IA. **(J)** Spatial distribution of the three ITC subtypes in the sections shown in (I), assigned by deconvolution of VisiumHD cell segmentations. ITC_1 and ITC_2 form clustered islands in spatially distinct territories, whereas ITC_3 is distributed throughout the intercalated space, including cells bordering ITC_2 islands and within the LA and BM. Dashed outlines indicate domain boundaries from (I). **(K)** Relative proportion of ITC subtypes across the five VisiumHD samples. **(L)** Density of each ITC subtype in VisiumHD (arbitrary units). Each point is one sample; boxes show median and interquartile range. **(M)** Local spatial correlation (Pearson’s r) between ITC subtypes and oligodendrocytes. ITC_1 correlates with oligodendrocyte abundance, consistent with its localization to dense white matter tracts, whereas ITC_2 and ITC_3 correlate with one another. **(N)** Relative ITC subtype proportions within the IA, WM, and neuropil BANKSY domains.

To test these predicted identities, we mapped the three ITC subtypes onto our VisiumHD single-cell spatial data using cell type deconvolution with the Siletti et al. reference (Methods; **Fig. I,K**). Consistent with their predicted identities, we found that ITC_1 and ITC_2 localized to densely packed intercalated islands residing either within white matter tracks dorsal to the LA (ITC_1) or distributed throughout the white matter space separating neighboring amygdala subnuclei (ITC_2), whereas ITC_3 was distributed throughout the intercalated space and neighboring some island populations (**Fig. 6J,L**). Although ITC_2 and ITC_3 subtypes were consistently identified across all five VisiumHD samples, ITC_1 was primarily detected in two of the five samples (**Fig. 6K**). Cell type neighborhood analysis additionally revealed that ITC_1 neurons are preferentially colocalized with oligodendrocytes, whereas ITC_2 and ITC_3 were more likely to colocalize with one another (**Fig. 6M**). Moreover, ITC_1 was enriched with the WM spatial domain, while ITC_2 and ITC_3 were both enriched in the IA domain and sounding neuropil between subnuclei (**Fig. 6N**). Together, these findings define three molecularly and spatially distinct ITC subtypes in the human amygdala: two distinct populations forming densely packed islands (ITC_1 and ITC_2), and a diffusely distributed population found throughout the intercalated space and neighboring ITC_2 islands, potentially corresponding to the large aspiny neurons (ITC_3).

### 2.7 Spatial cell type mapping resolves anatomical subdivisions of the human central amygdala

The CeA is the principal output nucleus of the amygdala, coordinating defensive, autonomic, and appetitive responses through its striatal-like GABAergic neurons. In rodents, decades of work have revealed a rich cellular diversity within the CeA, resolving molecularly distinct and functionally opposed populations across the lateral and medial CeA subdivisions. By contrast, the cellular composition of the human CeA remains poorly resolved. To define the molecular organization of the human CeA, we integrated our Visium spatial atlas with RCTD-based cell type mapping of three putative CeA-derived inhibitory neuron populations from the Yu et al. snRNA-seq reference (**Fig. 7A-C**). Rather than occupying a common CeA domain, the three putative inhibitory neuron CeA populations (*DRD2+/ISL1*+, *DRD2+/PAX6+,* and *PRKCD+*), displayed strikingly distinct spatial distributions (**Fig. 7C**). PRKCD+ neurons localized to the CeA proper, immediately adjacent to the MeA, whereas the *DRD2+/ISL1*+ and *DRD2+/PAX6+* populations localized to neighboring regions lateral to the CeA and dorsal to the LA, corresponding to the ventral putamen (PuV) and amygdalostriatal transition area (ASt), respectively (**Fig. 7C**). We therefore re-annotated these molecularly defined domains as CeA, PuV, and ASt for downstream analyses. These regions were most clearly resolved in donors Br2743 and Br6471, which captured the broadest extent of the CeA and surrounding striatal transition areas due to their acquisition at the intermediate level of the A-P (**Fig. 7D**). Consistent with these annotations, canonical MSN marker genes (*PPP1R1B, ADORA2, TAC1, DRD1, DRD2,* and *GPR88*) were enriched in both the PuV and the ASt (**Fig. 7E,F**). In contrast, the CeA domain was characterized by expression of canonical CeA markers (*SST*, *SCG2*, *HPCAL1*, *CARTPT*, and *PNOC*) (**Fig. 7E,F**). Spatial expression of these representative markers revealed sharply restricted expression patterns that closely matched the RCTD cell type weights, confirming assignment of the *PRKCD*+ neuron class to the CeA proper and the two *DRD2+* populations to the adjacent PuV and ASt regions (**Fig. 7E,F**).

**Fig. 7:**
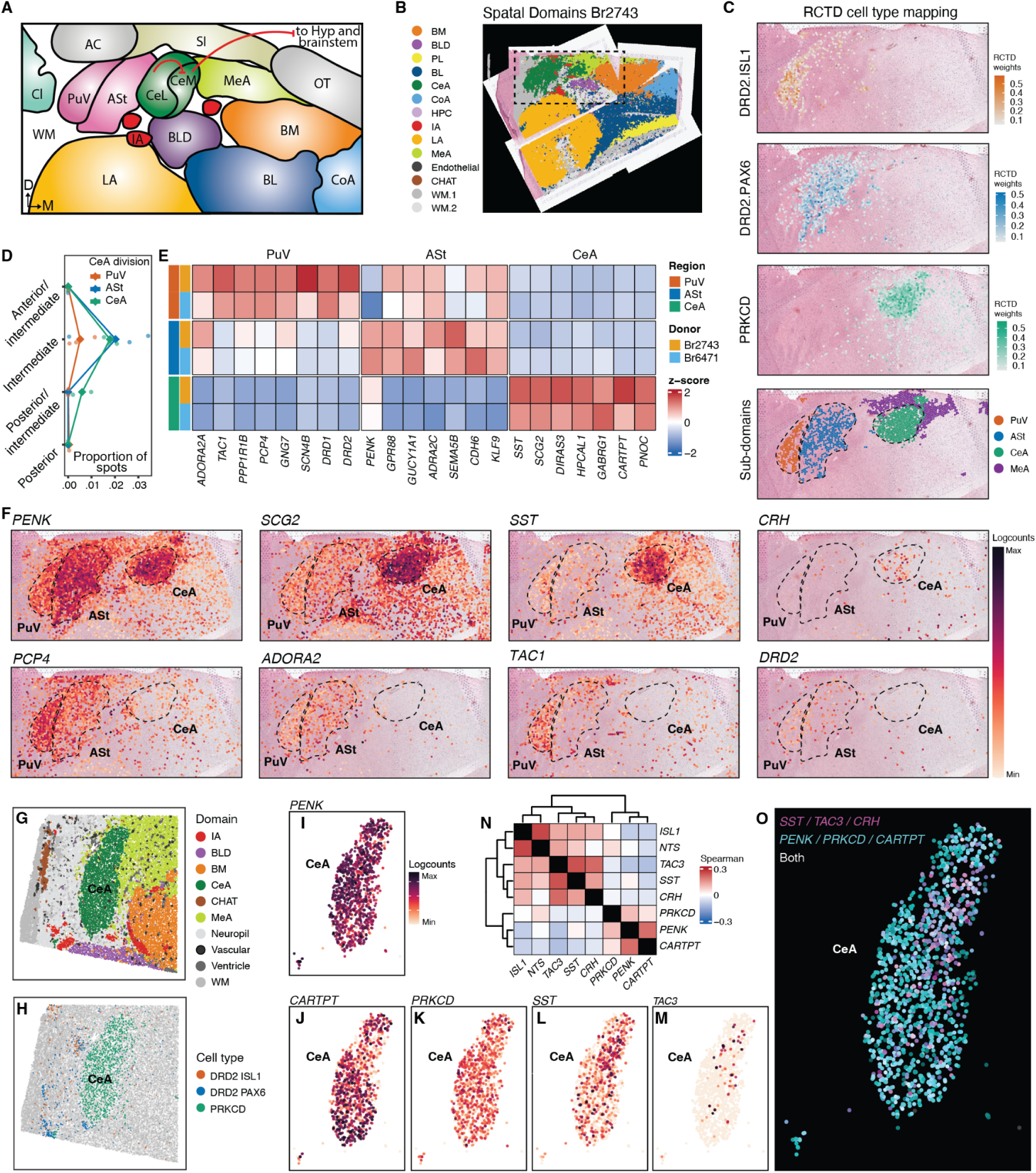
Spatial transcriptomics resolves PuV, ASt, and CeA subdivisions and reveals transcriptional heterogeneity within the human central amygdala. **(A)** Schematic of the anatomical organization of the central amygdala and adjacent structures at an intermediate anterior–posterior level. **(B)** Visium spatial domains of the intermediate anterior–posterior section from donor Br2743. Dashed box indicates the region shown in (C) and (F). **(C)** RCTD deconvolution weights for three CeA-associated cell types from the Yu et al. reference (DRD2_ISL1, DRD2_PAX6, PRKCD) projected onto Visium spots. Bottom panel shows the resulting subdivision assignments (PuV, ASt, CeA, and MeA) derived from these weights. **(D)** Abundance of each subdivision along the anterior–posterior axis, expressed as the proportion of total Visium spots per donor. Points are individual donors; lines connect means at each AP level. **(E)** Mean expression (z-scored log-normalized counts) of top marker genes for PuV, ASt, and CeA in the two intermediate-level donor samples (Br2743, Br6471). Row annotations indicate region and donor. **(F)** Spatial expression (log counts) of representative neuropeptide and canonical subdivision marker genes in Br2743. Dashed outlines mark the PuV, ASt, and CeA boundaries defined in (C); each gene is scaled independently. **(G)** BANKSY spatial domains from one VisiumHD capture array in donor Br9280, centered on the CeA. **(H)** RCTD cell type assignments for the section in (G), restricted to CeA-associated subtypes. **(I–M)** Expression of individual marker genes *PENK* (I), *CARTPT* (J), *PRKCD* (K), *SST* (L), and *TAC3* (M) across segmented cells within the CeA. **(N)** Spearman correlation of marker gene expression across CeA cells. Hierarchical clustering resolves two co-expression programs, *SST / TAC3 / CRH* and *PENK / PRKCD / CARTPT*, indicating transcriptional heterogeneity within the CeA. **(O)** Spatial distribution of cells expressing the *SST / TAC3 / CRH* program (magenta), the *PENK / PRKCD / CARTPT* program (cyan), or both (white), showing largely non-overlapping localization.

Having defined the molecular boundaries of the human CeA, we next asked whether additional molecular heterogeneity exists within the CeA itself using VisiumHD. Consistent with our Visium analysis, only the PRKCD+ population mapped to the CeA spatial domain in donor Br9280 (**Fig. 7G,H**). To determine whether the medial (CeM) and lateral (CeL) subdivisions could be resolved molecularly, we examined the spatial expression of canonical CeA neuropeptide markers at single-cell resolution. While *PENK* and *PRKCD* largely were broadly expressed throughout the CeA, *CARTPT* displayed a pronounced gap in expression in the center of the nucleus. This central region was instead enriched for *SST* and *TAC3* expression (**Fig. 7I-M**), resembling the cellular core classically attributed to the CeL. To quantify these relationships, we computed pairwise correlations of CeA marker gene expression across single cells, revealing two largely anticorrelated transcriptional programs: a *PRKCD+/PENK+/CARTPT+* program and an *SST+/TAC3+/CRH+* program **(Fig. 7N)**. Summed expression of these gene programs defined largely non-overlapping neuronal populations, with the SST-associated program concentrated within the central core of the nucleus, likely corresponding to the CeL, and the PRKCD-associated program, corresponding to the CeM, surrounding it (**Fig. 7O)**

### 2.8 Psychiatric genetics rick shows both broad neuronal enrichment and finer subnuclear organization in the human amygdala

To determine where genetic liability for complex traits is localized within the human amygdala, we applied two complementary approaches to integrate 21 trait GWAS summary statistics with spatial gene expression. gsMap^34^ partitions genome-wide heritability to individual spatial locations by linking spatial gene specificity to nearby single nucleotide polymorphisms (SNPs) and testing the resulting annotations using stratified linkage-disequilibrium score regression, whereas scDRS^35^ calculates spot-level disease scores from the expression of the top 1,000 MAGMA-prioritized genes for each trait relative to matched control gene sets (**Fig. 8A-B**).

**Fig. 8:**
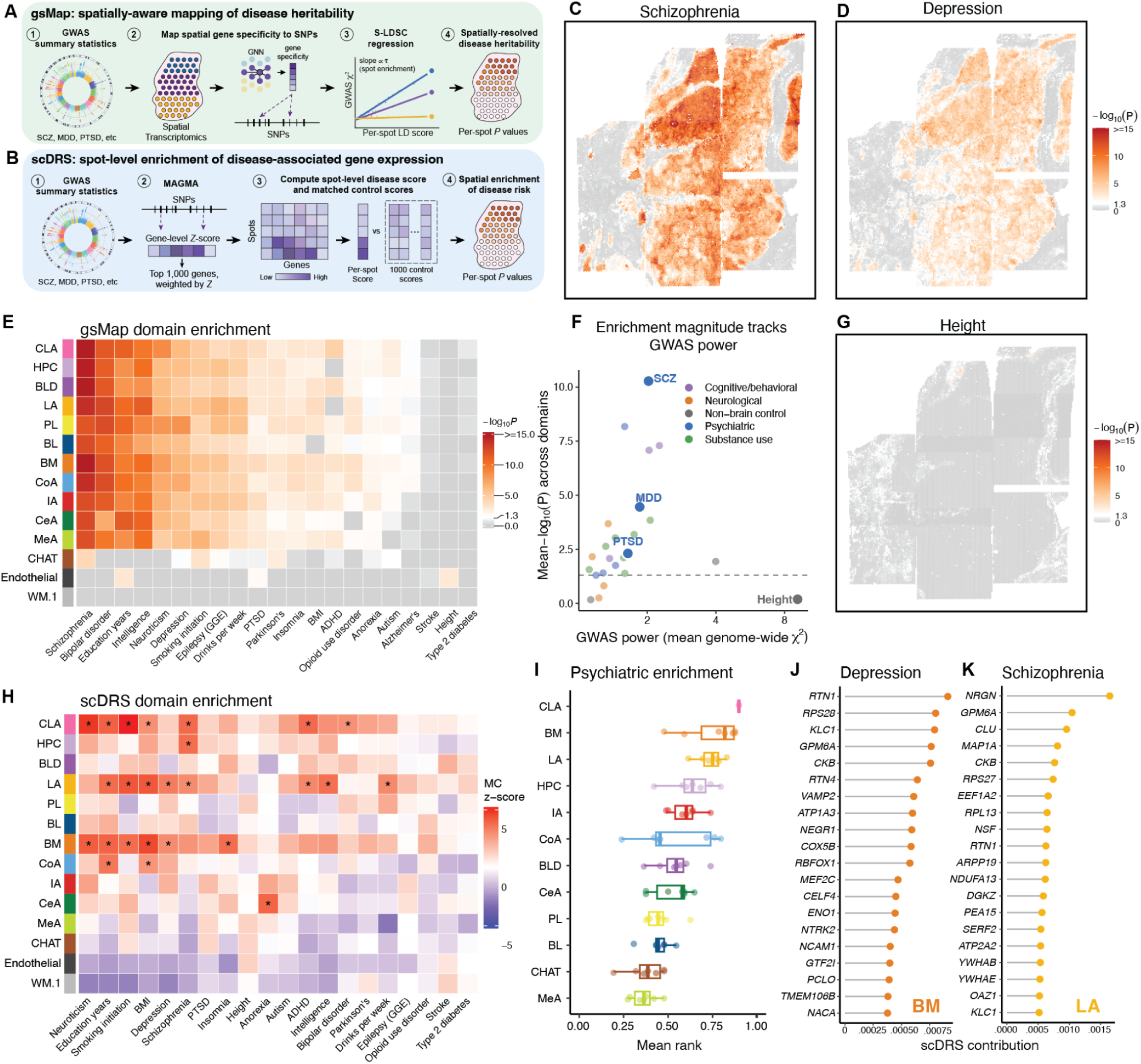
Spatial mapping of psychiatric disease heritability across the human amygdala. **(A)** Schematic of the gsMap workflow for spatial mapping of disease heritability. GWAS summary statistics are integrated with spatial transcriptomic gene expression to assign trait-relevant SNPs to spots, followed by stratified LD score regression (S-LDSC) to compute per-spot enrichment *P* values. **(B)** Schematic of the scDRS workflow for spot-level enrichment of disease-associated gene expression. GWAS summary statistics are processed through MAGMA to derive gene-level *Z* scores, and the top 1,000 genes are used to compute a spot-level disease score, which is compared against matched control gene sets to yield per-spot *P* values. **(C–D)** Spatial distribution of gsMap per-spot enrichment (−log₁₀ *P*) for schizophrenia (C) and depression (D) in a representative donor (Br8325). **(E)** Heatmap of gsMap enrichment (−log₁₀ *P*) across spatial domains (rows) and GWAS traits (columns). **(F)** Relationship between mean gsMap enrichment across spatial domains and GWAS power (mean genome-wide χ²). Traits are grouped by category: psychiatric, cognitive/behavioral, neurological, substance use, and non-brain controls. Enrichment magnitude for brain-related traits scales with GWAS power. **(G)** Spatial distribution of gsMap enrichment for height, included as a non-brain negative control trait. **(H)** Heatmap of scDRS enrichment across spatial domains (rows) and GWAS traits (columns), shown as Monte Carlo (MC) *z*-scores. Asterisks denote statistically significant domain–trait associations (FDR < 0.05). **(I)** Distribution of mean scDRS rank across psychiatric traits for each spatial domain, ordered by median enrichment. **(J–K)** Top genes contributing to scDRS enrichment for depression within the BM domain (J) and schizophrenia within the LA domain (K), ranked by scDRS contribution.

gsMap revealed widespread enrichment of psychiatric genetic risk throughout amygdala gray matter, as shown for both schizophrenia and major depression (**Fig. 8C-D)**. This pattern was consistent across donors and traits (**Fig. 8E**), with psychiatric and other brain-related traits showing strong enrichment across all gray-matter domains, but comparatively little enrichment in white matter or endothelial regions. Surprisingly, schizophrenia showed the strongest enrichment of all psychiatric traits, exceeding that observed for disorders more traditionally associated with the amygdala, including major depression and post-traumatic stress disorder (**SFig. 22**). However, we found that the magnitude of brain-related trait enrichment scales with GWAS power (**Fig. 8F**), suggesting that the particularly strong schizophrenia signal reflects the power of its GWAS rather than amygdala specificity. Conversely, height showed little spatial enrichment throughout the amygdala despite its highly powered GWAS (**Fig. 8G**), supporting the biological specificity of the broad enrichment observed for brain-related traits. This broad enrichment across gray matter is consistent with the highly polygenic nature of psychiatric traits in which liability is distributed across regulatory programs shared throughout disease-relevant neuronal populations.

In contrast to gsMap, scDRS revealed substantially greater heterogeneity among amygdala spatial domains (**Fig. 8H**). Psychiatric disease scores were preferentially elevated within a subset of amygdala domains, with the BM and LA consistently ranking among the most enriched domains across donors and psychiatric traits (**Fig. 8I**). The CLA also showed elevated psychiatric scores, although this domain was captured in only a single donor. We further decomposed representative scDRS disease scores to identify the individual genes contributing most strongly within enriched domains. Depression-associated scores in the BM were driven in part by several established MDD risk genes, including *NEGR1*, *RBFOX1*, and *PCLO* ^36^, alongside genes with roles in synaptic function and neuronal structure such as *RTN1*, *GPM6A*, *VAMP2*, and *CCK*. Similarly, the largest contributor to schizophrenia scores in the LA was *NRGN*, which is located within one of the earliest identified genome-wide significant schizophrenia loci, and encodes a protein with well-established roles in synaptic plasticity ^37,38^. Together, these analyses identify two distinct levels of spatial organization of psychiatric genetic risk: broadly distributed heritability across neuronal tissue, and finer subnuclear organization among the most strongly prioritized disease-associated genes.

## 3 Discussion

Here we combined complementary spatial transcriptomic approaches to resolve the molecular and cellular architecture of the human amygdala across multiple scales. We used transcriptome-wide, mesoscale profiling to discover novel marker genes and co-expression programs that define 13 spatial domains spanning the BLA complex, CeA, MeA, and CoA nuclei, as well as the IA islands. We additionally mapped molecularly defined cell types by integrating multiple existing human snRNA-seq references with image-based, single cell resolution spatial transcriptomics. We found that we were able to localize distinct excitatory populations to specific amygdala subnuclei, including two *LAMP5+* populations that differentially map to the magnocellular and parvocellular domains of the BL nucleus. Moreover, we used VisiumHD, which combines transcriptome-wide profiling at single cell resolution, to characterize the intricate IA islands and CeA nucleus. Combining three different single cell references, we found three robust IA cell types which show different spatial distributions, two of which formed compact islands while the third was found to be dispersed throughout the intercalated space. In the CeA, we found that two putative *DRD2+* CeA cell types mapped to the neighboring ASt and PuV, while only the *PRKCD+* cell type cluster mapped to the CeA. Despite this, we found considerable cell type diversity in the CeA that closely resembles cell types found in the rodents. Finally, we mapped common genetic risk or psychiatric disorders and other traits onto this framework, revealing both broad neuronal enrichment across amygdala neurons and finer subnuclear organization among strongly prioritized disease genes. Overall, these data provide a molecular and anatomical framework for investigating how the diverse cell types of the human amygdala contribute to behavior and disease.

A principal advance of this study was the molecular resolution of classical subdivisions of the human BLA complex. Although the magnocellular and parvocellular domains of the BLA complex have long been recognized based on cytoarchitecture and connectivity, their underlying molecular identities have remained poorly defined. By spatially mapping transcriptionally defined neuronal populations, we found that the parvocellular PL is composed predominantly of *LAMP5 + / COL25A1+* neurons, whereas the adjacent magnocellular BLD is enriched for *LAMP5+ / ABO+* population. Notably, these two domains occupy distinct positions in amygdalar circuitry in human and non-human primates. For example, the parvocellular PL is preferentially targeted by the hippocampus, receiving dense projections from the uncal region of the CA1 ^39,40^, and in turn projects to the CeA, intermediate BL, and ventral striatum, whereas the magnocellular BLD is reciprocally connected to the medial and orbital prefrontal cortex ^40^. Consistent with the increased neuronal density of the PL, our differential expression analyses found that the PL is enriched for gap junction associated genes (*GJB3, GJB5*), raising the possibility that electrical coupling between neighboring neurons promotes local synchronization that underlies coordinated oscillatory activity between the PL and target regions. Such coordination is notable given the known connectivity between PL and the hippocampal CA1 ^40^, as human intracranial recordings have shown that theta-and gamma-band coupling between the amygdala and hippocampus is dynamically engaged during aversive learning, salience processing, and memory formatting, with distinct oscillatory patterns underlying different behavioral states ^41–44^. It is likely the excitatory populations we localize here may be differentially recruited across oscillatory states, with the electrically coupled PL poised to integrate hippocampal input into broader amygdalar network activity and influencing downstream CeA and striatal circuits. The molecular and cellular diversity of these circuits may therefore provide a framework for investigating how specific amygdalar populations and oscillatory states are altered in psychiatric disorders.

Our analyses also revealed the diversity and spatial organization of the ITC system in the human amygdala. By integrating three independent human snRNA-seq datasets with our spatial transcriptomics data, we resolved three robust IA cell types that share expression of the canonical transcription factors that specify IA identity, including *FOXP2* and *TSHZ1* ^45^, but occupy distinct spatial territories. Expression of this shared dorsal lateral ganglionic eminence signature supports a common lineage for these three populations, but their spatial segregation aligns with the view that these cells represent discrete clusters with distinct connectivity and functional roles ^45^. Two populations (ITC_1 and ITC_2) formed densely packed islands in non-overlapping territories, while ITC_3 did not form islands and was instead dispersed throughout the intercalated space. ITC_1 localized between the white matter fibers dorso-lateral to the LA and BLD nuclei, while ITC_2 localized between the white matter fibers separating the neighboring amygdalar subnuclei, LA, BLD, CeA, and BM. The molecular profile of ITC_3, including enrichment for specific glutamate (*GRM1*) and GABA (*GABRA1*) receptors, was consistent with the large, aspiny intercalated neurons described in rodents ^32^ and primates ^32,33^. The distinct molecular profile further suggests functional specialization of the ITC_3 population. First, the enrichment of *DSCAM*, a homophilic cell-surface molecule that mediates cell-cell avoidance^46^, raises the possibility that differential cell adhesion contributes to its unique spatial distribution. Second, expression of *ADCYAP1R1*, which encodes the pituitary adenylate cyclase-activating polypeptide (PACAP) receptor, links this population to PAC1-dependent regulation of fear learning and extinction^47^. Given that polymorphisms in the PACAP-PAC1 pathway are associated with post-traumatic stress disorder (PTSD) symptom severity ^48,49^, these findings suggest ITC_3 as a new cellular candidate for investigating the molecular and circuit basis of PACAP-related vulnerability to PTSD. Together, these findings establish the ITC system in the human amygdala as a heterogenous collection of spatiomolecularly distinct.

Our findings also provides unprecedented insight into the molecular organization of the human CeA, one of the principal output nuclei of the amygdala that is composed predominantly of GABAergic neurons and plays a central role in defensive behaviors, autonomic regulation and emotional learning ^30^. Prior work in rodents and non-human primates has demonstrated that the CeL and CeM subdivisions comprise molecularly and functionally distinct inhibitory populations ^12,17,21,50–53^. In humans, however, the molecular organization of the CeA has remained poorly understood. By integrating existing snRNA-seq reference data with our spatial atlas, we revealed that several inhibitory populations previously annotated as CeA cell types instead localized to neighboring regions. Mapping putative CeA inhibitory populations from Yu et al. ^20^, only the *PRKCD+* population localized to the CeA, while the *DRD2+ / ILS1+* and *DRD2+ / PAX6+* populations mapped to the adjacent PuV and ASt. These revised annotations were supported by canonical MSN signatures and further corroborated by a recent multimodal atlas of the macaque striatum, which identified two novel *RGS6+* MSN subtypes, which were distinguished by *ISL1* and *PAX6* and localized to the PuV and ASt, respectively ^54^. Examining distribution of cell types and spatial gene expression gradients within the CeA, we noted that canonical CeA marker genes were not uniformly distributed, but rather partitioned into two largely anticorrelated transcriptional programs consisting of either *PRKCD* / *PENK* / *CARTPT* or *SST* / *TAC3* / *CRH*. This molecular dichotomy recapitulates the lateral vs medial divisions, which have been extensively characterized in rodents ^12,50^ These data provide the first evidence that these opposing neuropeptidergic populations, which correlate with anatomically localized CeA microcircuits in rodents, are conserved in humans. We again highlight that the snRNA-seq reference produced only a single CeA cluster, and that resolving cell type heterogeneity was only possible with spatial context, suggesting that the rich cellular diversity of CeA remains substantially under-sampled in available snRNA-seq atlases. Future studies integrating larger, more deeply sequenced snRNA-seq datasets with high resolution spatial transcriptomics will be necessary to resolve the rich diversity of neuropeptidergic and functionally specialized neuronal populations that underlie CeA function.

Mapping common genetic risk onto our spatial framework revealed two levels of organization within the human amygdala. Partitioning genome-wide heritability with gsMap produced broad enrichment across gray-matter domains with relatively little subnuclear variability, whereas scDRS revealed finer anatomical structure among strongly prioritized disease-associated genes, particularly within the BM and LA. These findings likely reflect complementary features of highly polygenic architecture. The broad gsMap signal is compatible with the omnigenic model ^55^, in which disease heritability is distributed across extensive regulatory networks active within disease-relevant cell populations. Shared neuronal programs may therefore dominate genome-wide enrichment, distinguishing neuron-rich gray matter from white-matter and endothelial regions while providing relatively little resolution among amygdala nuclei. In contrast, the more restricted set of highly weighted genes used by scDRS may capture disease-associated programs that are expressed less uniformly across nuclei, revealing spatial organization obscured at the level of genome-wide heritability. We must also caution comparisons across traits given the strong correlation to GWAS power. The magnitude of gsMap enrichment closely tracked genome-wide association strength, consistent with the recognized sensitivity of spatial heritability mapping to differences in GWAS power ^34^. For example, the particularly strong schizophrenia signal should not be interpreted as evidence that the amygdala is more centrally involved in schizophrenia than in depression or PTSD, and the absence of significant subnuclear enrichment for PTSD does not provide evidence against amygdala involvement. PTSD had comparatively weak genome-wide signal in our analysis and is known to have lower genetic discoverability than most other psychiatric disorders, even in recent large-scale GWAS ^56,57^. Larger and more deeply phenotyped studies may therefore be required before PTSD genetic liability can be localized at the level of individual amygdala nuclei. Importantly, GWAS power alone was not sufficient to generate enrichment given that height showed little amygdala enrichment despite a highly powered GWAS, indicating that power modulates the sensitivity of biologically specific signals.

A key strength of our study is the integration of complementary spatial transcriptomic technologies, enabling both transcriptome-wide profiling and single-cell resolution across the human amygdala. Nevertheless, several limitations should be considered. First, the size and anatomical complexity of the human amygdala make comprehensive spatial profiling inherently challenging, and no single technology currently provides simultaneous whole transcriptome coverage, single-cell resolution and complete anatomical coverage. Second, our analyses were necessarily limited to a subset of coronal levels, and we observed substantial variation in domain and cell type representation across the A-P axis. Certain structures, including the CeA and surrounding striatal transition areas, were most comprehensively sampled at intermediate A-P levels, emphasizing the need for future studies spanning the A-P extent to fully resolve regional cellular diversity. Finally, our cell type mapping is fundamentally constrained by the available snRNA-seq references. Although spatial deconvolution accurately localized well-defined neuronal populations, it remains limited by under-clustered reference populations, particularly within the CeA subpopulations, and by developmentally dynamic cell states that may be incompletely represented in atlases derived from adult donors

Together, these data establish a cellular, molecular and anatomical framework for the human amygdala by placing transcriptionally defined cell types within their native spatial context. In doing so, they resolve long-standing ambiguities in human amygdala organization, including the molecular identities of magnocellular and parvocellular divisions of the BLA complex, the spatial organization of distinct intercalated cell populations, and the cellular composition of the CeA. More broadly, our findings demonstrate that spatial context is essential for accurately interpreting transcriptionally defined neuronal populations, and to localize disease-associated molecular changes within anatomically defined microcircuits of the human amygdala. We anticipate that this resource will provide a foundation for future studies investigating the development, evolution, circuit organization and disease associated changes in the human amygdala and linking the diverse cell types of the human amygdala to the behaviors they support.

## 4 Methods

### 4.1 Postmortem human tissue samples

Postmortem human brain tissue from neurotypical adult donors of European ancestry (*N* = 10) were obtained at the time of autopsy following informed consent from legal next-of-kin, through the Maryland Department of Health IRB protocol #12–24, and from the Department of Pathology at Western Michigan University Homer Stryker MD School of Medicine, the Department of Pathology at University of North Dakota School of Medicine and Health Sciences, and Gift of Life Michigan, all under the WCG protocol #20111080. Using a standardized strategy, all donors were subjected to clinical characterization and diagnosis. Macroscopic and microscopic neuropathological examinations were performed, and subjects with evidence of significant neuropathology were excluded. Additional details regarding tissue acquisition, processing, dissection, clinical characterization, diagnoses, neuropathological examination, RNA extraction and quality control (QC) measures have been previously published^114^. Demographic information for all neurotypical control donors is listed in **Supplementary Table 1**. Each tissue block (approximately 25 X 25 X 10 mm) was dissected from a frozen coronal slab of the medial temporal lobe at the level of the amygdala using a hand-held dental drill. Tissue blocks were stored in sealed cryogenic bags at -80°C until cryosectioning.

### 4.2 Tissue processing and quality control

The following landmarks were used to guide the amygdala (AMY) dissections: anterior commissure dorsally, the optic tract medially, the entorhinal cortex ventrally, and the claustrum laterally. All tissue blocks were subjected to quality control by H&E to ensure dissected blocks contained the entire amygdala. Fresh frozen tissue blocks were acclimated to -14°C for 30 minutes inside the cryostat (Leica CM3050s), mounted on a round chuck with Optimal Temperature Compound (TissueTek Sakura, Cat #4583), and ∼50 µm of tissue was trimmed and discarded to achieve a flat surface. Several ∼10 µm sections were collected on pre-chilled microscope slides (VWR SuperFrost Microscope Slides, Cat #48311703). H&E staining was performed according to the manufacturer’s instructions as previously described^57^, and images were acquired using an Aperio CS2 slide scanner (Leica). To prepare tissue for the Xenium assay, four blocks were acclimated in the cryostat, trimmed to a flat surface, and the AMY was scored out of the block. One 10 µm section was mounted onto the pre-chilled Xenium Spatial Gene expression slide (Part Number 1000460, 10x Genomics), per donor. Following Xenium, all blocks were prepared for the Visium assay: acclimated in the cryostat, trimmed to a flat surface, and the areas of the blocks corresponding to the AMY were scored with a razor blade vertically in ∼6.5 mm strips to match the width of the Visium capture areas. Because each AMY occupied six to eight capture arrays, multiple non-overlapping, horizontally-adjacent or partially-overlapping vertically-adjacent 10 µm tissue strips containing AMY were mounted onto pre-chilled Visium Spatial Gene Expression slides (part number 2000233, 10x Genomics) to cover the entire AMY. Several unscored and scored sections were also mounted onto regular pre-chilled glass slides and banked at -80°C for later use. Following successful completion of the Visium assay, previously banked sections from two blocks were selected for the VisiumHD assay, including specific Regions of Interests (ROIs), such as IA, MeA, and CeA.

### Cresyl Violet staining

Fresh-frozen human tissue sections (20 µm) were air-dried at room temperature (RT) for 1 h and fixed in 10% neutral buffered formalin (NBF; Cat. No. HT501128-4l, Sigma-Aldrich, St. Louis, MO, USA) for 30 min. Sections were washed twice in 1× PBS for 3 min each, followed by a 2 min rinse in deionized (DI) water. Slides were then incubated in 45 ml of filtered Cresyl Violet staining solution (Cat. No. C5042-10G, Sigma-Aldrich, St. Louis, MO, USA) for 20 min and rinsed twice in DI water. Differentiation was performed in 95% ethanol for 1 min, followed by dehydration through graded ethanol solutions (70%, 95%, and 100%; 3 min each). Sections were subsequently cleared in xylene (2 × 3 min), coverslipped with 75 µl of EcoMount mounting medium, and allowed to dry overnight before imaging. High-resolution brightfield images were acquired using Leica Aperio CS2 slide scanner equipped with a 20x/0.75NA objective and a 2x doubler.

### 4.3 Spatially resolved transcriptomics (SRT) data generation

#### Visium Data Generation

Visium Spatial Gene Expression slides were processed as previously described^57^. Tissue optimization experiments were performed according to the manufacturer’s protocol (CG000160, revision B, 10x Genomics) to ensure optimal permeabilization time. Briefly, eight AMY tissue sections were exposed to permeabilization enzymes for 3 to 36 min. cDNA synthesis was performed using a fluorescently labelled nucleotide (CG000238, revision D, 10x Genomics). The slide was then coverslipped and fluorescent images were acquired at 10x magnification with a TRITC filter (ex 550nm/em 600nm) on a Cytation C10 Confocal Imaging Reader (Agilent). The optimal permeabilization time of 18 min was selected for all subsequent experiments.

Six to eight 10μm sections from each of the 10 AMY blocks were mounted onto two 10x Visium Gene Expression slides, and processed according to the manufacturer’s instructions (10x Visium Gene Expression protocol number CG000239, Rev G) as previously described^57^. Briefly, H&E staining was performed (protocol CG000160, revision B, 10x Genomics), after which slides were coverslipped and high-resolution, brightfield images were acquired on a Leica CS2 slide scanner equipped with a 20x/0.75NA objective and a 2x doubler. Following the removal of the coverslips, tissue was permeabilized, cDNA synthesis was performed, and sequencing libraries were generated for all capture areas following the manufacturer’s protocol. Libraries were loaded at 300 pM and sequenced on a NovaSeq 6000 (Illumina) at the Johns Hopkins Single Cell Transcriptomics core according to the manufacturer’s instructions at a minimum depth of 60,000 read pairs per spot.

### Visium HD Data Generation

Tissue slides banked for Visium HD were processed in pairs using the Visium HD Spatial Gene Expression platform by 10x Genomics (Pleasanton, CA). After slides were removed from -80°C storage, tissues were fixed in chilled methanol (Millipore sigma 322415), dehydrated in isopropanol (Millipore Sigma I9516), and H&E-stained following the procedures adapted from CG000684 rev B. In brief, Gills II Hematoxylin (Leica 3801520) was applied uniformly over the tissues. After a 1-minute incubation, the hematoxylin was discarded and tissues were washed 3x in Milli-Q ultrapure water (800mL/wash) (Thermo Scientific 50131948) and then incubated in Bluing Buffer (Agilent CS70230) for 1 minute at room temperature. After an additional wash in Milli-Q ultrapure water (800mL/wash), slides were stained in Alcoholic Eosin (Leica 3801615) and washed 3x in Milli-Q ultrapure water (800mL/wash). Slides were then mounted in 85% glycerol (Acros Organics 327255000) diluted in 0.2mm filtered nuclease-free water (IDT 11-05-01-04) and 600U of Protector RNase inhibitor (Roche RNAIHN-RO). Slides were images with a 40x 0.75 NA objective mounted on a Leica Aperio CS2 digital pathology slide scanner (Leica Biosystems). Following coverslip removal, ROIs for both tissue slides were selected via precise application of the Visium Cassette S3 gasket (CG000730 Rev A). ROIs were subsequently destained in 0.1N HCL (Fisher Chemical SA54-1) for 15 minutes at 42°C and washed 3x in Tris-EDTA pH 8 (ThermoFisher Scientific BP24731) followed by 1 wash of 1x PBS pH 7.4 (ThermoFisher Scientific AM9624). Steps pertaining to *In situ* hybridization, ligation, CytAssist-Enabled probe release, and library construction steps were then followed according to the procedures outlined in CG000685 rev B. Following a 15-minute permeabilization at room temperature in a 0.7% Tween-20 (Thermo Fisher Scientific 28320) 1xPBS pH 7.4 solution, the Visium Human Transcriptome probe kit v2 (PN-1000466) was applied to each ROI and allowed to hybridize to accessible RNA at 50°C overnight. Hybridized tissues were then washed in FFPE Post Hyb Wash Buffer (PN-2000424) and ligated using a ligation enzyme (PN-2000425) diluted in 2x probe ligation buffer (PN-2000445) for 1 hour at 37°C. While tissues were washed in post ligation wash buffer (PN-2000419) and then allowed to acclimate to room temperature, a Visium HD slide (PN-1000670) was removed from -80°C storage warmed to room temperature for 30-60 minutes, after which it was washed 3x in a total of 60mL 0.1x SCC (Millipore Sigma S66391L). If applicable, additional washes were implemented until both 6.5mm spatially-barcoded oligonucleotide capture arrays were visibly free of debris. The Visium HD slide was then placed in a Visium 2-port cassette S2 (PN-1000669) and RNase enzyme (PN-3000605) was allowed to equilibrate the arrays. During the Visium HD slide equilibration, the two tissue slides were stained in 10% Alcoholic Eosin, washed 3x in 1mL 1xPBS, and then mounted on the Visium CytAssist instrument (PN-1000441). After the Visium HD slide equilibration and subsequent drying, the CytAssist run was initiated, during which time ligated probes on tissue slides were transferred and captured onto the Visium HD arrays. Immediately upon completion of the run, the Visium HD slide was washed in 3x in Buffer EB (1mL/wash) (Qiagen, 19086) and placed in a Visium 2-port cassette. Extension enzyme (PN-2000389) and extension buffer (PN-2000409) were then applied to each capture array and probes were extended over two cycles of a 30min incubation at 53°C. Following probe extension, probes were eluted and then pre-amplified using AMP B (PN-2000567) and TS primer Mix B (PN-2000567) to generate ample material for downstream library construction. To determine the ideal number of additional cycles needed to amplify each library for the sample index PCR, pre-amplified samples were quantified using KAPA SYBR Fast qPCR Master Mix Universal (Roche KK4600) run on the CFX Opus 96 (Bio-Rad 12011319). Unique indices from the dual index plate TS Set A (PN-3000511) were assigned to each sample so that Visium HD libraries could be pooled and demultiplexed in downstream sequencing runs. CytAssist Spatial Gene Expression libraries were sequenced on the Illumina NovaSeq X Series 25B flow cell, targeting ∼25000 reads per spot at Psomagen (Rockville, MD).

### Xenium Data Generation

AMY tissue sections from a subset of 4 donors were collected on individual Xenium slides. Xenium slides were fixed and permeabilized according to 10X Genomics’ Xenium *In Situ* for Fresh Frozen Tissues – Fixation & Permeabilization Protocol (CG000581, Rev C). Briefly, slides were warmed for 1 min prior to submersion in 3.7% formaldehyde for 30 min. Slides were then permeabilized with 1% sodium dodecyl sulfate solution (SDS) for 2 min and rinsed with PBS. This was followed by a 1 h incubation in 70% methanol on ice. Following PBS washes, slides were placed in their respective cassettes and incubated with 0.05% PBS-T. For probe hybridization, ligation, and amplification, the 10X Genomics protocol: Xenium In Situ Gene Expression was followed (CG000582, Rev E). The probe hybridization solution was then prepared by combining the 10X Genomics off-the shelf human brain probe panel (Xenium Human Brain Gene Expression Panel, Part No.1000599) with a custom probe panel (10X Genomics; Xenium Custom Gene Expression Panel, Part No. 1000651; **Table 1**). Probe mix was applied to each slide, slides were covered with a lid and incubated at 50°C overnight.

The next day, the slides were washed twice in 1X PBS-T followed by an incubated wash in post hybridization wash buffer (Xenium Post Hybridization Wash Buffer, Part No. 2000395, 10X Genomics). Three washes of 1X PBS-T were completed before addition of ligation mix (Xenium Ligation Buffer, Part No. 2000391, Xenium Ligation Enzyme A, Part No. 2000397 and Xenium Ligation Enzyme B, Part No. 2000398, 10X Genomics) to each slide. Slides were then incubated at 37°C for 2 h. After ligation, the slides were washed three times in 1X PBS-T and then incubated in an amplification master mix (Xenium Amplification Mix, Part No. 2000392 and Xenium Amplification, Part No. 2000399, 10X Genomics) at 30°C for 2 h. After amplification, slides were washed two times in TE buffer followed by three washes in 1X PBS and then incubated in diluted reducing reagent B (Reducing Agent B, Part No. 2000087, 10X Genomics) for 10 min at room temperature. Slides were then washed in 70% ethanol followed by two washes of 100% ethanol. Autofluorescence solution (Xenium Autofluorescence Mix, Part No. 2000753, 10X Genomics) was then added to each slide and incubated in the dark for 10 min at room temperature. Slides were then rinsed three times with 100% ethanol before drying on a 37°C preheated thermocycler for 5 min. Additional washes were performed before adding Xenium Nuclei Staining Buffer (Part No. 2000762, 10X Genomics) which incubated for 1 min in the dark at room temperature.

Slides were then washed three times in 1X PBS-T and stored at 4°C prior to being loaded on the Xenium analyzer at Psomagen (Rockville, MD). The solutions needed to run the Xenium Analyzer (10x Genomics) were prepared according to the manufacturer’s instructions. Imaging of all slides took place on the Xenium Analyzer.

### 4.4 Spatial Transcriptomics data processing and analysis

#### Image processing

For both standard Visium and VisiumHD, sample slide H&E images were processed using `VistoSeg` (Tippani et al., 2023). Briefly, the `splitSlide` function was used to segment each full-slide image into separate capture-area images corresponding to the four Visium arrays (A1, B1, C1, and D1). These capture-area images were then provided as input to `spaceranger` (10x Genomics, v3.0.1) for downstream alignment and quantification. Standard Visium capture areas with overlapping regions from a single brain donor were combined into a single sample using the `VisiumStitched` package ^58^ (**SFig. 1-7**). Tissue landmarks within the overlapping regions were visually identified, and stitching was performed using Fiji with the TrakEM2 plugin. Slide and capture area metadata were saved for downstream batch correct and dataset integration. No further image processing was performed for Xenium.

#### Raw data processing

For Visium, the capture-area images generated by `VistoSeg` were next registered to their corresponding spot grids in Loupe Browser (10x Genomics) to ensure accurate alignment between histology and spatial barcodes. Samples were then processed with `spaceranger` v3.0.1, using the JSON file obtained from the `Loupe Browser`, the capture-area image, and associated FASTQ files to generate spatial feature-count matrices. For VisiumHD, capture-area images were automatically registered to the underlying 2 micron bins using `spaceranger` v4.0.1, and automatic cell segmentation was performed on the H&E images with a nuclear expansion distance of 5 um. By default, `spaceranger` also exports count matrices for 8 µm and 16 µm square bins, which were used for spatial domain detection. For Xenium, cell segmentation was performed using `xeniumranger` v2.0.0 (10x Genomics) with a nuclear expansion distance of 5 um. Resulting spatial feature-count matrices from each platform were output to R for downstream analyses.

### Quality control and normalization

Across all platforms, standard filtering was performed to remove genes with zero UMI counts across all spots, spots with all-zero counts, and spots outside the tissue mask defined in `Loupe Browser`. Spot-and cell-level quality-control metrics were added using `addPerCellQC()` from the `scuttle` Bioconductor package, including library size (total UMIs per spot), number of detected genes, and mitochondrial expression rate (McCarthy et al., 2017). For standard Visium, we used `SpotSweeper` to identify and remove local outliers based on spatial neighborhoods^59^. Briefly, `SpotSweeper` computes a neighborhood-based z-score for each spot by comparing its QC value to the distribution of values in nearby spots, enabling detection of spatially focal artifacts that may not appear as global outliers. We applied this procedure to all three QC metrics (library size, number of detected genes, and mitochondrial percentage) and excluded spots with local z-scores indicating markedly reduced gene complexity (local z < −3 for UMIs and detected genes) or elevated mitochondrial percentage (local z > 3 for mitochondrial rate). Finally, we applied a conservative filter to all datasets to remove any remaining spots/cells with very little biological information for each Visium (< 50 unique genes and/or total UMIs), VisiumHD (< 10 genes and/or UMIs), and Xenium (< 10 genes and/or UMIs).

For Visium and VisiumHD, raw count matrices were normalized using size-factor-based log-normalization with the `logNormCounts` function from the `scuttle` R/Bioconductor package. Counts for each spot/cell were scaled by the corresponding library size factor and log-transformed as log2-normalized counts with a pseudocount. The resulting log-normalized expression values were for all downstream analyses otherwise stated. For Xenium data, we used cell-size based log-normalization with `normalizeCounts` from the `scuttle` package with the nucleus area serving as the size factors. We used area-based rather than library-size-based normalization to avoid potential biases introduced by the targeted gene panel composition ^60^.

### Feature selection

For standard Visium, we identified spatially variable genes using the `nnSVG` Bioconductor package ^61^. `nnSVG` was run separately within each sample after filtering lowly expressed genes with `filter_genes` and re-computing log2-normalized expression with `logNormCounts` via the `scater` R/Bioconductor package. For each sample, `nnSVG` produced a ranked list of genes by spatial variance. We then integrated results across samples as recommended by averaging gene ranks across samples and counting how often each gene appeared in the top 1,000 SVGs per sample. We defined “replicated” SVGs as genes in the top 1,000 in at least two samples (1,487 genes) and ranked this set by mean rank. For VisiumHD feature selection prior to spatial clustering, highly variable genes were identified with modelGeneVar from the scran R/Bioconductor package using log-normalized counts. The top 2,000 HVGs were selected from downstream analysis. To ensure that we could discover small IA islands, we additionally included the top 100 marker genes discovered from the standard Visium IA domains (see details below), for a total of 2,100 features. All 366 genes in the Xenium gene panel were used for spatial clustering of Xenium datasets.

### Unsupervised clustering of stitched Visium data

Because each coronal section was tiled across 6-8 adjacent capture areas and computationally reconstructed, we devised a robust, reference-guided method to enable multi-donor clustering while reducing the impact of technical batch and slide effects. We first performed unsupervised spatial clustering on a single intermediate section from a well-characterized donor (Br8325) using the `BayesSpace` package, applied to batch-corrected principal components derived from the top 2,000 SVGs. `Harmony` batch correction was applied across capture arrays for this initial step. Resulting spatial domains were manually annotated by alignment to the Human Brain Atlas ^27^. We then used `findMarkers` from the `scran` package to identify the top 100 differentially expressed genes per spatial domain from this reference donor, yielding a curated set of 1,600 domain-discriminating marker genes free of technical or donor effects.

To enable robust multi-donor clustering, we re-computed principal components using only these 1,600 marker genes across all donors, followed by `Harmony` batch correction across both capture arrays and donors. To perform joint spatial clustering across donors, we introduced spatial coordinates offsets to place all samples in a shared coordinate space, as recommended by `BayesSpace`. `BayesSpace` clustering was then applied to this joint coordinate space embedding with k=16 clusters and a spatial smoothing parameter of gamma=3 over 10,000 iterations. This strategy worked exceedingly well to discover well-established amygdalar domains that were consistent across donors, while also not prohibiting the discovery of domains specific to only one donor. For example, the most posterior sample, Br6660, was found to contain a large portion of the claustrum that was not captured in any of the other donors.

Intercalated cell islands (AI), which are small and scattered in between larger domains, proved difficult to reliably capture with `BayesSpace` clustering. To identify them separately, we performed cell type mapping using non-negative matrix factorization (NMF) to discover IA-specific cell type patterns from the Yu et al snRNA-seq dataset ^20^, and then projected these patterns into the Visium data. We have successfully used this cell type-mapping strategy before ^62^. Using the `RcppML` package ^63^, we first used cross-validation to determine the optimal rank (k), which was determined to be k=50. We then performed NMF on logcounts from the Yu et al human dataset. NMF factor weights were then projected onto the Visium data, and factors enriched for IA cell type signatures (factors 44 and 65) were identified. Putative IA spots were classified using Gaussian mixture models (k=2) applied to the projected factor weights, where spots assigned to the high-value component for either factor were labeled as IA. To reduce false positives, labels were spatially smoothed using a k-nearest neighbors approach, requiring at least 50% spatial neighbors to be classified as IA to retain the label. IA annotations were confirmed by high expression levels of *TSHZ1* and *FOXP2,* known developmental transcription factors of amygdalar intercalated neurons. We then integrated these annotations into the final domain labels for all donors.

### Unsupervised clustering of Xenium and Visium HD data

Spatial domains in the Xenium data were identified using the BANKSY R package ^64^. Feature selection was restricted to the 366 targeted gene panel, BANKSY neighborhood matrices were computed on nucleus-based normalized counts using a geometric neighborhood of `k_geom=36` and embedded by BANKSY PCA (50 PCs) at `lambda=0.8` to increase spatial smoothing and bias clustering towards coherent domains, as recommended by BANKSY. To integrate across samples, per-sample array coordinates were shifted into a shared, non-overlapping coordinate space. Sample batch effects were then corrected by running `Harmony` on the BANKSY embedding across samples, and domains were identified using Leiden clustering on the Harmony-corrected embeddings using a sweep of clustering resolutions (`resolution` was set to 0.8 - 2.0 in intervals of 0.2) and found that `resolution=1.8` best resolved the anatomical structure of the amygdala. These clusters were then manually annotated based on their gene expression profile and anatomical location.

To discover coherent spatial domains in the VisiumHD data, we used 16 µm bins output by spaceranger rather than single cell segmentations. As mentioned above, spatial clustering was run on 2,100 total features composed of the top 2,000 highly variable genes and, to preserve sensitivity to IA islands, the top 100 IA marker genes from the Visium dataset. BANKSY neighborhood matrices were computed on log-normalized counts with the azimuthal Gabor filter enabled (`compute_agf=TRUE`) using geometric neighborhoods of the 25 and 50 nearest neighbors (`k_geom=c(25, 50)`), and embedded by BANKSY PCA (50 PCs) with `lambda=0.8.` As with Xenium, per-sample array coordinates were shifted into a shared, non-overlapping coordinate space for cross-sample integration. Sample batch effects were then corrected by running `Harmony` on the BANKSY embedding across samples, and domains were identified using Leiden clustering on the Harmony-corrected embeddings using a sweep of clustering resolutions (`resolution` was set to 0.2 - 1.2 in intervals of 0.2). A `resolution` of 0.6 best resolved the amygdala subnuclei and intercalated islands. These clusters were also manually annotated based on their gene expression profile and anatomical location.

### Comparison of Xenium and Visium spatial domains

To assess how well Visium and Xenium spatial domain corresponded to one another, we computed the Pearson correlation of the gene expression between domains using the 366 genes in the targeted Xenium gene panel by performing spatial registration from the `spatialLIBD` framework ^65^. Each dataset was pseudobulked by spatial domain with `registration_pseudobulk` and domain-level enrichment statistics were computed for each platform via `registration_model, registration_block_cor`, and `registration_stats_enrichment`, resulting in per-gene enrichment *t*-statistics for every domain.Treating Visium domains as the reference, we correlated the Xenium per-gene enrichment t-statistics against the Visium enrichment modeling results using `layer_stat_cor`, and visualized the resulting domain-by-domain correlation matrix as a heatmap (l`ayer_stat_cor_plot`). Only amygdalar domains were retained for plotting. For individual domains of interest (LA and PL), the t-statistics were additionally compared directly between platforms as scatterplots, annotated with the Pearson correlation coefficient and the top five concordantly enriched genes.

### Marker Gene Detection Across Spatial Domains

Marker genes for the 16 Visium BayesSpace domains were identified using two complementary approaches. First, for discovering domain-specific features to use for multi-sample clustering, spot-level enrichment markers were computed with findMarkers from the scran package grouping spots by spatial domain, using a Wilcoxon rank-sum test. The top 100 genes per domain were retained and used for the multi-sample clustering across all donors. To discover domain-specific marker genes that were robustly expressed across all donors, we used a pseudobulk approach by applying the `spatialLIBD` spatial registration framework ^65^. In short, gene counts were aggregated across all spots within each spatial domain and donor using `registration_pseudobulk,`requiring a minimum of 10 spots per pseudobulk sample. PCA on the pseudobulked logcounts was inspected against spatial domain, donor, and QC metrics to assess source of variance. Six pseudobulk samples driving the top principal component were flagged as low-library-size outliers and dropped prior to differentially expression analysis. Domain-level models were fit with `registration_model` and within-donor correlation was estimated with `registration_block_cor` and supplied as a blocking term. We use the enrichment model statistics which contrasts each domain against all others to define domain-specific marker genes, defined as genes with FDR < 0.05 and log_2_ fold-change > 1.5. These statistics were visualized as volcano plots (SFig. 13) and pseudobulk expression boxplots. The top 50 marker genes per domain can be found in **Supplementary Data 1**.

### Rasterization of Xenium gene expression and cell counts

To improve visualization, Xenium gene expression was rasterized into a uniform grid using `SEraster` ^66^. Raw counts were summed within regularly tiled hexagonal bins via `rasterizeGeneExpression` at a resolution of 259 pixels (∼55 um). Summed gene expression was then visualized using the `plotRaster` function. A similar approach was used to quantify the density of excitatory neurons. In short, the number of excitatory neurons were summed within hexagonal bins at a resolution of 259 pixels, providing a density estimation (number of cells / 55 um). The distribution of excitatory neuron density was then calculated per BANKSY spatial domain.

### Spatial co-expression analysis construction

Spatial gene co-expression networks were constructed using the `Smoothie` Python package ^67^ applied to all 7 Visium samples. Quality control filtering was performed to remove spots with fewer than 50 total counts (`filter_cells`) and genes detected in fewer than 10 total spots or with fewer than 100 total counts (`filter_genes function`). Expression values were normalized using counts-per-thousand (CPT, `target_sum=1000`) followed by log1p transformation. Gene counts were then smoothed spatially using Smoothie’s Gaussian kernel with the Gaussian standard set to 415.5 pixels, corresponding to approximately 150 microns based on a pixel-to-micron ratio of 2.77 (median nearest-neighbor spot distance of 277 pixels at ∼100um spot spacing in Visium). In-place smoothing (`grid_based=FALSE`) was used as recommended for cell-sized and binned resolution data, with a minimum of 3 spots required under each Gaussian kernel (`min_spots_under_gaussian=3`).

Smoothed expression matrices from all 7 donors were concatenated and pairwise Pearson correlation coefficients were computed via Smoothie’s `compute_correlation` across approximately 17,963 shared genes. Diagnostic analysis following Wang et al (2022) revealed strong mean-correlation bias in the resulting correlation matrix: the interquartile range (IQR) of correlations for the highest-expression gene bin pairs was approximately 13-fold greater than for the lowest-expression bin pairs (IQR = 0.317 vs 0.023), indicating that highly expressed genes exhibited systematically inflated correlation while lowly expressed gene had correlations compressed towards zero ^68^. To correct for this, we implemented spatial quantile normalization (SpQN) as a pure Python/NumPy port of the R/Bioconductor span package. The corrected correlation matrix was computed using `nomalize_correlation`. This sorts genes by mean expression, partitions them into 20 overlapping expression-level bins, and applies quantile normalization within each bin pair to equalize correlation distributions, using the highest-expression bin as the reference distribution. After correction, IQR values were largely uniform across all expression bin pairs, confirming successful removal of the mean-correlation bias. The Python implementation of SpQN (SpQNpy) can be found at https://github.com/MicTott/SpQNpy.

Network construction and module detection was carried out using standard Smoothie workflows. Hyperparameter selection was performed using `select_clustering_params` by sweeping Pearson correlation coefficient (PCC) cutoffs from 0.3-0.8 and hybrid soft-hard clustering power (`clustering_powers`) from 1-9 on both raw and SpQN-corrected correlation matrices. The SpQN-corrected matrix retained approximately 18,000 genes at a PCC cutoff of 0.5 compared to approximately 3,500 before correction, and produced 80-100 modules. We chose to proceed with a PCC cutoff of 0.8 and clustering power of 9 in order to maximize the mean gene margin while retaining approximately 60 modules with 4000 genes. Modules with fewer than 10 genes were excluded from downstream analysis. Modules M12 and M29 were additionally excluded due to sex-linked gene enrichment and low quality, respectively. Module activity scores were computed per spot as follows: for each gene in a module, logcounts expression values were min-max normalized, then the rescaled values were summed across all genes in the module. Module scores were then averaged across spots within each spatial domain and z-scored across domains to enable comparison of domain specificity across modules. Ten modules were identified as domain-specific. Co-expression network visualizations were generated using `ggraph` and `graphlayouts` in R. For improved visualization, within-module edges were down-weighted (x0.75) and between-modules edges were up-weighted (x1.5) to enhance within-module cohesion and between-module separation. Module boundaries were visualized using `geom_mark` from the `ggforce` R package. Finally, the top 5 hub genes per module were labeled based on within-module degree ranking.

### Transcriptional factor detection within spatial networks

To characterize domain-specific modules, transcription factors (TF) were identified using the Lambert et al (2018) human TF census (1,639 total TFs) ^69^. TFs present in the module’s gene list were extracted and their mean logcounts were computed per spatial domain. Expression values were z-scored across domains to identify domain-enriched TFs. TF-centric network visualizations were generated by filtering the Smoothie edge list to train only edge where at least one endpoint was a TF, highlighting TF hub connectivity within each module. Domain-specific marker genes identified using pseudobulk differential expression methods (see above) were also highlighted within these TF hub networks. Finally, Gene Ontology (GO) differential enrichment analysis was performed on domain-specific modules using the compareCluster function from the `clusterProfiler` R/Bioconductor package. Enrichment was computed separately for each biological process (BP), molecular function (MF), and cellular compartment (CC) ontologies across each module. Redundant GO terms were reduced using the `simplify` function. Enrichment results were visualized as heatmaps with the top 5 terms per module.

### Cell type deconvolution

Cell type deconvolution methods were used to both estimate cell type weights in Visium data and perform label transfer to Xenium and VisiumHD datasets. We used robust cell type decomposition (RCTD) ^70^ as implemented in the `spacexr` R package (v2.2.0), using the Yu et al human amygdala snRNA-seq atlas at fine cell type resolution as the reference ^20^. Cell type clusters with less than 25 nuclei were dropped from the snRNA-seq reference to ensure stable expression profiles. The reference was assembled with the `Reference` function from the raw counts matrix, cell type identities, and per-nucleus UMI totals. For all SRT data, raw counts, array coordinates, and per-spot UMI totals were used to construct a `SpatialRNA` object, while dropping any spots with less than 100 UMI. For Visium, RCTD was run in `multi` mode with a maximum number of 5 cell types to be fit per spot to accommodate the mixed cell type content of 55 µm Visium spots. This yielded per-spot cell type weights across all donors. Cell type weights were summed within each spatial domain across donors and then z-scored across domains separately for each cell type, resulting in a spatial domain enrichment score for each cell type.

For Xenium and VisiumHD, RCTD was run in the `doublet` mode to perform label transfer, such that each cell segmentation was assigned a first and second cell type label. The Reference and SpatialRNA objects were built as described above, and cell segmentations with less than 25 total UMIs were dropped. The first predicted cell type label was retained and used to visualize and quantify cell type composition across spatial domains. To resolve the spatial organization of ITC populations using VisiumHD, we mapped the three reproducible ITC subtype labels (see Methods below) back into the Yu et al snRNA-seq data prior to label transfer with RCTD using the `doublet` mode. These cell type labels were spatially visualized to determine which subtypes were island populations.

### Integration and comparison of ITC neuron snRNA-seq datasets

Intercalated cell (ITC) populations were integrated across three human snRNA-seq datasets. Each data was subset to the ITC populations (TSHZ1 labeled populations in Yu et al and Totty et al, and the eccentric medium spiny neuron population in Siletti et al) and joined into one SingleCellExperiment object. The combined object was log-normalized and highly variable genes were selected with modelGeneVar and get TopHVGs (top 2,000, blocked by dataset). The top 30 PCA components were then integrated across both dataset and sample with Harmony, and UMAP was run on the corrected embeddings. MetaNeighbor was used to assess the replicability of ITC cell types found across datasets. Using highly variables gene selected per study by the variableGenes function, unsupervised MetaNeighbor (MetaNeighborUS, fast_version=TRUE) computed pairwise AUROC scores between cell types across datasets at the fine ITC-type level, and the results were visualized as cluster AUROC heatmaps. Based on the MetaNeighbor fine-type dendrogram, Siletti eti al ITC fine clusters from all three datasets were grouped into three cross-dataset subtypes (ITC_1, ITC_2, ITC_3). Subtype marker genes were identified with the findMarkers from scran, retaining the top 50 per subtype, and characterized with marker heatmaps, canonical spiny/aspiny gene panels, and neurotransmitter-receptor dot plots (glutamate, GABA, catecholamine receptor families).

### Spatial mapping and colocalization analysis of ITC subtypes

ITC subtypes labels were transferred to Yu et al snRNs-seq prior to cell type mapping in the VisiumHD dataset. To transfer the three subtype labels from Siletti et al to Yu et ali, we performed reference mapping with Symphony. In short, query cells from Yu et al were mapped to the Siletti et al reference using k=25 nearest neighbor prediction. Label transfer was then performed from the Yu et al dataset to the VisiumHD datasets using RCTF as described above. Spatial relationships among ITC subtypes and other cell types were assessed using hoodscanR after removing low-confidence assignments (RCTD predicted weight <0.65). For each sample, neighborhood composition was computed over the k=100 nearest neighbors, and colocalization was quantified as the Pearson correlation between cell types’ neighborhood-probability vectors across the pooled matrix, visualized as clusters heatmaps. Local density of each ITC subtype was additionally computed with point-pattern analysis via statspat. Each sample’s marked point patterns were built from cell coordinates, and neighbor counts within 5x the median nearest-neighbor distance (closepairs) were summarized for each subtype.

### Cell type mapping of PuV, ASt, and CeA domains

For spatial mapping of putative CeA cell types of the Yu et al dataset (*DRD2+/ISL1+, DRD+/PAX6+,* and *PRKCD+*) in standard Visium, we used the used the resulting weights from RCTD cell type deconvolution described above. Spots were assigned to the subdomain (PuV, ASt, and CeA) of their highest-weighted defining population. To characterize the topography of these subdomains, per-sample spot proportions of each subdomain were computed as a fraction of all spots and plotted across the four ordered anterior-posterior levels. Subdomain assignments were confirmed against the spatial expression of their top marker genes, found using findMarkers from the scran package.

### 4.5 Spatially-aware heritability enrichment with gsMap

#### Spatially-aware heritability enrichment (gsMap)

We mapped GWAS heritability onto amygdala spatial domains using gsMap v1.73.7 (Python 3.11.15) with 21 different traits ^71–88^. All seven Visium donor sections were analyzed (224,021 spots total; 19,389 protein-coding genes), with spots assigned to 15 spatial domains by the BS_k16_Semisupervised_wAI annotation. Latent representations were learned per section from raw counts, and gene specificity scores were computed with gene ranks placed on a common scale across sections (--gM_slices) so that domain-level estimates are comparable between donors.

Rather than gsMap’s default baseline, which conditions on a single gene-proximity annotation, we conditioned all analyses on the full 52-annotation baselineLD functional model. Stratified LD scores were generated per donor and chromosome against the 1000 Genomes EUR Phase 3 reference panel, restricted to HapMap3 SNPs, using a 50 kb window around each gene body (GENCODE v46lift37). Spatial LD-score regression was then run per donor with the additional baseline enabled, yielding a heritability enrichment *P* value at every spot. Reported enrichments therefore reflect signals beyond generic functional and gene-proximity effects; without this conditioning, enrichment *P* values are inflated by roughly four orders of magnitude and the non-brain control traits reach significance spuriously.

Per-spot *P* values were combined within each domain and donor by the Cauchy combination test, then across all seven donors, giving one P value per domain per trait. Domains absent or sparsely sampled in some donors (CLA 1/7, LA 6/7, AI 6/7 sections) average over fewer sections accordingly.

### 4.6 Spatial enrichment of disease-associated genes with scDRS

#### Gene-level association statistics

GWAS summary statistics from the same 21 traits were converted to SNP-level p-values, using the effective sample size where available (N, else NEFF, else 2 × NEFF_HALF, else 4/(1/N_case + 1/N_control)). Gene-level statistics were computed with MAGMA v1.10 using the SNP-wise mean model, a 10 kb upstream / 1.5 kb downstream window, NCBI37.3 gene coordinates, and the 1000 Genomes Phase 3 European reference panel (489 individuals, 9,997,231 variants). Per-trait disease gene sets were built with scDRS munge-gs, retaining the top 1,000 genes by MAGMA z-score with weights clipped at 10.

#### scDRS

Spot-level disease scores were computed with scDRS v1.0.2 (scanpy 1.11.5, numpy 2.4.6) using 1,000 control gene sets matched on mean expression and expression variance. Covariates were total UMI (log), number of detected genes, and donor (six dummy variables, Br2743 as reference); covariate correction was applied implicitly on the sparse count matrix. Sparse spots were removed by scDRS’s internal filter, leaving 213,127 of 224,021 spots scored per trait.

Domain-level enrichment was assessed with scDRS perform-downstream, which returns a Monte Carlo association statistic (assoc_mcz) and p-value (assoc_mcp, bounded below by 1/1001) per domain × trait pair. 42 traits were scored. Multiple-testing correction was applied once across the full as-run family of 630 tests (15 domains × 42 traits), yielding 38 significant pairs at FDR < 0.05. Height served as a negative control. Spatial organization of each score was quantified two ways: domain spread (the range of domain-mean normalised scores) and Moran’s I on a k = 6 nearest-neighbour graph built within capture areas, using the merged full-resolution pixel coordinates.

### 4.7 Single nucleus RNA-seq datasets

#### Whole amygdala snRNA-seq reference atlas

Single-nucleus transcriptomic data of the whole human amygdala were obtained from Yu et al. ^20^, from which we used only the human subset. This dataset consists of 91,699 high-quality human nuclei profiled by 10x Genomics Chromium v3 platform from three postmortem donors. We downloaded the processed Seurat/R object directly from the Gene Expression Omnibus (GEO SE195445). We used this dataset as the primary reference for cell type mapping as it is currently the most complete snRNA-seq atlas of the human amygdala.

#### Amygdalar intercalated neurons snRNA-seq subset

Single-nucleus transcriptomic data of ITCs were extracted from the Siletti et al. whole-human-brain (WHB) snRNA-seq atlas, accessed through the Allen Brain Cell (ABC) atlas. Nuclei collected from amygdala dissections were collected from the atlas and read into R, yielding 66,072 nuclei and 59,357 genes across 4 donors. Nuclei from the eccentric medium spiny neuron (EMSN) supercluster (39,276) were then subset, producing the ITC object that was used for downstream analyses.

## 4.8 Data and code availability

All analysis code and final `SpatialExperiment` R objects used for analysis can be openly accessed on github (https://github.com/LieberInstitute/spatialAmygdala/tree/devel). FASTQ raw data, as well as `CellRanger` and `SpaceRanger` output files, can be found through GEO accessions for each Visium (GSE342709), Xenium (GSE342289), and VisiumHD (GSE342738) datasets. SRT data can be interactively accessed and viewed through a series of Vitessce ^89^ apps, found on the project github page. Any additional information required to reanalyze the data reported in this paper is available from the lead contact upon request.

## 4.9 Acknowledgements, Funding, Authorship Contributions

## Acknowledgements

We gratefully thank the families who donated tissue to make this research possible. We thank the Essel Foundation and the families of Connie and Stephen Lieber and Milton and Tamar Maltz for their generous support of this work. We would like to acknowledge the contributions of Dr. Fernando Goes, the late Dr. Llewellyn B. Bigelow, Amy Deep-Soboslay, Anna Brandtjen, and James Tooke for sample curation and clinical characterization. We also thank the Office of the Chief Medical Examiner of the State of Maryland, the Department of Pathology at Western Michigan University Homer Stryker MD School of Medicine, the University of North Dakota School of Medicine and Health Sciences, Department of Pathology, and Gift of Life Michigan for their collaboration on tissue collection. We thank members of the Martinowich, Maynard, Page, and Hicks labs for critical reading of the manuscript and feedback. We thank the Joint High Performance Computing Exchange (JHPCE) for providing computing resources for these analyses. Portions of some figures were created with BioRender.com.

## Funding

This project was supported by R01DA053581 (KM) and the Lieber Institute for Brain Development. MST was supported by F32MH123620.

## Conflict of Interest

The authors declare no competing interests.

## Author Contributions

Conceived and Designed the Study: MST, KM

Performed Experiments and Collected Data: SVB, MRV, SEM, IDRA

Software: MST, RAM

Formal Analysis: MST

Data Curation: MST, MT, RAM

Tissue Resources: SVB, JEK, TMH

Writing – original draft: MST, SVB, MT

Writing – review & editing: MST, SVB, MT, SCP, SCH, KM

Visualization: MST, MT

Supervision: KRM, SCP, TMH, SCH, KM

Funding acquisition: MST, KRM, SCH, KM

Project Administration: SCH, KM

## Notes

### Competing Interest Statement

The authors have declared no competing interest.

