## Supplementary Figures for "Multiscale spatial transcriptomics resolves the cellular and molecular architecture of the human amygdala"

**Title**

### Supplementary Figures

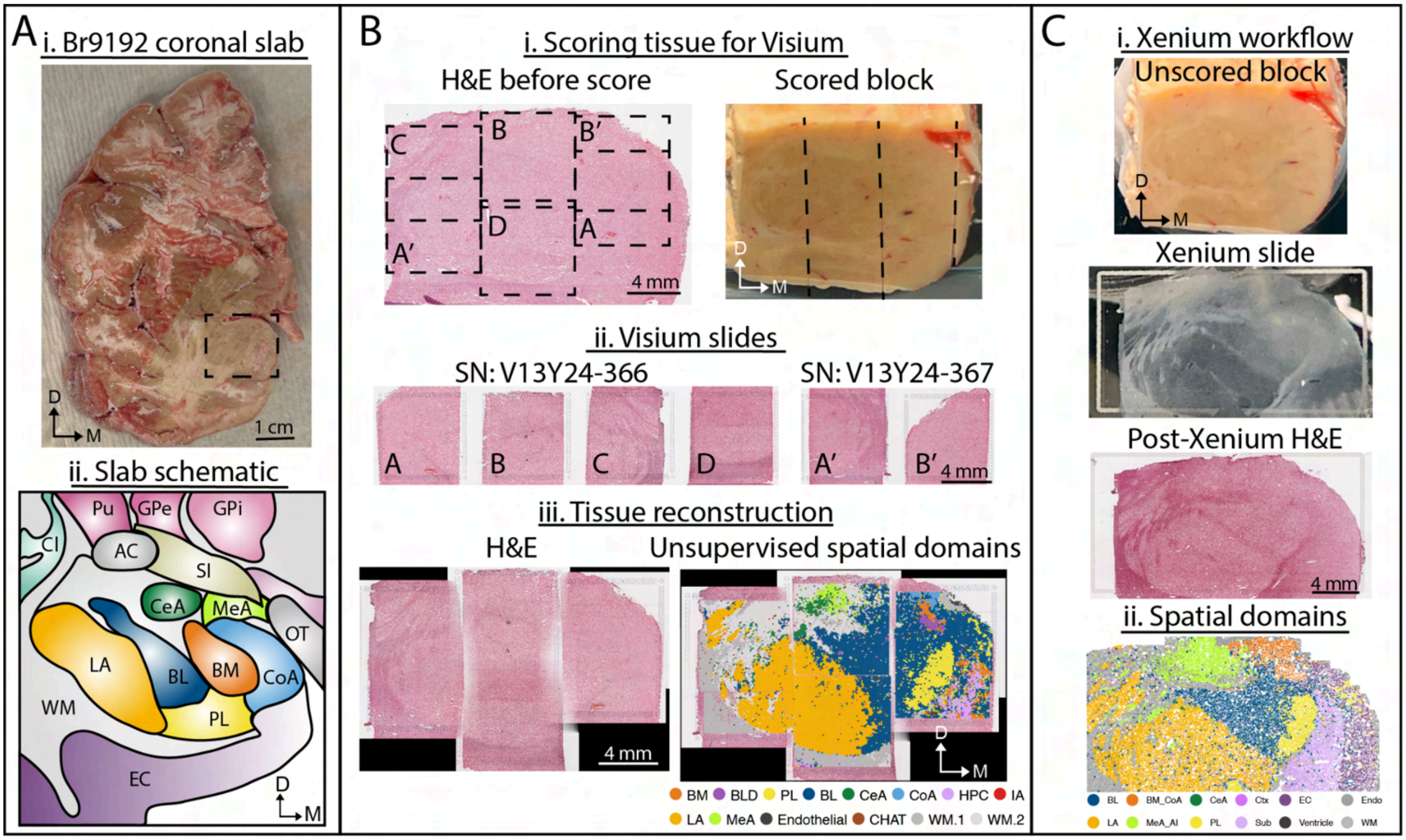


**Supplementary Fig 1. Anatomical orientation of anterior-intermediate AMY block from donor Br9192. (A)** Fresh frozen coronal slab of donor Br9192 at the level of anterior-intermediate amygdala (i) and a schematic representation of AMY subnuclei at the corresponding A-P level (ii). Dashed box denotes dissected block boundaries. Scale bar 1 cm. **(B)** Visium workflow illustrating H&E stained AMY tissue section and the brain block scored in 6.5 mm stripes to fit the dimensions of the Visium capture arrays (i); H&E stain of scored AMY tissue on 6 Visium arrays across 2 slides (ii); H&E stained AMY tissue after stitching the capture arrays and the overlaid unsupervised spatial domains (iii). A-D, A', B' represent different Visium capture arrays. Scale bars 4 mm. **(C)** Xenium workflow illustrating the unscored AMY tissue block (i), AMY tissue slice on the Xenium slide (i), post-Xenium H&E stain on the Xenium slide (i), and the spatial domain clusters (ii). Scale bars 4 mm.

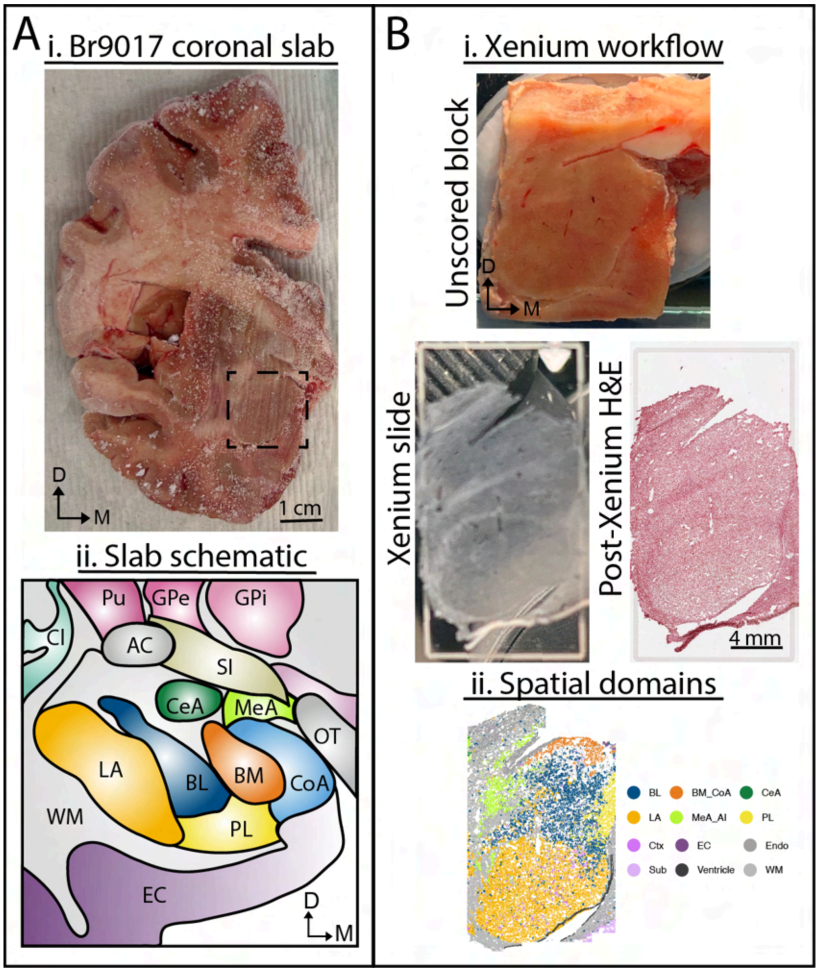


**Supplementary Fig 2. Anatomical orientation of anterior-intermediate amygdala block from donor Br9017. (A)** Fresh frozen coronal slab of donor Br9017 at the level of anterior-intermediate amygdala (i) and a schematic representation of AMY subnuclei at the corresponding A-P level (ii). Dashed box denotes dissected block boundaries. Scale bar 1 cm. **(B)** Xenium workflow illustrating the unscored AMY tissue block (i), AMY tissue slice on the Xenium slide (i), post-Xenium H&E stain on the Xenium slide (i), and the spatial domain clusters (ii). Scale bars 4 mm.

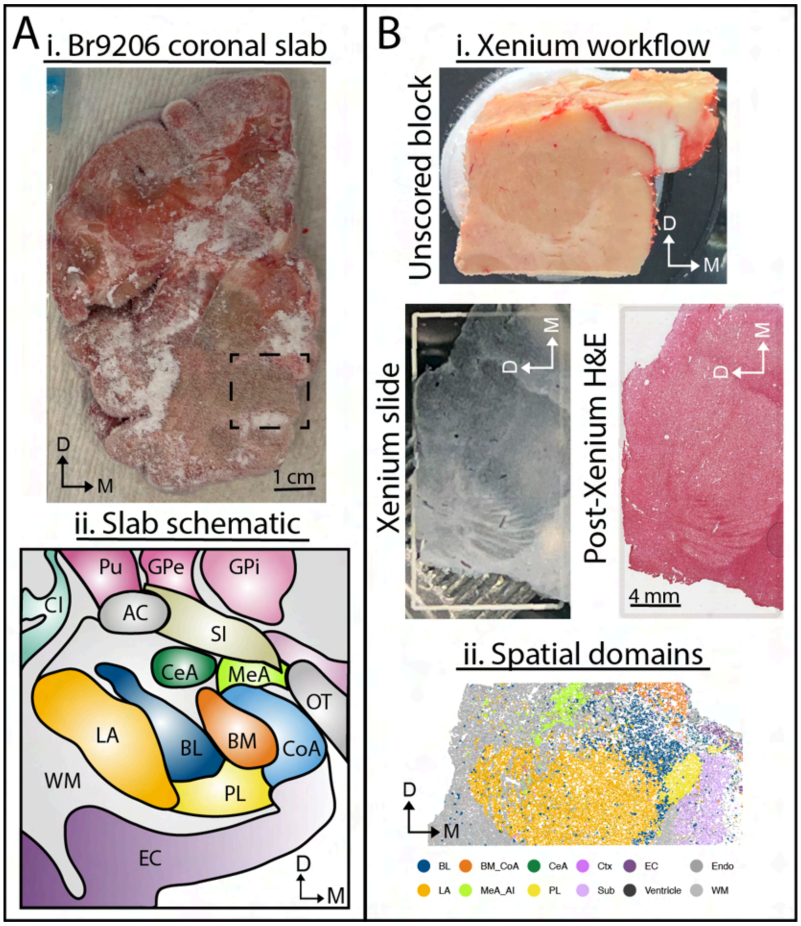


**Supplementary Fig 3. Anatomical orientation of anterior-intermediate amygdala block from donor Br9206. (A)** Fresh frozen coronal slab of donor Br9206 at the level of anterior-intermediate amygdala (i) and a schematic representation of AMY subnuclei at the corresponding A-P level (ii). Dashed box denotes dissected block boundaries. Scale bar 1 cm. **(B)** Xenium workflow illustrating the unscored AMY tissue block (i), AMY tissue slice on the Xenium slide (90° rotation) (i), post-Xenium H&E stain on the Xenium slide (90° rotation) (i), and the spatial domain clusters (ii). Scale bars 4 mm.

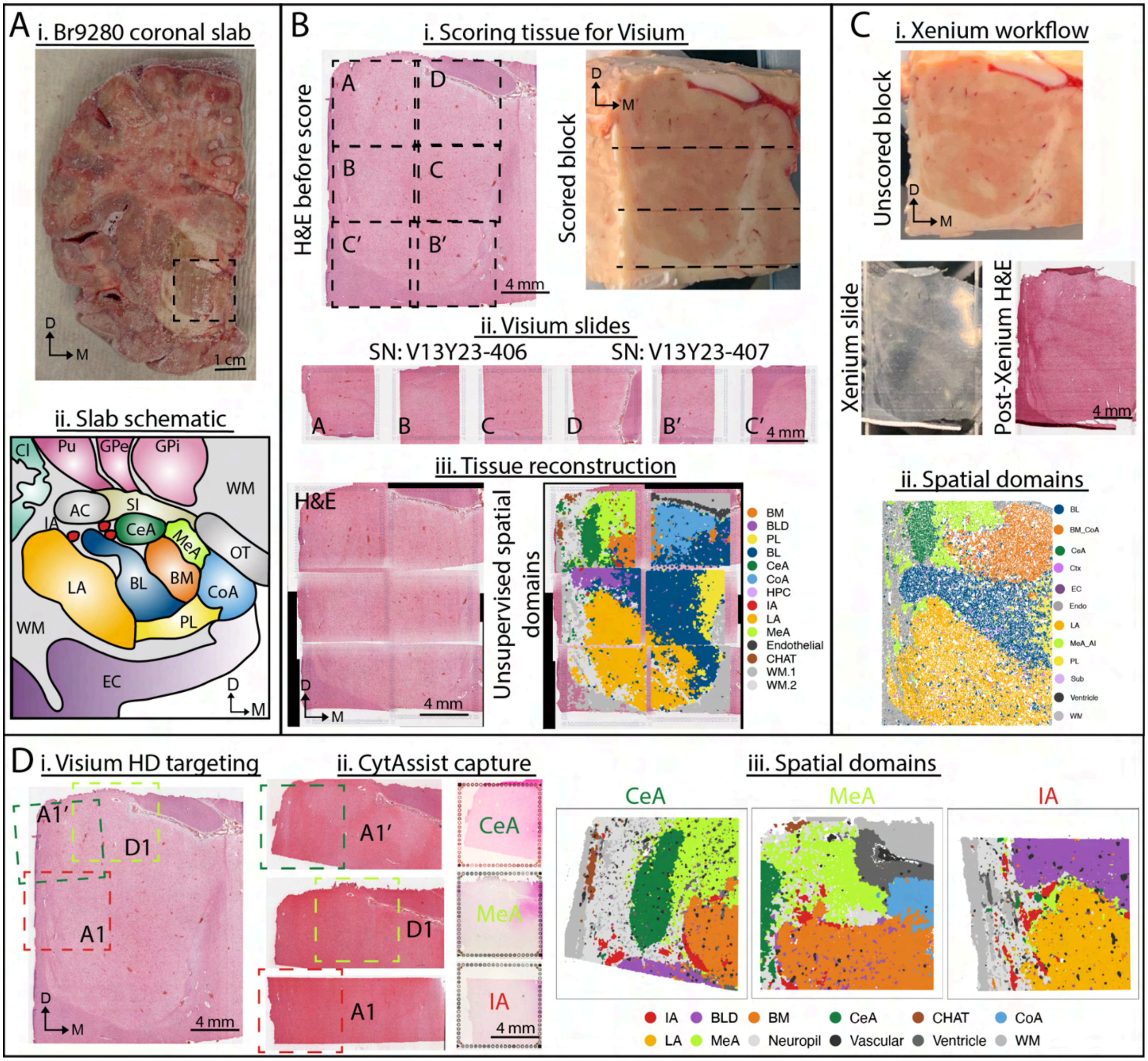


**Supplementary Fig 4. Anatomical orientation of intermediate amygdala block from donor Br9280. (A)** Fresh frozen coronal slab of donor Br9280 at the level of intermediate amygdala (i) and a schematic representation of AMY subnuclei at the corresponding A-P level (ii). Dashed box denotes dissected block boundaries. Scale bar 1 cm. **(B)** Visium workflow illustrating H&E stained AMY tissue section and the brain block scored in 6.5 mm stripes to fit the dimensions of the Visium capture arrays (i); H&E stain of scored AMY tissue on 6 Visium arrays across 2 slides (ii); H&E stained AMY tissue after stitching the capture arrays and the overlaid unsupervised spatial domains (iii). A-D, B', C' represent different Visium capture arrays. Scale bars 4 mm. **(C)** Xenium workflow illustrating the unscored AMY tissue block (i), AMY tissue slice on the Xenium slide (i), post-Xenium H&E stain on the Xenium slide (i), and the spatial domain clusters (ii). Scale bars 4 mm. **(D)** Visium HD workflow illustrating H&E stained AMY tissue section (i), scored AMY tissue used for CytAssist before and after capture (ii), and resulting spatial domains targeted to capture the central amygdala (CeA), the medial amygdala (MeA), and the intercalated amygdalar islands (IA) (iii). A1, D1, A1' represent different VisiumHD capture arrays. Scale bars 4 mm.

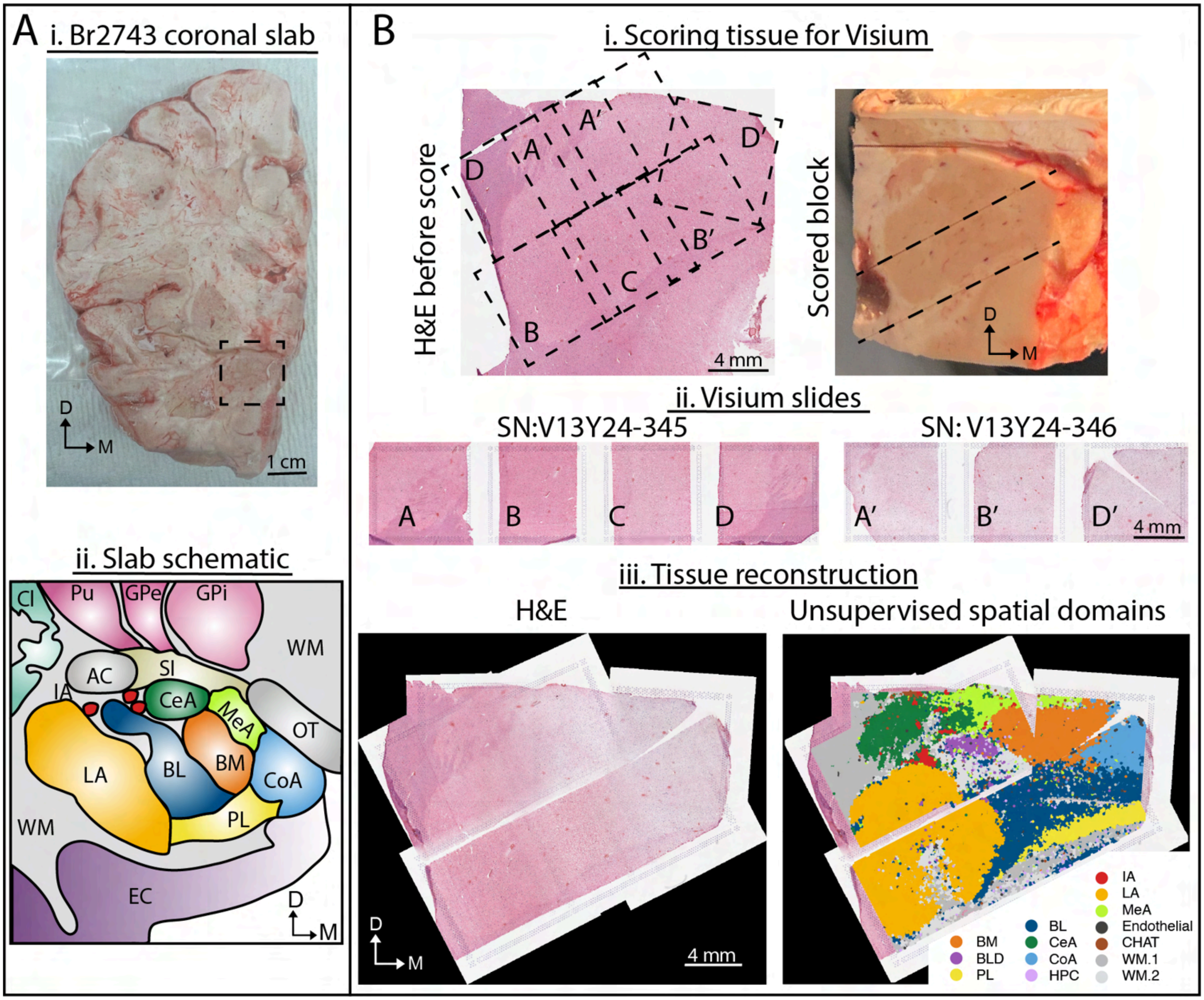


**Supplementary Fig 5. Anatomical orientation of intermediate amygdala block from donor Br2743. (A)** Fresh frozen coronal slab of donor Br2743 at the level of intermediate amygdala (i) and a schematic representation of AMY subnuclei at the corresponding A-P level (ii). Dashed box denotes dissected block boundaries. Scale bar 1 cm. **(B)** Visium workflow illustrating H&E stained AMY tissue section and the brain block scored in 6.5 mm stripes to fit the dimensions of the Visium capture arrays (i); H&E stain of scored AMY tissue on 7 Visium arrays across 2 slides (ii); H&E stained AMY tissue after stitching the capture arrays and the overlaid unsupervised spatial domains (iii). A-D, A', B', D' represent different Visium capture arrays. Scale bars 4 mm.

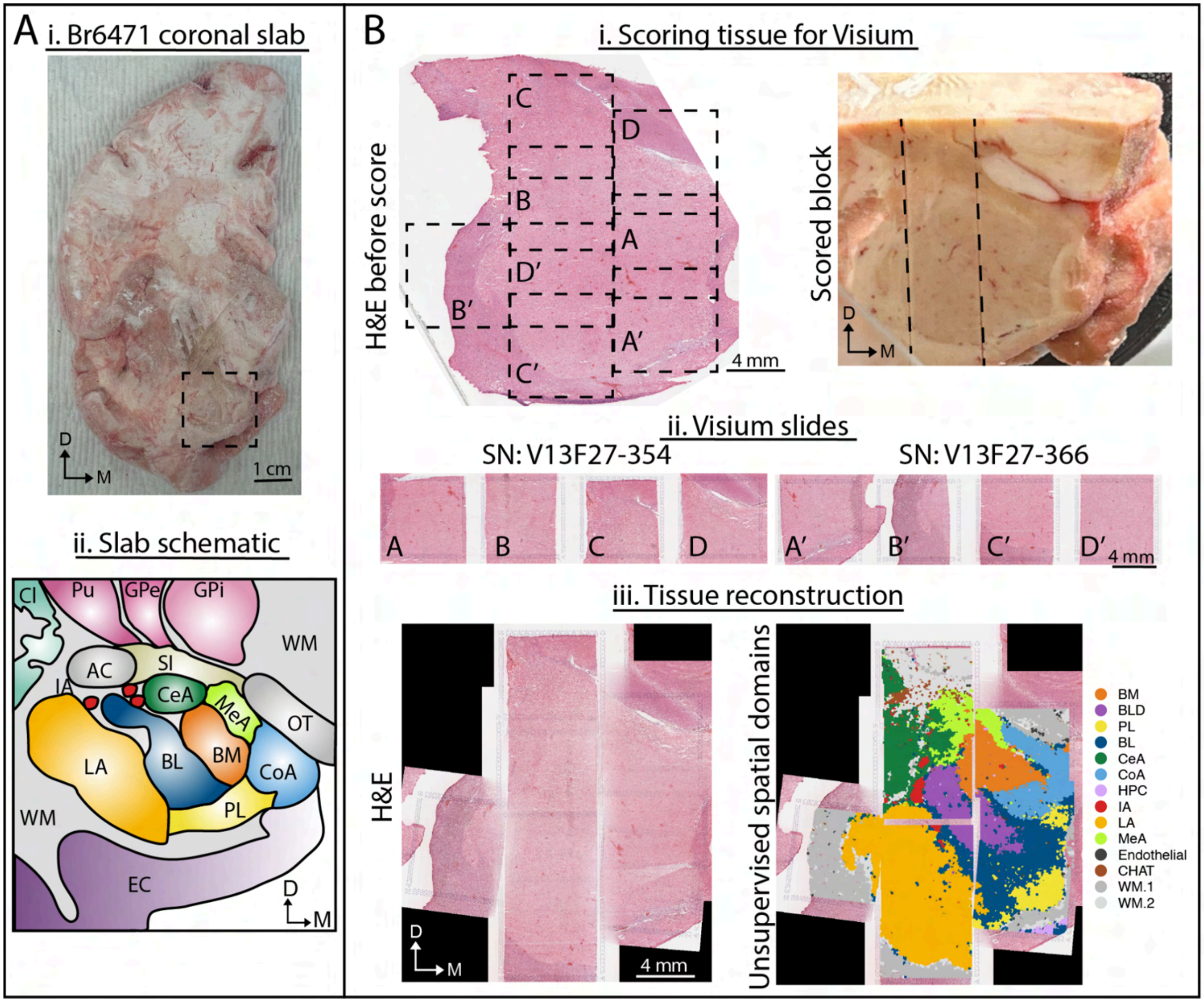


**Supplementary Fig 6. Anatomical orientation of intermediate amygdala block from donor Br6471. (A)** Fresh frozen coronal slab of donor Br6471 at the level of intermediate amygdala (i) and a schematic representation of AMY subnuclei at the corresponding A-P level (ii). Dashed box denotes dissected block boundaries. Scale bar 1 cm. **(B)** Visium workflow illustrating H&E stained AMY tissue section and the brain block scored in 6.5 mm stripes to fit the dimensions of the Visium capture arrays (i); H&E stain of scored AMY tissue on 8 Visium arrays across 2 slides (ii); H&E stained AMY tissue after stitching the capture arrays and the overlaid unsupervised spatial domains (iii). A-D, A'-D' represent different Visium capture arrays. Scale bars 4 mm.

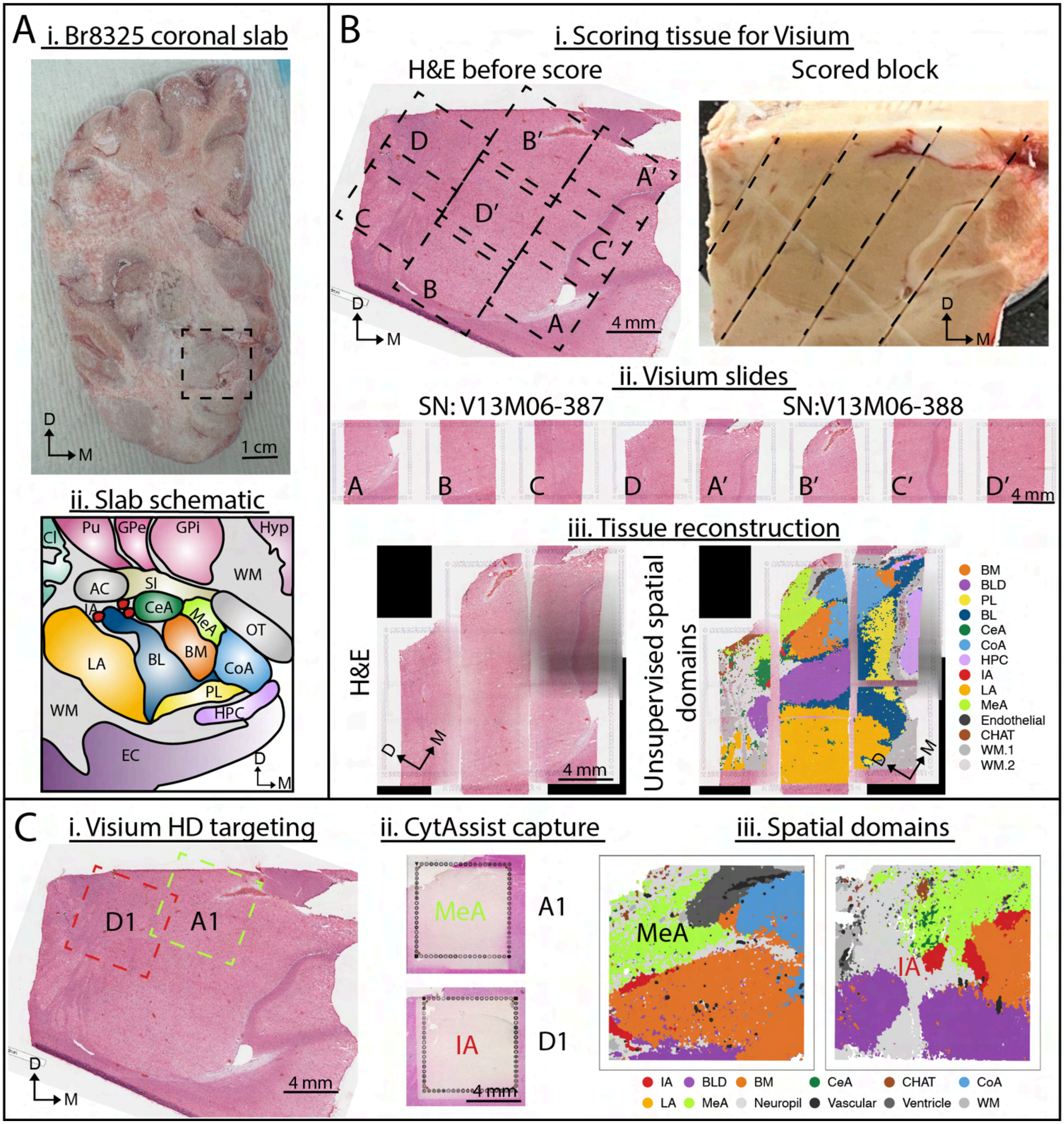


**Supplementary Fig 7. Anatomical orientation of posterior-intermediate amygdala block from donor Br8325. (A)** Fresh frozen coronal slab of donor Br8325 at the level of posterior-intermediate amygdala (i) and a schematic representation of AMY subnuclei at the corresponding A-P level (ii). Dashed box denotes dissected block boundaries. Scale bar 1 cm. **(B)** Visium workflow illustrating H&E stained AMY tissue section and the brain block scored in 6.5 mm stripes to fit the dimensions of the Visium capture arrays (i); H&E stain of scored AMY tissue on 8 Visium arrays across 2 slides (ii); H&E stained AMY tissue after stitching the capture arrays and the overlaid unsupervised spatial domains (iii). A-D, A'-D' represent different Visium capture arrays. Scale bars 4 mm. **(C)** Visium HD workflow illustrating H&E stained AMY tissue section (i), scored AMY tissue used for CytAssist after capture (ii), and resulting spatial domains targeted to capture the medial amygdala (MeA)

and the intercalated amygdalar islands (IA) (iii). A1, D1, A1’ represent different VisiumHD capture arrays. Scale bars 4 mm.

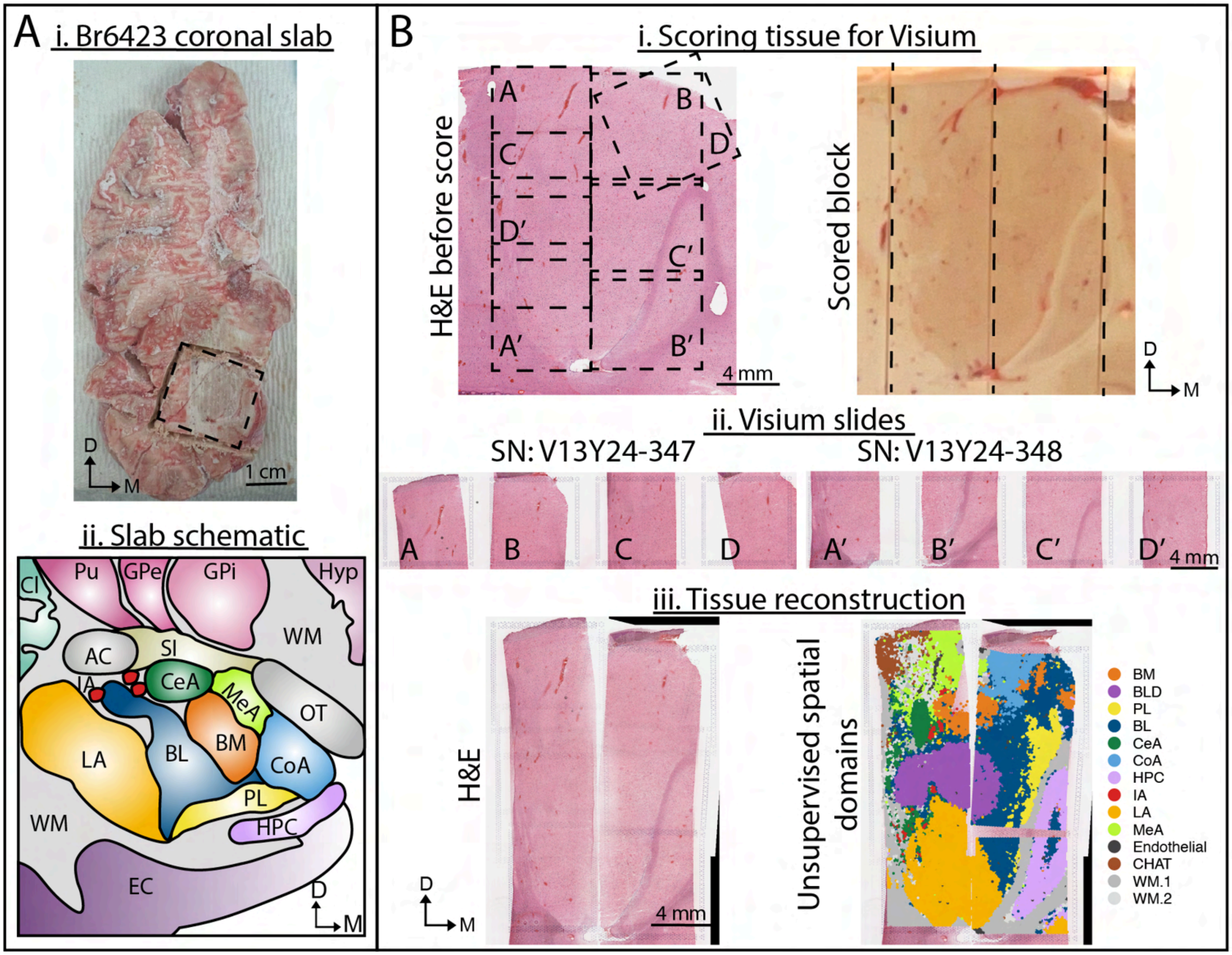


**Supplementary Fig 8. Anatomical orientation of posterior-intermediate amygdala block from donor Br6423. (A)** Fresh frozen coronal slab of donor Br6423 at the level of posterior-intermediate amygdala (i) and a schematic representation of AMY subnuclei at the corresponding A-P level (ii). Dashed box denotes dissected block boundaries. Scale bar 1 cm. **(B)** Visium workflow illustrating H&E stained AMY tissue section and the brain block scored in 6.5 mm stripes to fit the dimensions of the Visium capture arrays (i); H&E stain of scored AMY tissue on 8 Visium arrays across 2 slides (ii); H&E stained AMY tissue after stitching the capture arrays and the overlaid unsupervised spatial domains (iii). A-D, A’-D’ represent different Visium capture arrays. Scale bars 4 mm.

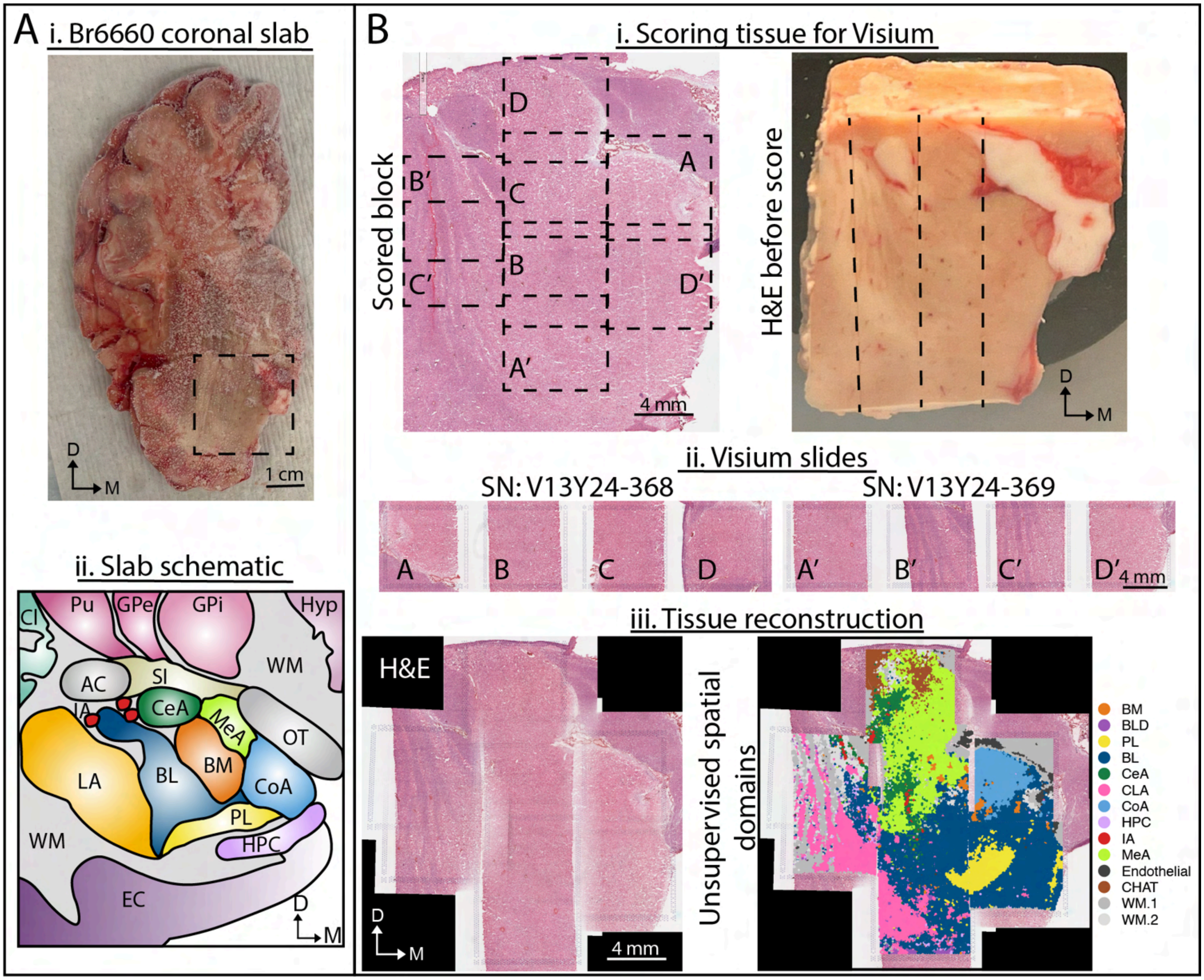


**Supplementary Fig 9. Anatomical orientation of posterior amygdala block from donor Br6660. (A)** Fresh frozen coronal slab of donor Br6660 at the level of posterior amygdala (i) and a schematic representation of AMY subnuclei at the corresponding A-P level (ii). Dashed box denotes dissected block boundaries. Scale bar 1 cm. **(B)** Visium workflow illustrating H&E stained AMY tissue section and the brain block scored in 6.5 mm stripes to fit the dimensions of the Visium capture arrays (i); H&E stain of scored AMY tissue on 8 Visium arrays across 2 slides (ii); H&E stained AMY tissue after stitching the capture arrays and the overlaid unsupervised spatial domains (iii). A-D, A'-D' represent different Visium capture arrays. Scale bars 4 mm.

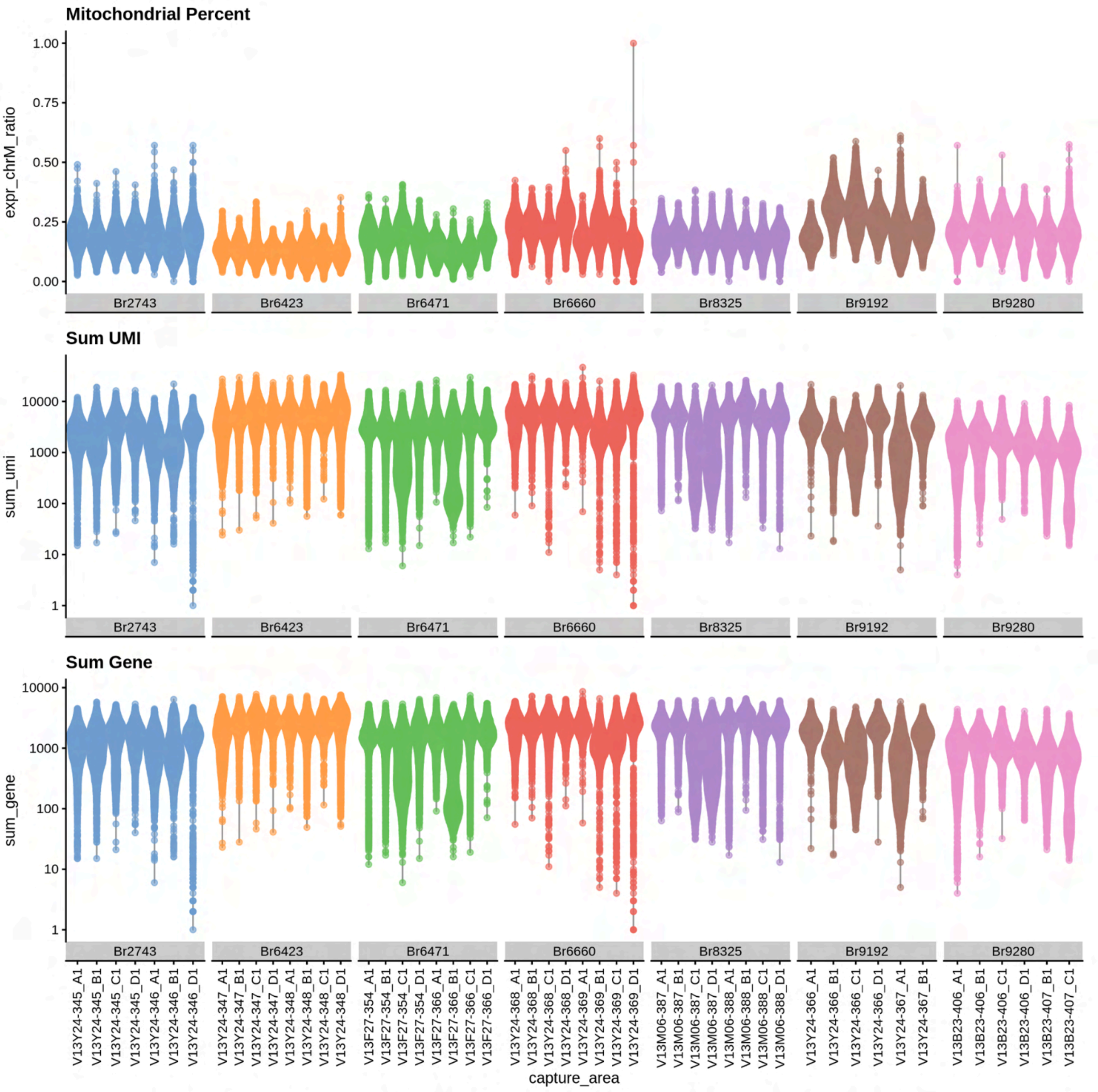


**Supplementary Fig 10. Visium quality control metrics across donors and capture areas.** Violin plots show (top) mitochondrial read fraction (chrM ratio), (middle) total UMIs per spot, and (bottom) detected genes per spot, stratified by donor and capture area within each donor. Dashed red lines indicate QC thresholds used to flag low-quality spots; points represent individual spots, and UMI/gene counts are shown on a log scale.

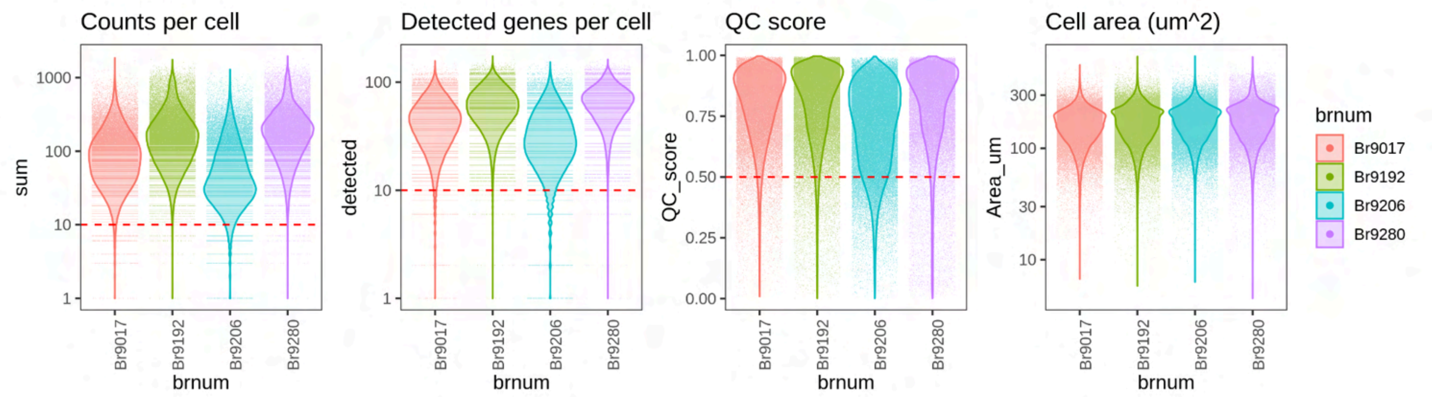


**Supplementary Fig 11. Xenium quality control metrics across donors.** Violin plots show sum of total mRNA counts, number of unique detected genes, the SpaceTrooper quality control (QC) scores, and cell area per segmented cell. Dashed red lines indicate QC thresholds used to flag low-quality spots.

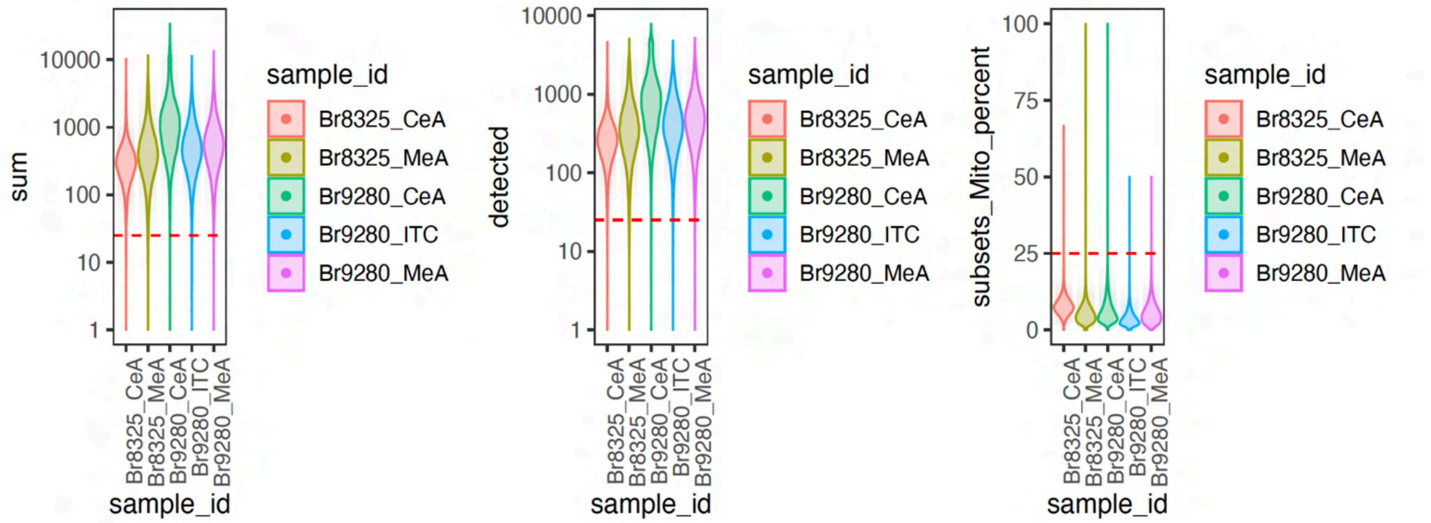


**Supplementary Fig 12. VisiumHD quality control metrics across donors and capture areas.** Violin plots show (left) sum of total mRNA transcripts, (middle) number of unique detected genes, and (right) percent of mRNA transcript mapping to mitochondrial genome per segmented cell. Dashed red lines indicate QC thresholds used to flag low-quality spots.

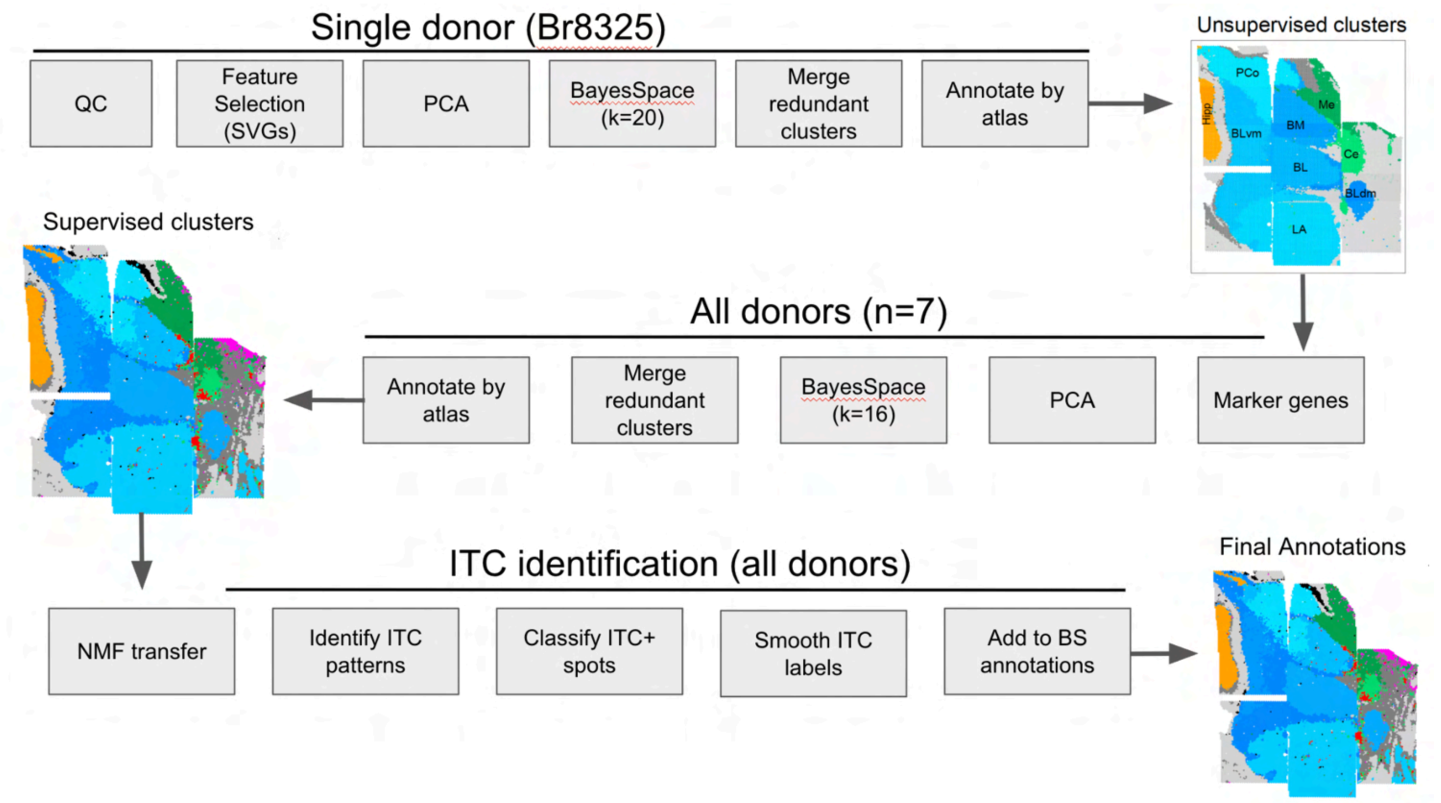


**Supplementary Fig 13. Spatial domain annotation strategy for human amygdala Visium data.** Schematic of the clustering and annotation workflow used to define spatial domains. Initial clustering was performed in a single representative donor (Br8325) using spatially variable genes, PCA, and BayesSpace, followed by merging of redundant clusters and atlas-guided anatomical annotation. Marker genes identified from these initial domains were then used to guide clustering across all seven donors via BayesSpace clustering, cluster merging, and atlas-based annotation. Intercalated islands (ITCs) were identified separately by transferring NMF-derived expression patterns, classifying ITC-positive spots, and spatially smoothing ITC labels before incorporation into the final spatial domain annotations.

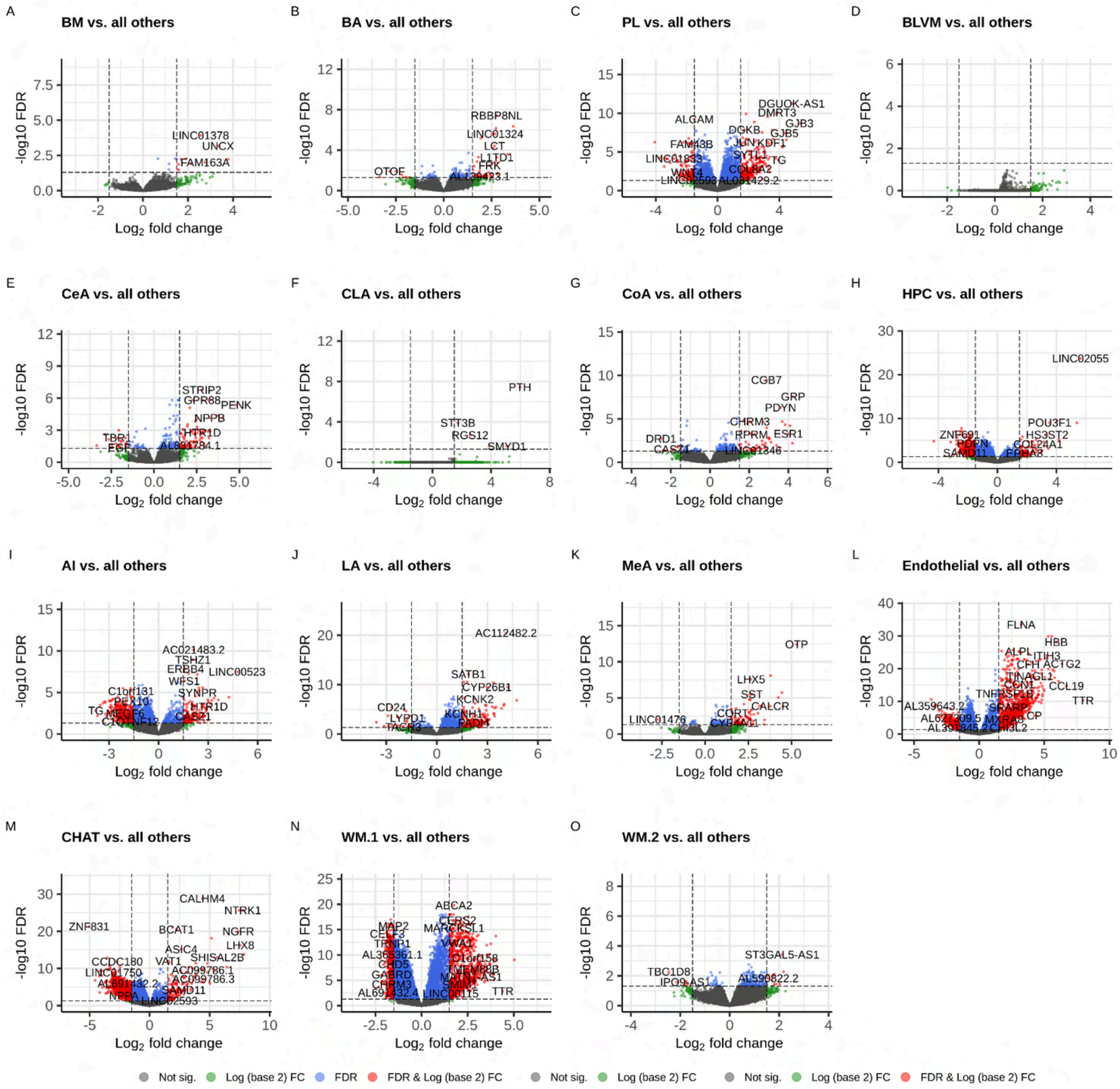


**Supplementary Fig 14. One-vs-all pseudobulk differential expression identifies marker genes for Visium spatial domains. (A–O)** Volcano plots show log2 fold change (x-axis) versus −log10 FDR (y-axis) for pseudobulk differential expression comparing each spatial domain to all other domains. Dashed vertical and horizontal lines indicate fold-change and FDR significance thresholds, respectively. Points are colored by significance (FDR), effect size (log2FC|), or both; selected top hits are labeled.

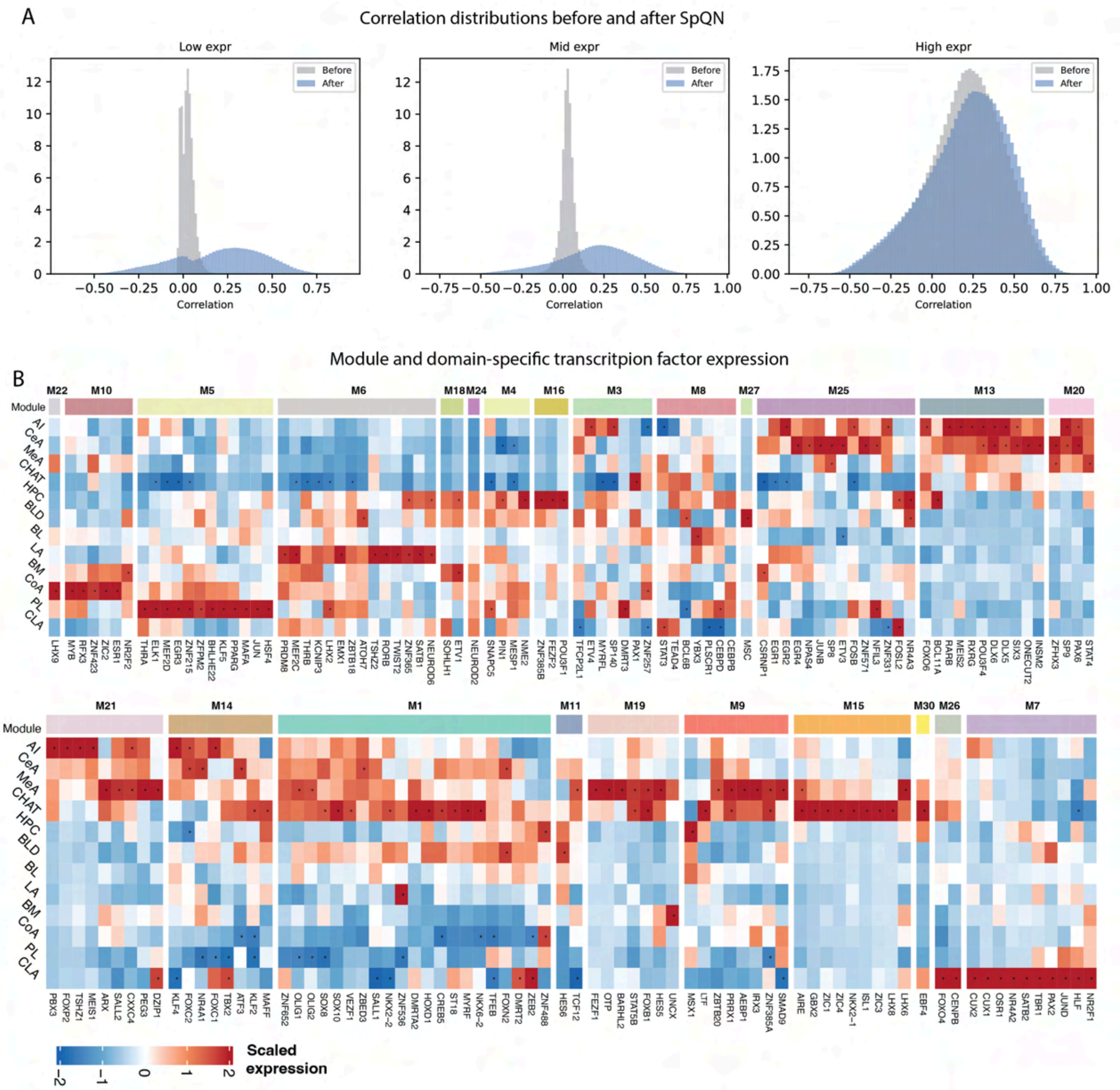


**Supplementary Fig 15. SpQN normalization and module-specific transcription factor expression across spatial domains.** **(A)** Distribution of pairwise gene–gene correlations before (grey) and after (blue) spatial quantile normalization (SpQN), stratified by mean expression level into low-, mid-, and high-expression bins. Prior to normalization, correlations among low- and mid-expressed genes are sharply concentrated near zero relative to high-expressed genes, reflecting mean–correlation bias. After normalization, the three bins show comparable correlation distributions, indicating that co-expression strength is no longer confounded by expression magnitude. **(B)** Scaled mean expression (z-score) of transcription factors assigned to each co-expression module across spatial domains. Columns are grouped by module (top annotation bar, colored as in Fig. 3); rows are spatial domains. Related to Fig. 3E, which shows the subset of modules with domain-specific enrichment.

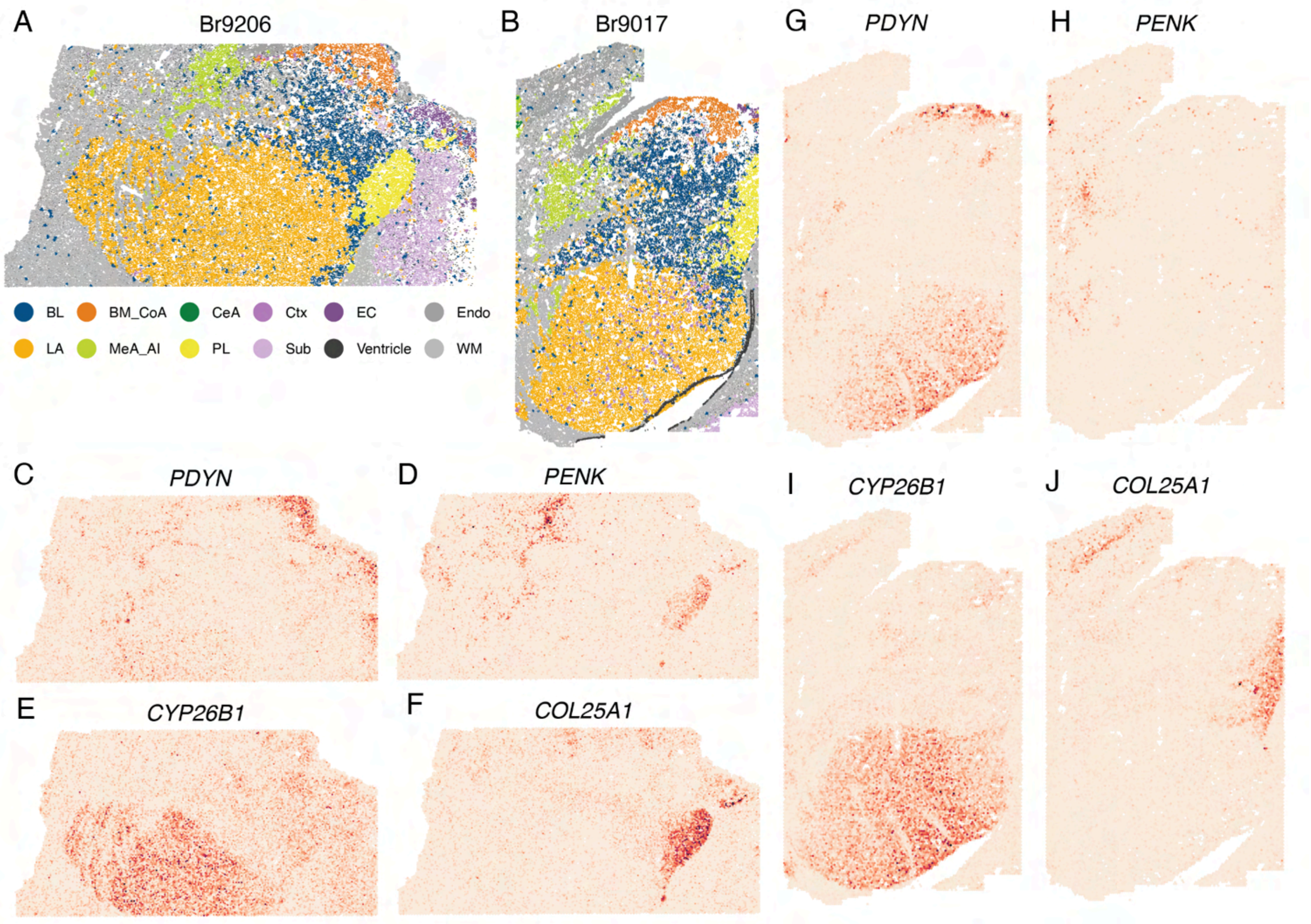


**Supplementary Fig 16. Xenium spatial domains and marker gene expression in donors Br9206 and Br9017. (A–B)** BANKSY-defined spatial domains in Xenium sections from donors Br9206 (A) and Br9017 (B). Each point is a segmented cell, colored by domain assignment: BL, BM_CoA, CeA, LA, MeA_IA, PL, cortex (Ctx), entorhinal cortex (EC), subiculum (Sub), ventricle, endothelium (Endo), and white matter (WM). Domain colors match those in Fig. 4. **(C–F)** Rasterized (~25 μm bins) spatial expression of *PDYN* (C), *PENK* (D), *CYP26B1* (E), and *COL25A1* (F) in the Br9206 section shown in (A). **(G–J)** As in (C–F), for the Br9017 section shown in (B).

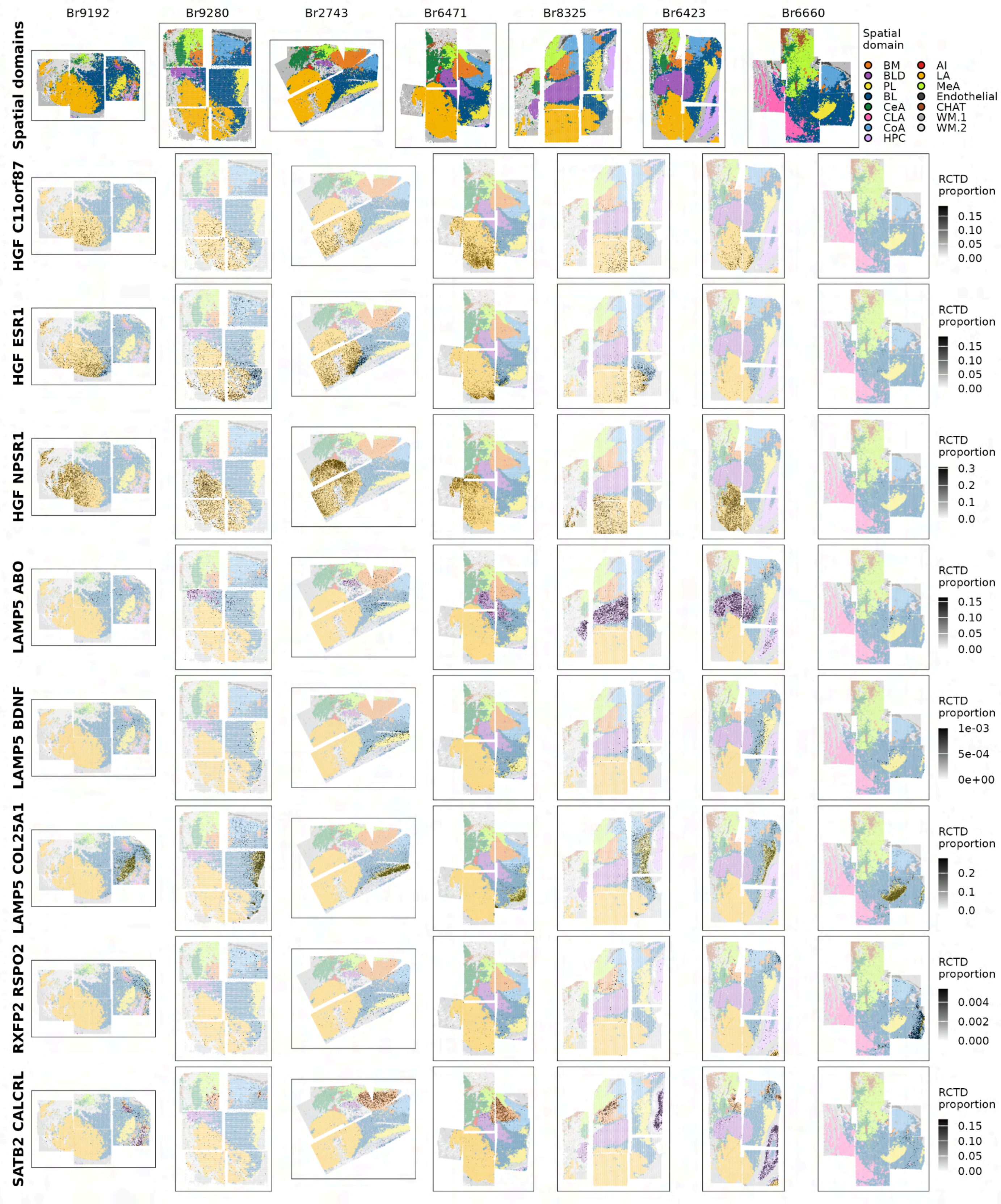


**Supplementary Fig 17. Spatial mapping of neuronal cell types across the human amygdala using RCTD.** Spatial distributions of neuronal cell types mapped onto Visium spatial transcriptomic data using RCTD with the human amygdala single-nucleus RNA-seq dataset from Yu et al. as a reference. Columns show individual Visium donors and rows show representative neuronal cell types. The top row displays spatial domain annotations for anatomical reference. For each cell type, grayscale intensity indicates the RCTD-estimated proportion within each Visium spot, overlaid on spatial domain annotations.

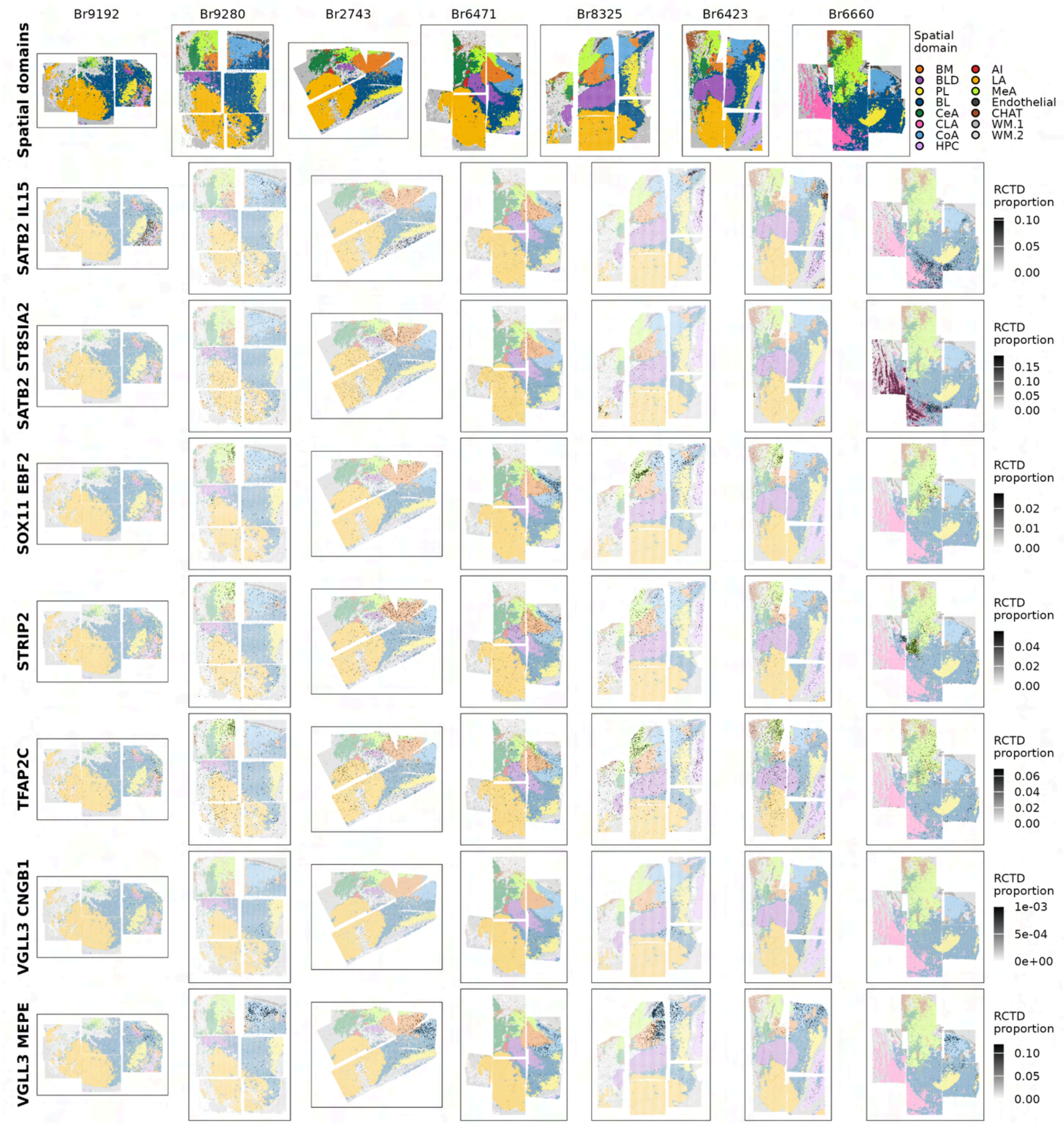


**Supplementary Fig 17 - continued.**

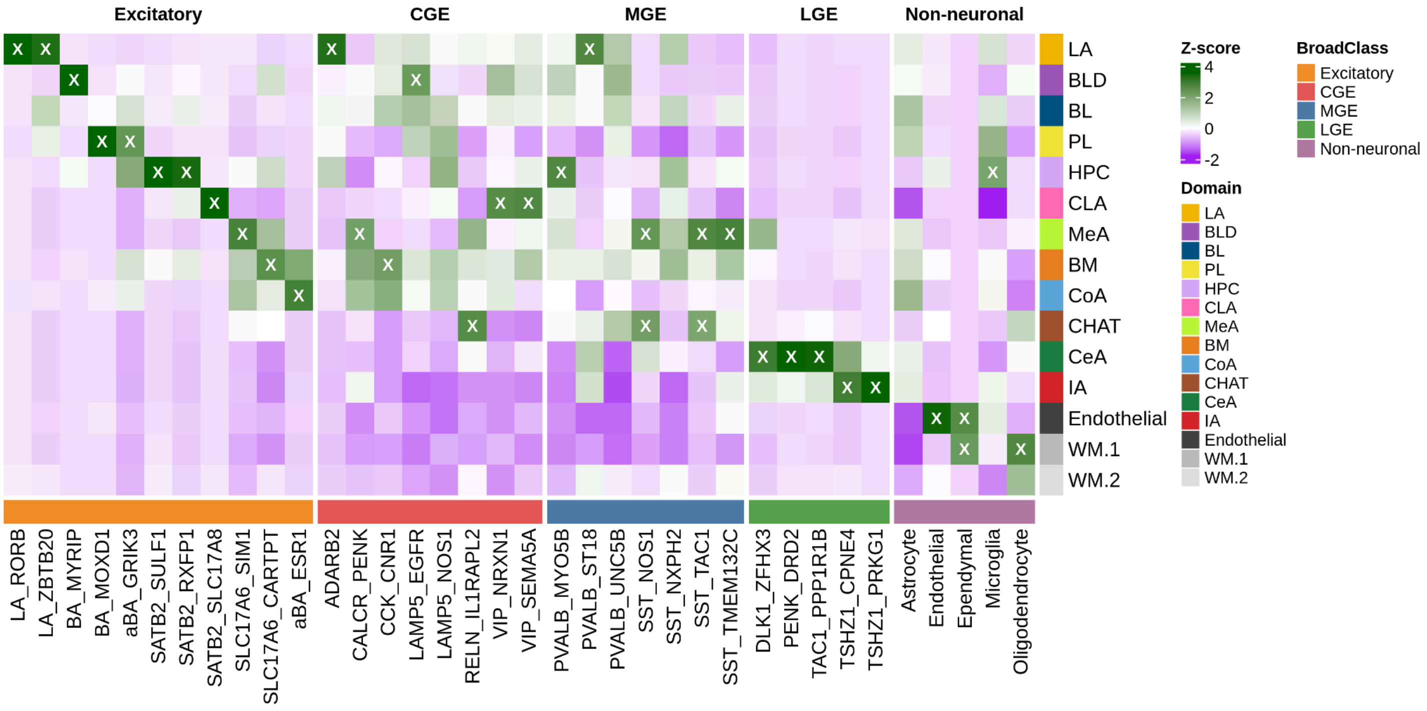


**Supplementary Fig 18. Spatial distribution of Totty et al. macaque cell types across the human amygdala.** RCTD was used to deconvolve Visium spatial transcriptomic data using the Totty et al. macaque snRNA-seq dataset as a reference [19]. The heatmap shows z-scored mean RCTD cell-type weights across spatial domains, with cell types grouped by broad class. Green indicates relative enrichment and purple indicates relative depletion. Crosses denote significant domain–cell-type enrichments.

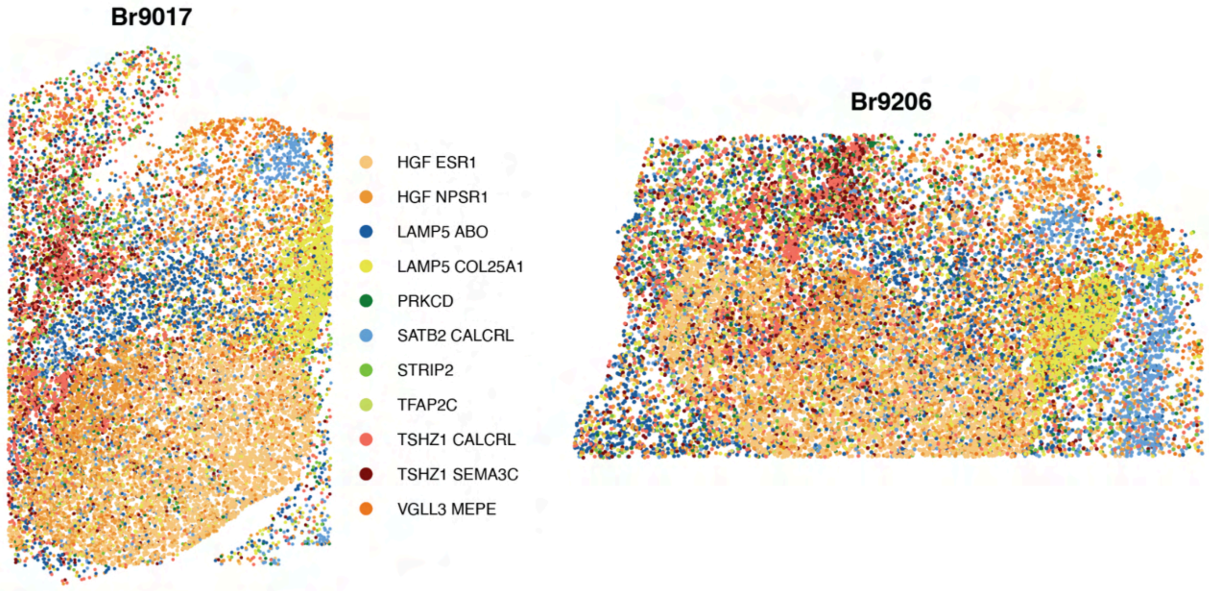


**Supplementary Fig 19. Xenium cell type labels for Br9017 and Br9206.** Spatial maps show the distribution of molecularly defined neuronal cell types across Xenium samples Br9017 and Br9206. Each point represents an individual cell colored according to its assigned cell type from Yu et al.

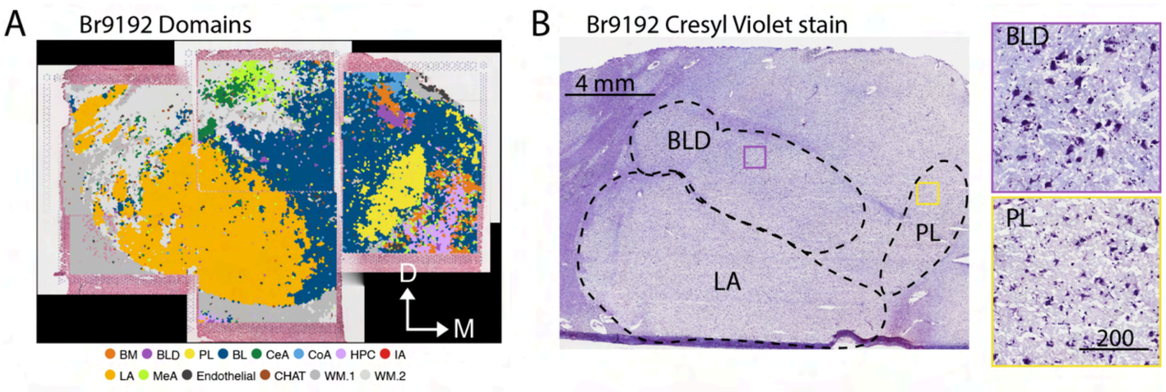


**Supplementary Fig 20. Cresyl Violet stained histology of Br9192 AMY. (A)** Unsupervised spatial domain annotation overlaid on H&E histology, delineating major amygdala nuclei and surrounding structures. Basolateral amygdala (BL), basolateral dorsal amygdala (BLD), basomedial amygdala (BM), central amygdala (CeA), cortical amygdala (CoA), hippocampus (HPC), intercalated amygdalar islands (IA), lateral amygdala (LA), medial amygdala (MeA), paralaminar amygdala (PL), white matter (WM). Tissue directionality is indicated by arrows medial (M) and dorsal (D). **(B)** Br9192 Cresyl Violet stained amygdala at low (left) and high (right) magnifications. Subnuclei are traced by dashed lines. Purple and yellow boxes indicate the locations used for high magnification images of the BLD and PL, respectively. Scale bars are 4 mm for low and 200 μm for high magnification images, respectively.

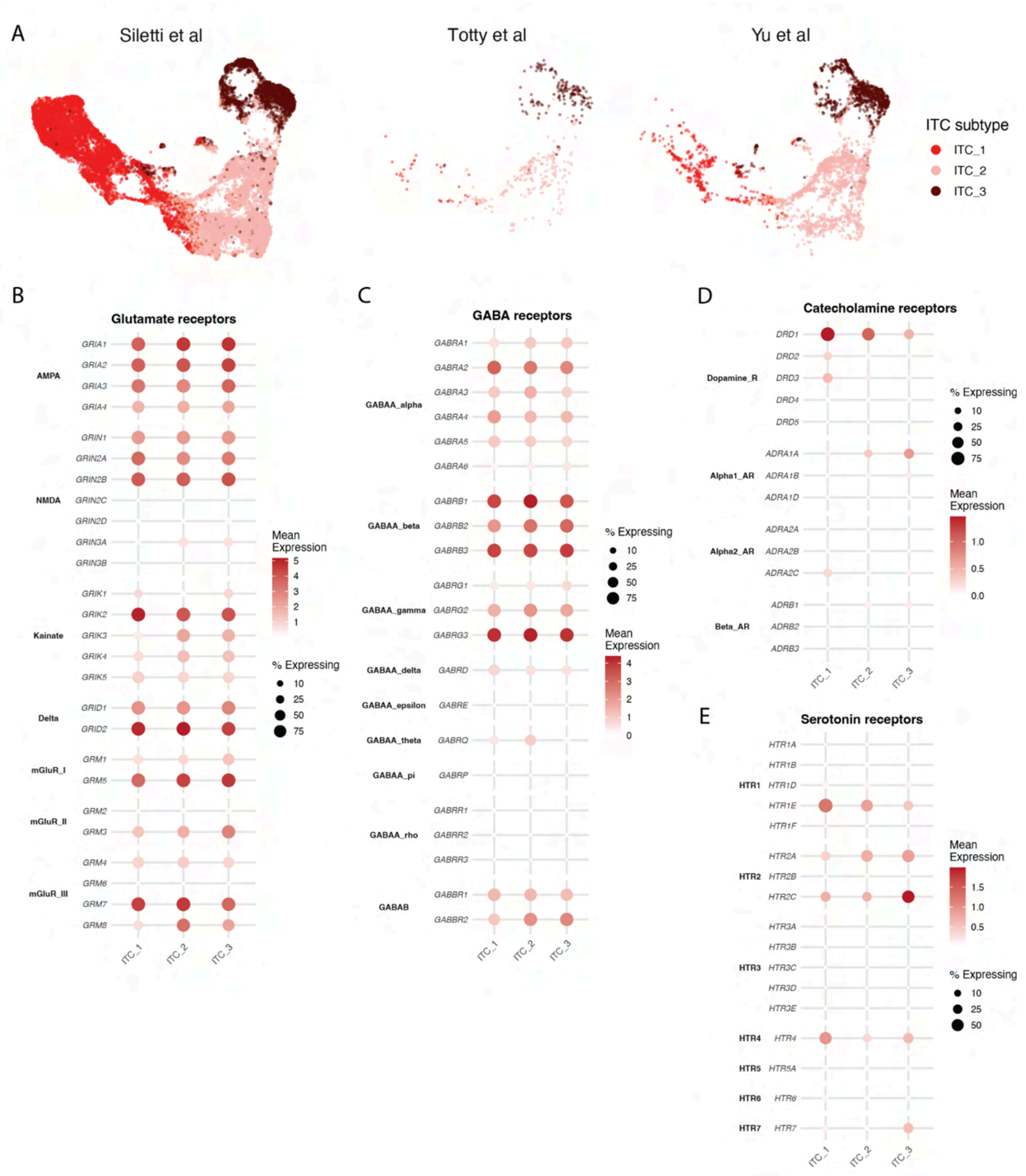


**Supplementary Fig 21. Integrated datasets and neurotransmitter receptor expression across human intercalated cell subtypes. (A)** Integrated UMAP visualization of ITC_1, ITC_2, and ITC_3 across three independent human single-nucleus RNA-seq datasets (Siletti et al., Totty et al., and Yu et al.). Cells are colored by integrated ITC subtype annotation and displayed separately by dataset. **(B–E)** Dot plots showing expression of genes encoding **(B)** glutamate, **(C)** GABA, **(D)** catecholamine, and **(E)** serotonin receptors across ITC subtypes. Dot size represents the percentage of cells expressing each gene and color indicates mean expression. Receptors are grouped by receptor family or subtype.

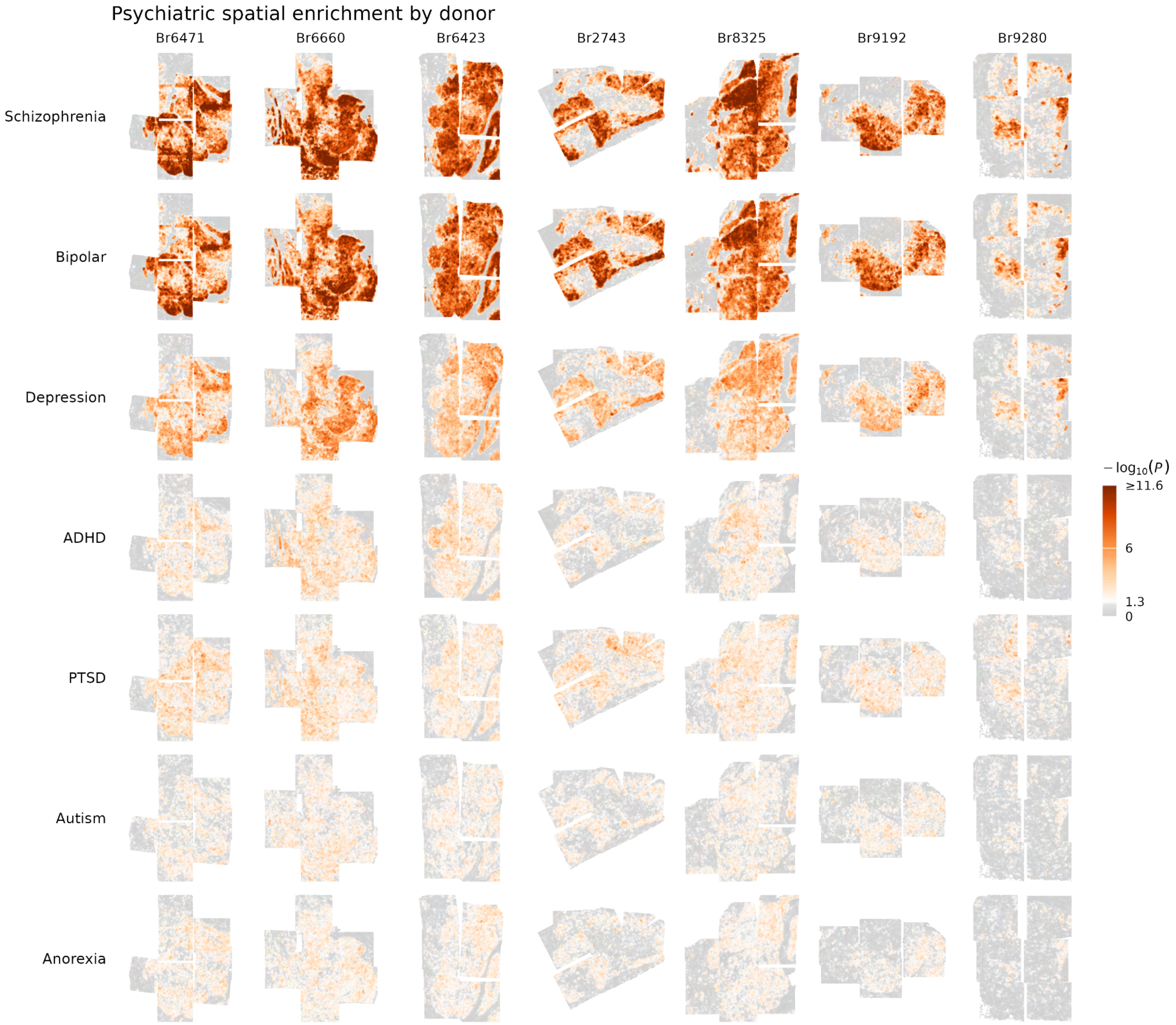


**Supplementary Fig 22. Psychiatric genetic enrichment across the human amygdala.**
Spatial maps show spot-level gsMap enrichment for schizophrenia, bipolar disorder, major depression, ADHD, PTSD, autism, and anorexia across seven human donors. Colors indicate the significance of spatial enrichment (−log10[*P*]), with gray indicating non-significant spots ($P \geq 0.05$). Schizophrenia and bipolar disorder showed the strongest and most widespread enrichment, whereas other psychiatric traits exhibited weaker and more spatially restricted enrichment.

**List of Abbreviations**

ASt - amygdalostriatal transition area
BL - basolateral amygdala
BLA - basolateral amygdalar complex
BLD - basolateral dorsal amygdala
BM - basomedial amygdala
CeA - central amygdala
CeL - lateral subdivision of CeA
CeM - medial subdivision of CeA
CHAT - cholinergic-enriched domain
CLA - claustrum
CoA - cortical amygdala
Ctx - cortex
EC - entorhinal cortex
Endo - endothelial
HPC - hippocampus
IA - intercalated amygdalar islands
ITC - intercalated cells
LA - lateral amygdala
MeA - medial amygdala
OT - optic tract
PL - paralaminar amygdala
PuV - ventral putamen
SUB - subiculum
WM - white matter
